# Cardiomyocyte NLRP3 signaling in right heart failure is sexually dimorphic via estrogen receptor α

**DOI:** 10.64898/2026.02.12.705548

**Authors:** Rafael Sobrano Fais, Erica das Neves Palotta, Katrina W. Kopf, Karina M. Massad, Evandro M. Neto-Neves, Christopher Hoffer, Avram D. Walts, Andrea L. Frump, Neil M. Goldenberg, Sophie Givens, Alice Bourgeois, Chen-Shan C. Woodcock, Irina Petrache, Naomi C. Chesler, Kathleen C. Woulfe, Soni S. Pullamsetti, Olivier Boucherat, Steve Provencher, Brenda M. Ogle, Sébastien Bonnet, Tim Lahm

## Abstract

**Rationale:** RV adaptation in pulmonary hypertension is sexually dimorphic and more preserved in women. NLRP3 inflammasome activation contributes to RV failure (RVF) development. However, regulators and downstream effects of NLRP3 activation in the RV remain unknown.

**Objectives:** We investigated whether NLRP3 inflammasome activation in RVF is sexually dimorphic, whether NLRP3 is active in RV cardiomyocytes (RVCMs) and causes RVCM contractile dysfunction, and whether 17β-estradiol (E2) and its receptor ERα attenuate this process.

**Methods:** We studied RV tissues from PAH patients with RVF, RV tissues and RVCMs isolated from wild-type and ERα loss-of-function mutant rats with RVF, isolated perfused rat hearts, and human induced pluripotent stem cell (hiPSC)-derived cardiomyocytes. NLRP3 activation was assessed via RNA-sequencing, proteomics, immunostaining, and downstream target quantification. RV contractility was assessed via pressure-volume loops, perfused heart studies, and contractility and calcium assessments in isolated RVCMs.

**Measurements and Main Results:** NLRP3 was upregulated in RVCMs during RVF and resulted in altered RVCM calcium handling and RVCM contractile dysfunction. In human RVs, hiPSC-cardiomyocytes and rat RVs, NLRP3 activation and NLRP3-induced RVCM contractile dysfunction were sexually dimorphic and male-biased. Ovariectomy and loss of ERα in females eliminated this sex bias. E2, via ERα, prevented RVCM NLRP3 activation and NLRP3-induced RVCM contractile dysfunction in males and ovariectomized females during both acute and chronic RV pressure overload. ERα directly interacted with NLRP3.

**Conclusions:** NLRP3-driven RVCM contractile dysfunction is male-biased. E2 inhibits NLRP3 through ERα to preserve RVCM contractility. Targeting E2-ERα-NLRP3 signaling may offer novel therapeutic strategies for RVF in low estrogen states.

**Impact:** This is the first study to define a novel estradiol-estrogen receptor α-NLRP3 axis that modulates RV cardiomyocyte function and RV adaptation in pulmonary hypertension. We demonstrate for the first time that NLRP3 activation is therapeutically targetable in low estrogen states via NLRP3 inhibitors or 17β-estradiol. These findings have direct implications for therapeutic strategies aimed at preserving or restoring RV contractile function in pulmonary hypertension, a current area of unmet clinical need.

## Introduction

Right ventricle (RV) adaptation is a major determinant of survival in pulmonary arterial hypertension (PAH) (1). Despite the critical role of RV adaptation in PAH, no RV-targeting therapies exist (2, 3). A major contributor to RV failure (RVF) development in PAH is impaired cardiomyocyte contractility (4). However, the mechanisms regulating RV cardiomyocyte (RVCM) contractility in PAH are incompletely understood. A better understanding may lead to novel, RV-directed therapies for PAH and other types of pulmonary hypertension (PH).

RV adaptation in PAH is sexually dimorphic. Premenopausal women with PH exhibit better RV adaptation and survival than their male counterparts (5–9). 17β-estradiol (E2), the most abundant female sex steroid, has been linked to improved RV function and better RV adaptation (10–13). These effects are mediated at least in part through estrogen receptor (ER) α, which increases abundance of the cardioprotective mediators BMPR2 and apelin in RVCMs (14). However, effects of the E2-ERα-axis likely extend beyond the BMPR2-apelin system. In addition, it is unknown if E2 and ERα directly affect RVCM contractility. A deeper understanding of the mechanisms and targets of E2 and ERα in RVCMs may allow for developing RV-targeting therapies for PH patients with low endogenous E2 states, such as men and postmenopausal women.

Inflammasomes are multiprotein complexes of the innate immune system capable of maturating pro-inflammatory cytokines like IL-1β and IL-18. Nucleotide-Binding Domain-Like Receptor Protein 3 (NLRP3), the most relevant member of the inflammasome family (15, 16), is a crucial mediator of vascular (17), lung (18), and heart (19, 20) diseases. For example, NLRP3 is involved in the genesis and aggravation of cardiovascular diseases, promoting vascular (17) and left ventricle (LV) remodeling (19, 20). In LV failure, NLRP3 negatively regulates contractile signaling (21, 22), and impairs cardiomyocyte contractility (23). However, these findings cannot be extrapolated to the RV, as the LV and RV are embryologically, structurally, and physiologically distinct (24–26). NLRP3 activation specifically in recruited macrophages promotes the development of RVF (27). However, the role and regulation of NLRP3 signaling specifically in RVCMs remains unknown.

We previously identified that E2’s protective effects in RVF include anti-inflammatory effects (13). Since E2 attenuates NLRP3 activation in the inflamed airway (23, 28), we hypothesized that E2, via ERα, attenuates NLRP3 inflammasome activation in RVCMs and alleviates NLRP3-induced RVCM contractile dysfunction. We report for the first time that RVCM NLRP3 activation in rodent and human RVF is sexually dimorphic, that NLRP3 activation results in RVCM contractile dysfunction in males but not females, that ERα prevents NLRP3-induced in female RVCMs, and that E2, in an ERα-dependent manner, attenuates RVCM NLRP3 activation and NLRP3-induced RVCM contractile impairment in RVF. Some of the results have been published in abstract form (29, 30).

## Methods

Studies were performed in RV tissues from PAH patients with RVF, in RV tissues and RVCMs isolated from wild-type and ERα loss-of-function mutant (ERα^mut^) rats (14) with RVF, in isolated perfused rat hearts, and in human induced pluripotent stem cell (hiPSC)-derived cardiomyocytes (31–33). All experiments and analyses were performed in accordance with recent recommendations (2, 34), including randomization and blinding at the time of intervention, endpoint collection, and analysis. *Full methods are provided in the online data supplement*.

## Results

### NLRP3 activation in human RVF is sexually dimorphic and male-biased

We first assessed expression of RV NLRP3 inflammasome components in male and female PAH patients with RVF. We interrogated previously published transcriptome and proteome data from (35) and (36) (*Fig. 1A; hemodynamics in Table E2*). We found that RNA and protein for NLRP3 activators, NLRP3 inflammasome components, and NLRP3 downstream mediators indeed were upregulated in human PAH-RVs (*Fig. 1B&C*). Despite similar severity of RVF in both genders, this was sexually dimorphic and more upergulated in male RVs *(Fig. 1D&E*). Abundance of NLRP3 downstream mediators caspase-1, IL-1β, and IL-18 was negatively correlated with cardiac index (*Fig. 1F-H*), suggesting worse RV function with more NLRP3 activation.

**Fig 1:**
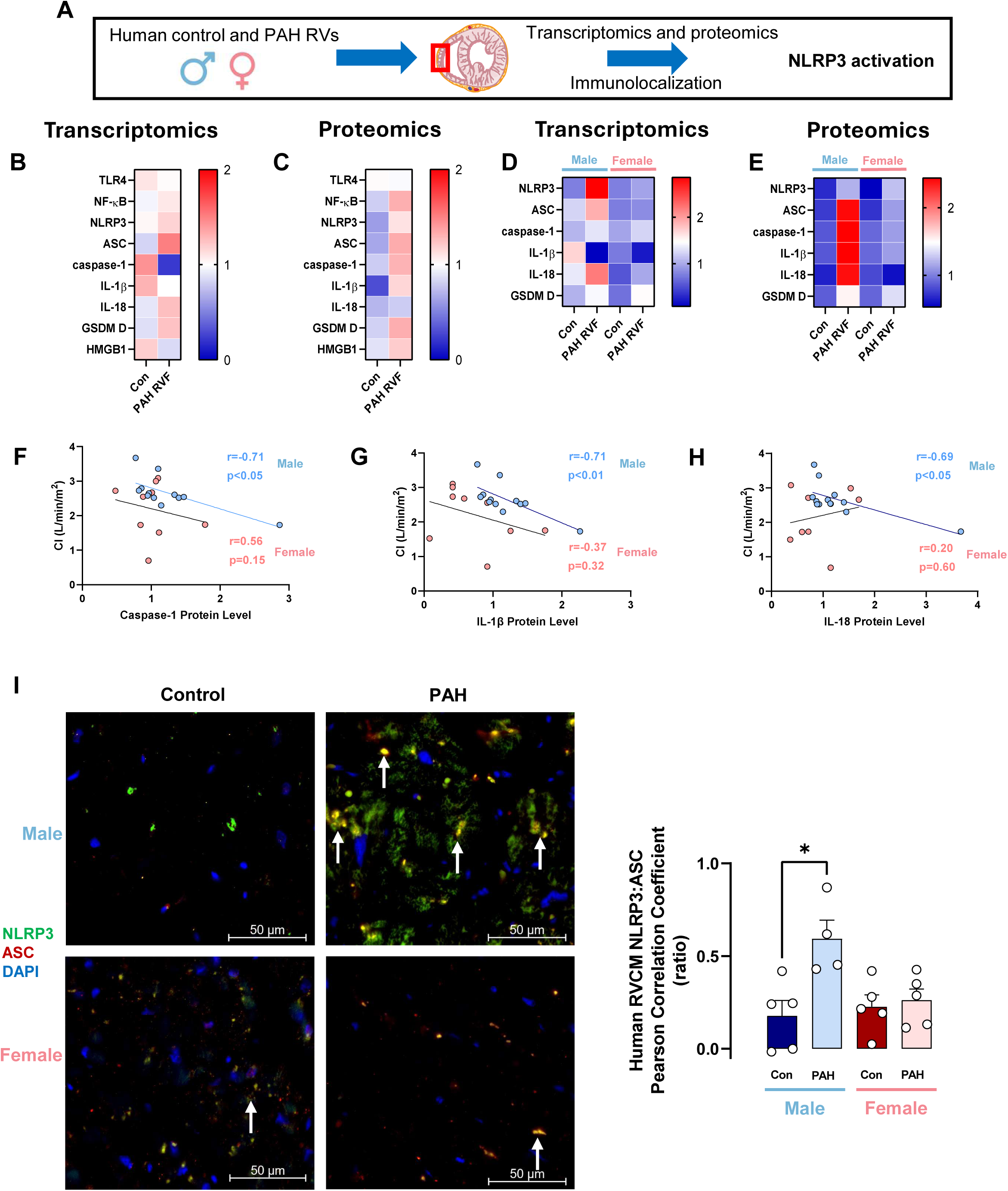
NLRP3 in RVs from pulmonary artery hypertension (PAH) patients with RV failure (RVF) is activated in a sexually dimorphic manner. (**A**) Experimental design. (**B, C**) Heat maps demonstrating RV expression of NLRP3 inflammasome pathway components in human control samples and samples from PAH patients with RVF by RNA-Seq (B) and Proteomics (C). N=13 for control; n=6 for RVF. Data were analyzed from publicly available data in references (35) and (36). (**D, E**) RNA and proteomic profiles of RV NLPR3 signaling components expression in control and RVF subjects stratified by sex. N=8 for male control, n=1 for male RVF, n=5 for female control, and n=5 for female RVF. (F-H) Correlations of protein abundance of NLRP3 inflammasome downstream mediators caspase-1, IL-1β, and IL-18 with cardiac index (CI) in male and female human PAH RVF samples. (**I**) Representative images (left) and quantification (right) of colocalization of NLRP3 and its binding partner ASC in RVs from male or female human control samples and samples from PAH patients with RVF. Magnification is 20x. Human RV samples were stained for NLRP3 (green), ASC (red) and DAPI (blue). Co-localization was analyzed in the whole RV sample; Magnification is 40x for representative images. Arrows indicate NLRP3-ASC co-localization (yellow). Co-localization was quantified by determining Pearson correlation coefficient between NLRP3 and ASC. *p<0.05 by two-way ANOVA followed by Holm-Šídák post-test. Each data point represents one RV sample from one subject (means ± SEM). Abbreviations used in heatmaps: TLR4 = toll-like receptor 4, NF-κB = nuclear factor-κB, NLRP3 = NOD-like family pyrin domain containing 3, ASC = apoptosis-associated speck-like protein containing a CARD, IL-1β = interleukin-1β, IL-18 = interleukin-18, GSDM D = gasdermin D, HMGB1 = high-mobility group box 1.

We corroborated these data by quantifying protein abundance and co-localization of NLRP3 and its binding partner ASC (*hemodynamics in Table E3).* We detected a 4-fold increase in ASC expression in male RVs (*Fig. E1*). NLRP3-ASC co-localization was increased 3-fold in male RVs, whereas no such increase was noted in females *(Fig. 1I*). These data suggest a male sex bias in NLRP3 activation in human RVF.

### NLRP3 inflammasome activation in pressure-overloaded RVs is more pronounced in male and ovariectomized female rats and is attenuated by E2 treatment

We next assessed NLRP3 activation in rodent models of RVF. In rats with monocrotaline (MCT)-induced RVF, NLRP3 was differently activated in male and female RVs, with increased NLRP3 and ASC abundance (*Fig. E2*), increased NLRP3-ASC co-localization (*Fig. 2A&B)*, and increased activation of downstream targets caspase-1 and IL-1β *(Fig. 2C&D*) in male RVs only. This occurred despite similar increases in RVSP in both sexes and more RV hypertrophy in females (*Table E4).* However, RV NLRP3 activation was noted in female MCT rats after ovariectomy (OVX; *Fig. 2B&C*). NLRP3 abundance, NLRP3-ASC co-localization, and downstream mediator activation in male RVs was attenuated by treatment with E2 to a similar degree than by treatment with NLRP3 inhibitor MCC950 (*Fig. 2A-D & E2*), demonstrating that E2 attenuates NLRP3 activation as potently as an established NLRP3 inhibitor. E2 also attenuated NLRP3 activation in female OVX RVs (*Fig. 2B&C*).

**Fig 2:**
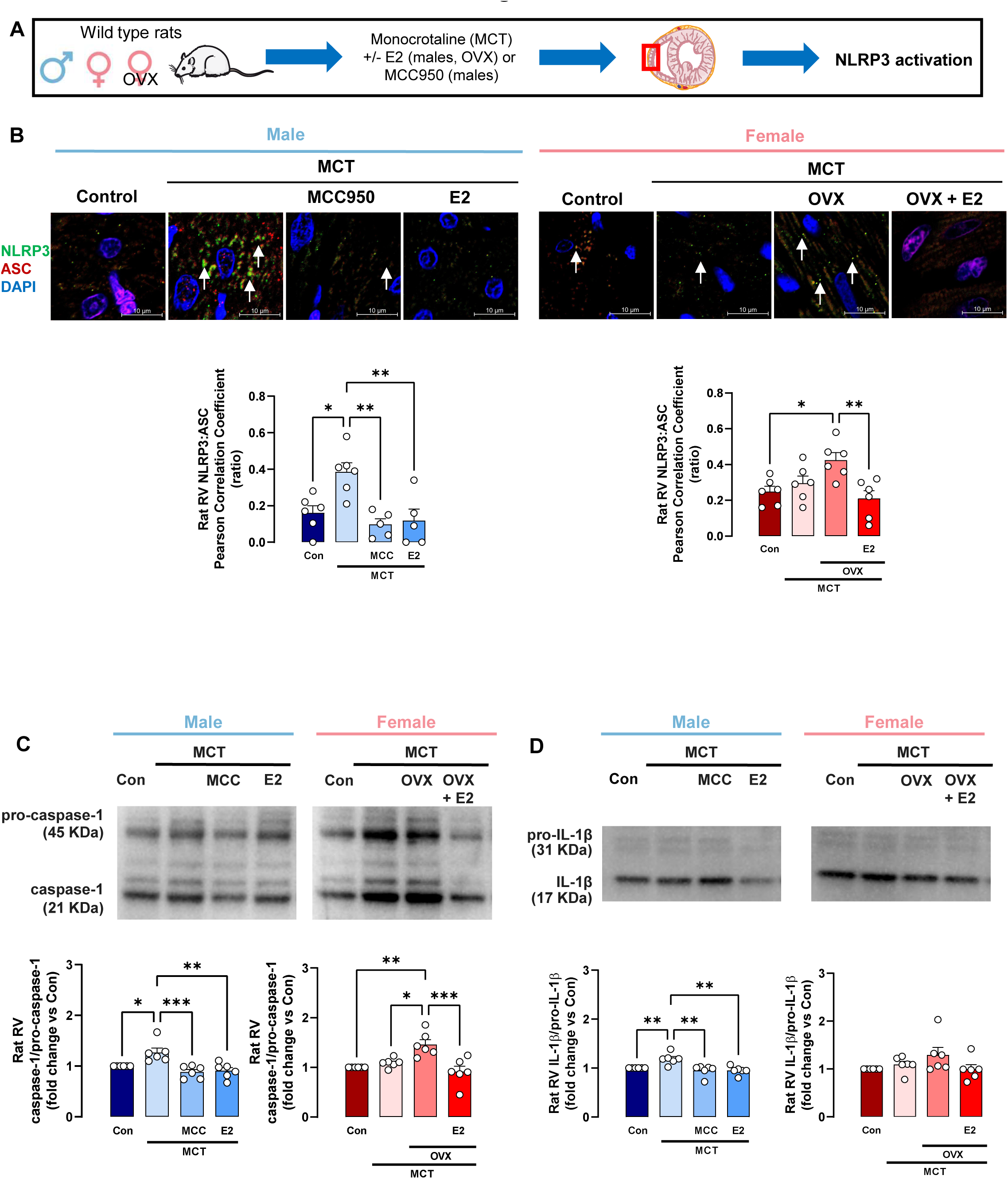
RV NLRP3 inflammasome activation in rats with monocrotaline (MCT)-induced PH exhibits a male bias that is due to inhibitory effects of 17β-estradiol (E2). (**A**) Experimental design. RVs were isolated from male, intact or ovariectomized (OVX) female control or MCT rats. Subgroups of MCT-PH males were treated with E2 (75 μg/kg/day by subcutaneous [s.c.] pellets) or NLRP3 inhibitor MCC950 (10 mg/kg by s.c. injection) for 4 weeks (starting at MCT injection). Subgroups of OVX MCT-PH females were treated with E2 (75 μg/kg/day by s.c. pellets) for 4 weeks (starting at MCT injection). (**B**) Representative immunofluorescence images and quantification of fluorescence intensity of RV NLRP3-ASC colocalization. RVs were stained for NLRP3 (green), ASC (red), and DAPI (blue). Co-localization images were analyzed in ten different fields, and values were then averaged. Images were obtained at 100x magnification. Representative images were digitally zoomed 5x. NLRP3 and ASC co-localization (yellow; arrows) were quantified by determining Pearson correlation coefficient. (**C, D**) Representative Western blot images and densitometric quantification of caspase-1/pro-caspase-1 ratio (**C**) and IL-1β/pro-IL-1β ratio (**D**). *p<0.05, **p<0.01, ***p<0.001, ns = not significant by one-way ANOVA with Holm-Šídák post-test. Each data point = one rat (means ± SEM).

To study sexual dimorphisms in NLRP3 signaling independently of potential changes in the pulmonary vasculature, we assessed NLRP3 signaling in RVs from male and female pulmonary artery banding (PAB) rats (*Fig. E3A*). Despite similar hemodynamic alterations (*Table E5*), NLRP3 activation again demonstrated a male bias, with increases in ASC abundance, NLRP3-ASC co-localization, and caspase-1 and IL-1β activation in male RVs only *(Fig. E3&4*).

Of note, no evidence of NLRP3 activation was observed in LVs of male PH-rats (*Fig. E5*). Together, these data demonstrate a male sex bias in RV NLRP3 activation in rats with RVF and suggest that E2 attenuates NLRP3 activation in RVF rats as potently as NLRP3 inhibitor.

### NLRP3 activation results in contractile dysfunction in male RVs that is inhibited by E2 treatment

We next investigated whether NLRP3 activation results in RV contractile dysfunction. We isolated hearts from male and female rats, and measured RV isovolumic contractile function after perfusion with the potent and established NLRP3 activators lipopolysaccharide (LPS) and ATP (37, 38) in presence or absence of NLRP3 inhibitor or E2 (*Fig. 3A*). LPS+ATP resulted in NLRP3 activation and RV contractile dysfunction, which was prevented by NLRP3 inhibition or E2 (*Fig. 3B&C*). Conversely, RVs from female rats did not exhibit NLRP3 activation and RV contractile dysfunction with LPS+ATP (*Fig. 3C*), and administration of NLRP3 inhibitor or E2 did not affect contractile function in female RVs.

**Fig 3:**
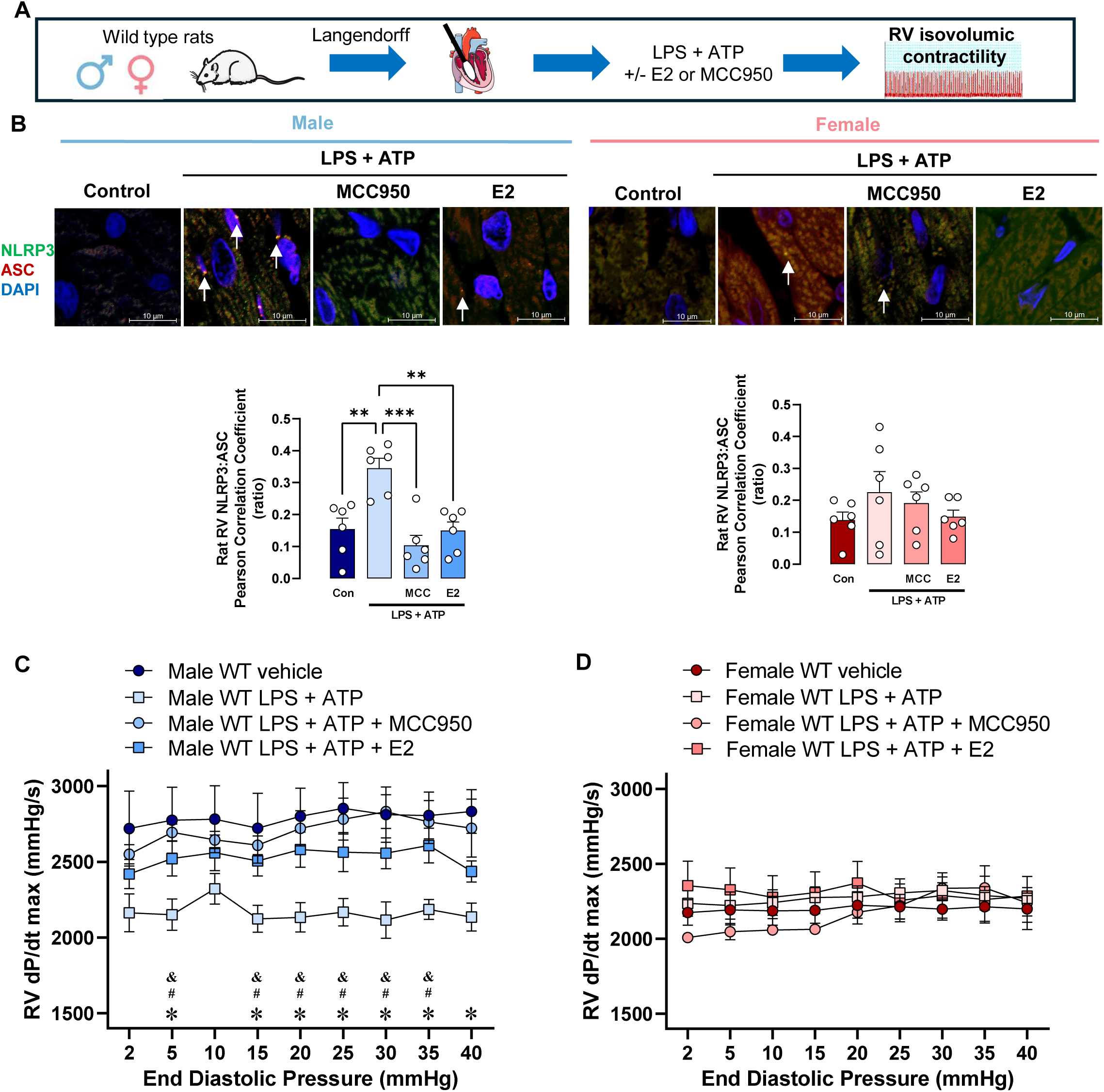
NLRP3 activation decreases RV isovolumic contractility in a sex-specific and male-biased manner, and this decrease is prevented by E2 in male RVs. (**A**) Experimental design. (**B**) Representative immunofluorescence images and quantification of fluorescence intensity of NLRP3-ASC co-localization in RVs from male or female control rat hearts perfused with lipopolysaccharide (LPS; 1 μg/mL, 130 min prior to start of dP/dt max measurement and continued throughout measurement) and ATP (2 mM, 10 min prior to start of dP/dt max measurement and continued throughout measurement) or vehicle, in presence or absence of MCC950 (1 μM, 30 min prior to LPS and continued throughout LPS exposure) or E2 (1 nM, 30 min prior to LPS and continued throughout LPS exposure). Rat RVs were stained for NLRP3 (green), ASC (red), and DAPI (blue). Images were obtained at 100x magnification. Co-localization images were analyzed in ten different fields per animal, and averaged values for each animal are shown. NLRP3 and ASC co-localization (yellow; indicated by arrows) was quantified by determining Pearson correlation coefficient. Representative images were digitally zoomed 5x. (**C, D**) Maximum rate of rise in RV pressure during systole (dP/dt max) in male (**C**) and female (**D**) rats. dP/dt max was measured at increasing amounts of RV end-diastolic pressure (2 to 40 mmHg). Data are shown as mean ± SEM (N=5-13/group). *p<0.05 for male LPS + ATP vs. male vehicle group, ^#^p<0,05 for male LPS + ATP vs. male LPS + ATP + MCC950, ^&^p<0,05 for male LPS + ATP vs. male LPS + ATP + E2 by one-way ANOVA with Holm-Šídák post-test.

These data indicate that NLRP3 activation results in RV contractile dysfunction in a male-biased manner, and that this process is attenuated by E2 treatment.

### RVCM NLRP3 inflammasome activation and NLRP3-induced RVCM contractile dysfunction in MCT-PH rats are sexually dimorphic, male-biased, and attenuated by E2

We next interrogated whether NLRP3 activation occurs specifically in RVCMs. We isolated RVCMs from male and intact as well as OVX female MCT rats (*Fig. 4A*). Indeed, NLRP3 was differently activated in male and female RVCMs, with increases in NLRP3 and ASC abundance (*Fig. E6*), NLRP3-ASC co-localization (*Fig. 4B*), and caspase-1 and IL-1β activation (*Fig. 4C&D*) in male MCT-RVCMs only. Functionally, NLRP3 activation was associated with decreased sarcomere shortening (a measure of RVCM contractility) and reduced transient Ca^2+^ (a measurement of free cytosolic Ca^2+^) in male but not female MCT-RVCMs (*Fig. 4E&F*). OVX, on the other hand, rendered female MCT-RVCMs susceptible to NLRP3 activation and contractile dysfunction (*Fig. 4B-F*).

**Fig 4:**
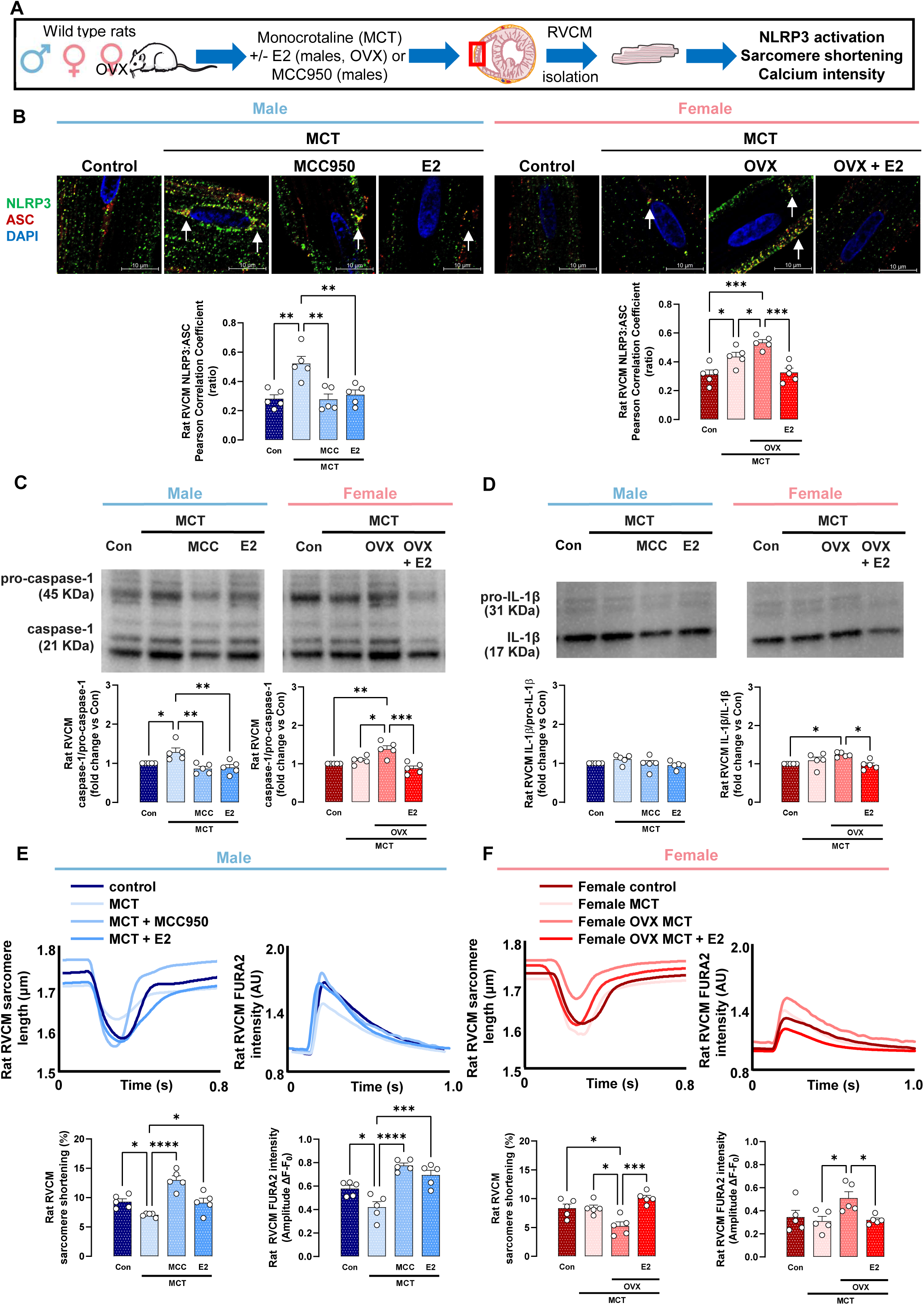
RV cardiomyocyte (RVCM) NLRP3 inflammasome activation and NLRP3-induced RVCM contractile dysfunction in MCT-PH rats are sexually dimorphic, male-biased, and attenuated by 17β-estradiol (E2). (**A**) Experimental design. (**B**) Representative immunofluorescence images and quantification of fluorescence intensity of NLRP3-ASC co-localization in RVCMs from male, intact or ovariectomized (OVX) female control or MCT rats. Subgroups of MCT-PH males were treated with NLRP3 inhibitor MCC950 (10 mg/kg by subcutaneous [s.c.] injection) or E2 (75 μg/kg/day by s.c. pellets) for 4 weeks (starting at MCT injection). Subgroups of MCT-PH females were ovariectomized and treated with E2 (75 μg/kg/day by s.c. pellets) for 4 weeks (starting at MCT injection). Rat RVCMs were stained for NLRP3 (green), ASC (red), and DAPI (blue). Images were obtained at 100x magnification. Co-localization images were analyzed in ten different RVCMs per animal, and averaged values for each animal are shown. NLRP3 and ASC co-localization (yellow; indicated by arrows) was quantified by determining Pearson correlation coefficient. Representative RVCM images were digitally zoomed 5x. (**C, D**) Representative Western blot images and densitometric quantification of caspase-1/pro-caspase-1 ratio (**C**) and IL-1β/pro-IL-1β ratio (**D**) in RVCMs from male or female control or MCT rats as well as male MCT-PH rats treated with MCC950 or E2 as well as female OVX MCT-PH rats treated with E2. (**E, F**) Representative tracings and quantification of rat RVCM sarcomere shortening and FURA2 intensity. Tracings of RVCM sarcomere shortening demonstrate changes in absolute length after electrical stimulation; quantification shows RVCM shortening in % (determined as [relaxed length - contracted length/relaxed length] x 100) in ten different RVCMs per dish. Tracings of FURA2 intensity demonstrate arbitrary units (AU); quantification shows the ratio of change in fluorescence (ΔF) to baseline fluorescence (F_0_; ΔF/F_0_). *p<0.05, **p<0.01, ***p<0.001, ****p<0.0001 by one-way ANOVA with Holm-Šídák post-test. Each data point = RVCMs from one rat (means ± SEM).

In vivo treatment of male and OVX female rats with E2 prevented MCT-induced NLRP3 pathway activation as well as decreases in RVCM sarcomere shortening and transient Ca^2+^ (*Fig. 4B-F*). Of note, E2 treatment replicated protective effects of MCC950. These data indicate 1) that MCT-induced NLRP3 activation impairs RVCM contractility and Ca^2+^ signaling in male and OVX female but not intact female rats, 2) that E2 attenuates NLRP3-induced RVCM dysfunction, and 3) that E2’s effect size is similar to that of an established NLRP3 inhibitor.

### LPS+ATP-induced NLRP3 activation in rat RVCMs is sexually dimorphic and prevented by E2

To determine the role of E2-induced NLRP3 inhibition in RVCMs independently of MCT, we isolated RVCMs from healthy rats and activated the NLRP3 inflammasome with LPS+ATP in vitro *(Fig. 5A)*. LPS+ATP indeed activated the NLRP3 inflammasome and increased caspase-1 and IL-1β activation in male, but not female, RVCMs (*Fig. 5B-D*). Functionally, LPS+ATP-induced NLRP3 activation was associated with a decrease in sarcomere shortening and transient Ca^2+^ in male but not female RVCMs (*Fig. 5E-F*).

**Fig 5:**
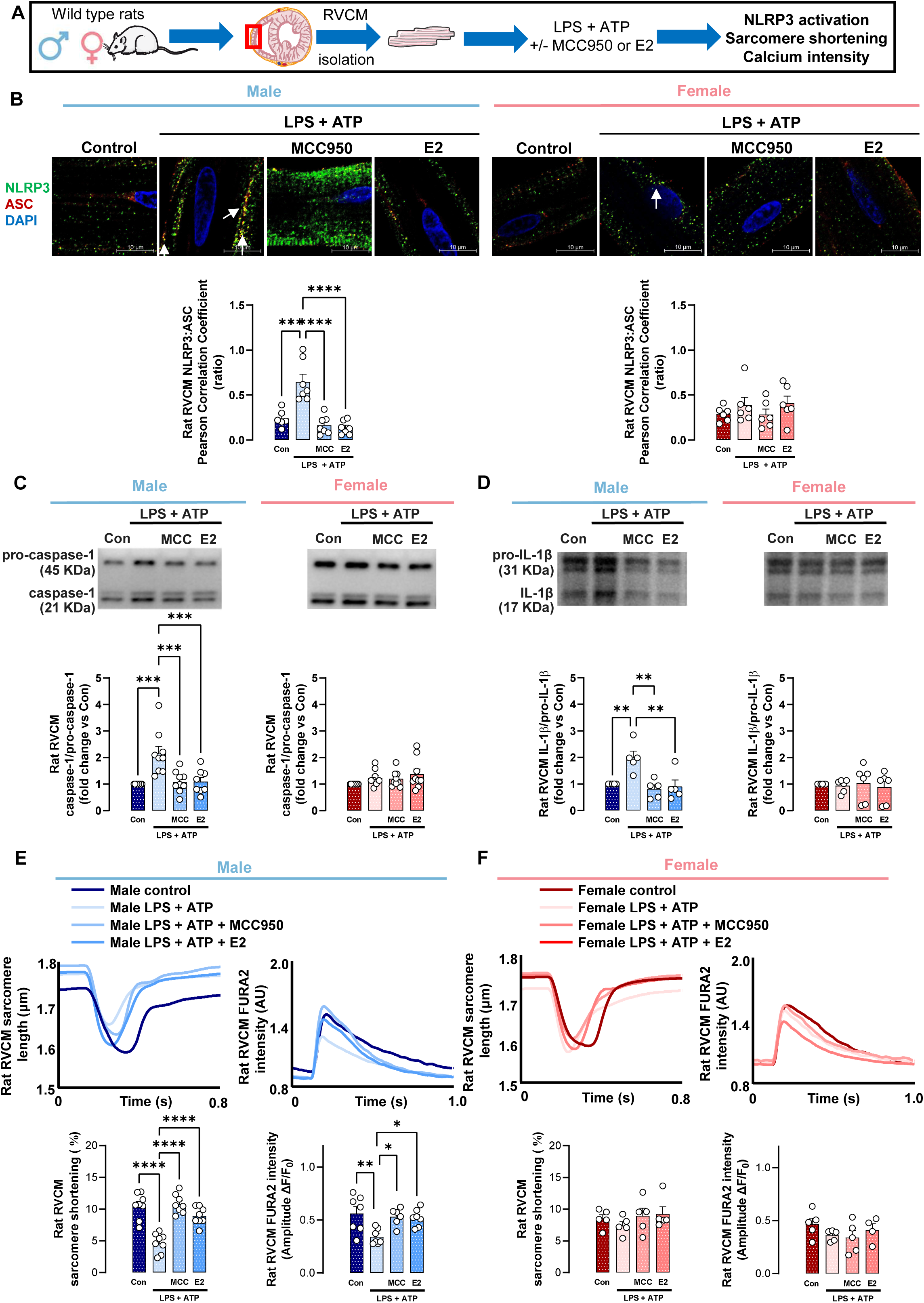
Lipopolysaccharide (LPS)- and adenosine triphosphate (ATP)-induced NLRP3 activation in RVCMs is sexually dimorphic and prevented by E2 in male RVCMs. (**A**) Experimental design. (**B**) Representative immunofluorescence images and quantification of fluorescence intensity of NLRP3-ASC co-localization in RVCMs from male or female control rats treated in vitro with LPS+ATP in absence or presence of NLRP3 inhibitor MCC950 or E2. RVCMs were treated with LPS (1 μg/mL) for 4 hours and ATP (2 mM) for 10 min prior to cell collection. MCC950 (1 μM) was given 30 min prior to LPS and continued throughout LPS and ATP exposure. E2 (1 nM) was given for 24 hours prior to LPS and continued throughout LPS and ATP exposure. Rat RVCMs were stained for NLRP3 (green), ASC (red), and DAPI (blue). Images were obtained at 100x magnification. Co-localization images were analyzed in ten different RVCMs per animal, and averaged values for each animal are shown. NLRP3 and ASC co-localization (yellow; indicated by arrows) was quantified by determining Pearson correlation coefficient. Representative RVCM images were digitally zoomed 5x. (**C, D**) Representative Western blot images and densitometric quantification of caspase-1/pro-caspase-1 ratio (**C**) and IL-1β/pro-IL-1β ratio (**D**) in RVCMs from male or female control rats treated in vitro with LPS+ATP in absence or presence of MCC950 or E2 as outlined above. (**E, F**) Representative tracings and quantification of rat RVCM sarcomere shortening and FURA2 intensity in RVCMs from male or female control rats treated in vitro with LPS+ATP in absence or presence of MCC950 or E2 as outlined above. Tracings of RVCM sarcomere shortening demonstrate changes in absolute length after electrical stimulation; quantification shows RVCM shortening in % (determined as [relaxed length - contracted length/relaxed length] x 100) in ten RVCMs per dish. Tracings of FURA2 intensity demonstrate arbitrary units (AU). Quantification shows the ratio of change in fluorescence (ΔF) to baseline fluorescence (F_0_; ΔF/F_0_). *p<0.05, **p<0.01, ***p<0.001, ****p<0.0001; ns = not significant by one-way ANOVA with Holm-Šídák post-test. Each data point = RVCMs from one rat (means ± SEM).

Co-treatment of male RVCMs with E2 attenuated NLRP3 activation and NLRP3-induced RVCM contractile and Ca^2+^ signaling dysfunction similar to MCC950 (*Fig. 5B-F*). On the other hand, E2 did not exert any effects in female RVCMs (where there was no NLRP3 activation; *Fig. 5B-F*). Data for NLRP3 and ASC as well as E2 dose responses are shown in *Fig. E7,8&9*.

These data indicate that LPS+ATP-induced NLRP3 activation and NLRP3-mediated contractile dysfunction in healthy rat RVCMs are sexually dimorphic, male-biased, and prevented by E2.

### Endothelin-1 (ET1) -induced NLRP3 activation in rat RVCMs is sexually dimorphic and prevented by E2

While LPS and ATP are potent NLRP3 activators, they are not involved in RVF pathogenesis. We therefore sought to determine whether a known mediator of PAH and RVF development results in RVCM NLRP3 activation and contractile dysfunction. We treated healthy male or female rat RVCMs with ET-1. As with LPS+ATP, NLRP3 activation was sexually dimorphic, male-biased, and prevented by E2 in male RVCMs (*Fig. E10&11*).

### NLRP3 activation in human iPSC-CMs exhibits a male bias and is attenuated by E2

To determine whether sexual dimorphisms in NLRP3 activation extend to human CMs, we stimulated hiPSC-CMs from three male and three female lines with ET-1 (*Fig. 6A*). We found that NLRP3 and ASC abundance as well as NLRP3-ASC colocalization increased 1.7- to 2-fold in male cells, whereas no increase was observed in female cells (*Fig. 6B & E12*). E2 attenuated this increase in male clones, similar to NLRP3 inhibitor (*Fig. 6B & E12*). This data suggests that NLRP3 activation is also sexually dimorphic, male-biased, and preventable by E2 in human CMs.

**Fig 6:**
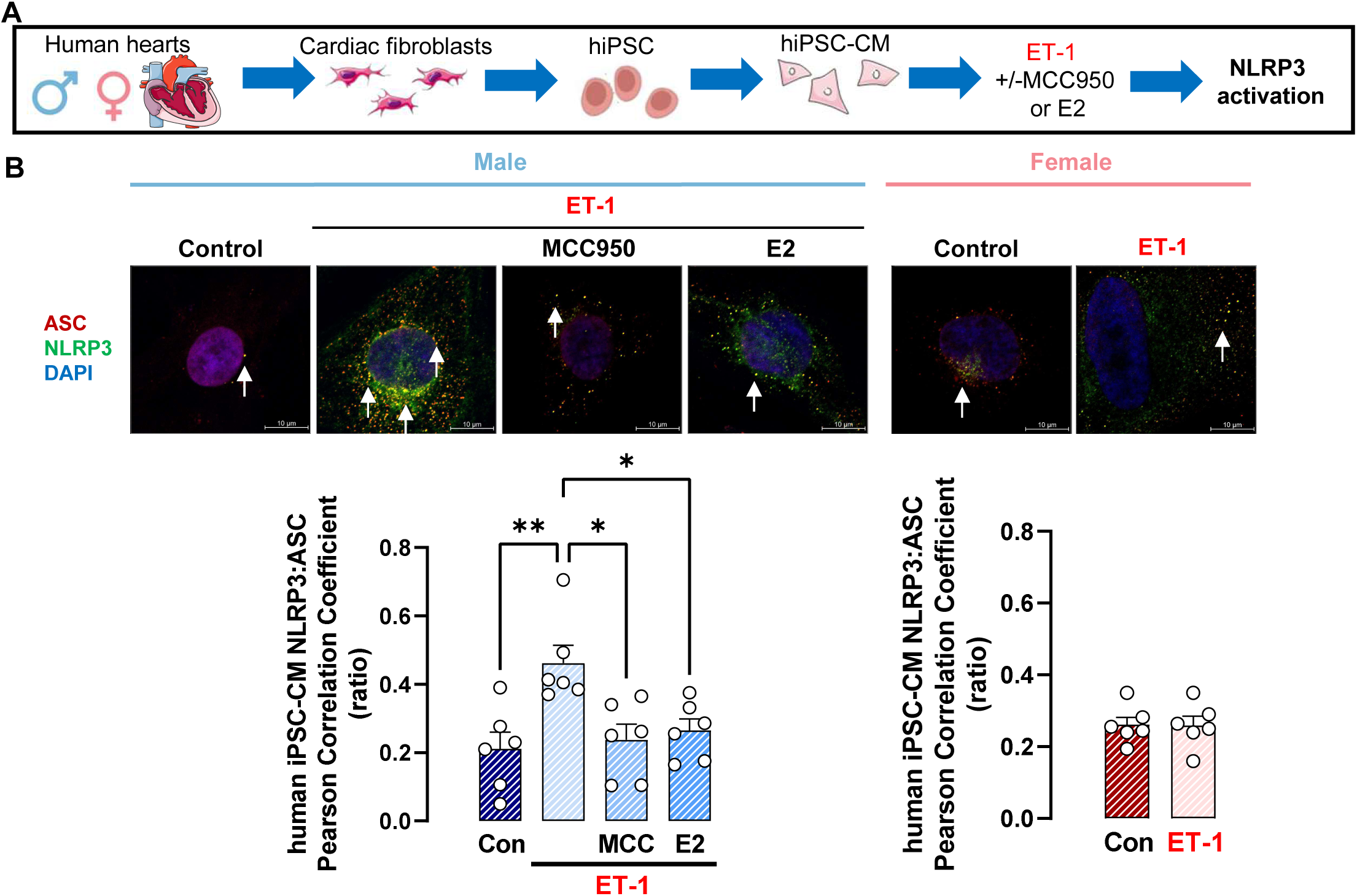
Endothelin 1-induced NLRP3 activation in human induced pluripotent stem cell derived cardiac myocytes (hiPSC-CMs) is sexually dimorphic, male biased, and prevented by E2 in male cells. (**A**) Experimental design. (**B**) Representative immunofluorescence images and quantification of fluorescence intensity of NLRP3-ASC co-localization in hiPSC-CMs treated with endothelin-1 (ET-1; 1 nM) given 4 hours prior to cell collection +/- MCC950 (1 μM; administered 30 min prior to ET-1 and continued throughout ET-1 exposure) or E2 (1 nM, administered 24 hours prior to ET-1 and continued throughout ET-1 exposure). Cells were stained for NLRP3 (green), ASC (red), and DAPI (blue). Co-localization was quantified in ten different hiPSC-CMs per patient line by determining Pearson correlation between NLRP3 and ASC, and values were then averaged. Each data point = one human iPSC-CM clone obtained from 3 male and 3 female iPSC-CM cell line (means ± SEM). *p<0.05, **p<0.01 by one-way ANOVA with Holm-Šídák post-test (male cells) or Student t-test (female cells).

### Lack of functioning ERα renders female rats susceptible to NLRP3-induced RV contractile dysfunction, eliminates sex biases, and prevents E2 from attenuating NLRP3-mediated RV contractile impairment

To investigate the potential cross-talk between ERα and NLRP3-induced RV contractile dysfunction noted in the isolated perfused heart model with LPS+ATP perfusion (*Fig. 3*), we repeated this experiment using previously described male and female ERα loss-of-function mutant (ERα^mut^) rats (14). We measured RV isovolumic contractile function in isolated hearts after perfusion with LPS+ATP (*Fig. 7A*). As previously seen in the WT (*Fig. 3B*), in RVs from male ERα^mut^ rats, LPS+ATP induced NLRP3 activation and contractile dysfunction (which was rescued after NLRP3 inhibition; *Fig. 7B&C*). On the other hand, and in contrast to the female WT *(Fig. 3C)*, female ERα^mut^ rat RVs also exhibited NLRP3 activation and RV contractile dysfunction with LPS+ATP *(Fig. 7D*). In both male as well as female ERα^mut^ rat RVs (and in contrast to male WT RVs), E2 was unable to prevent NLRP3 activation and NLRP3-induced contractile dysfunction *(Fig. 7B-D)*. These data indicate that ERα is necessary in female RVs to prevent NLRP3-induced contractile impairments, and for E2 to attenuate NLRP3-induced RV contractile dysfunction in both sexes.

**Fig 7:**
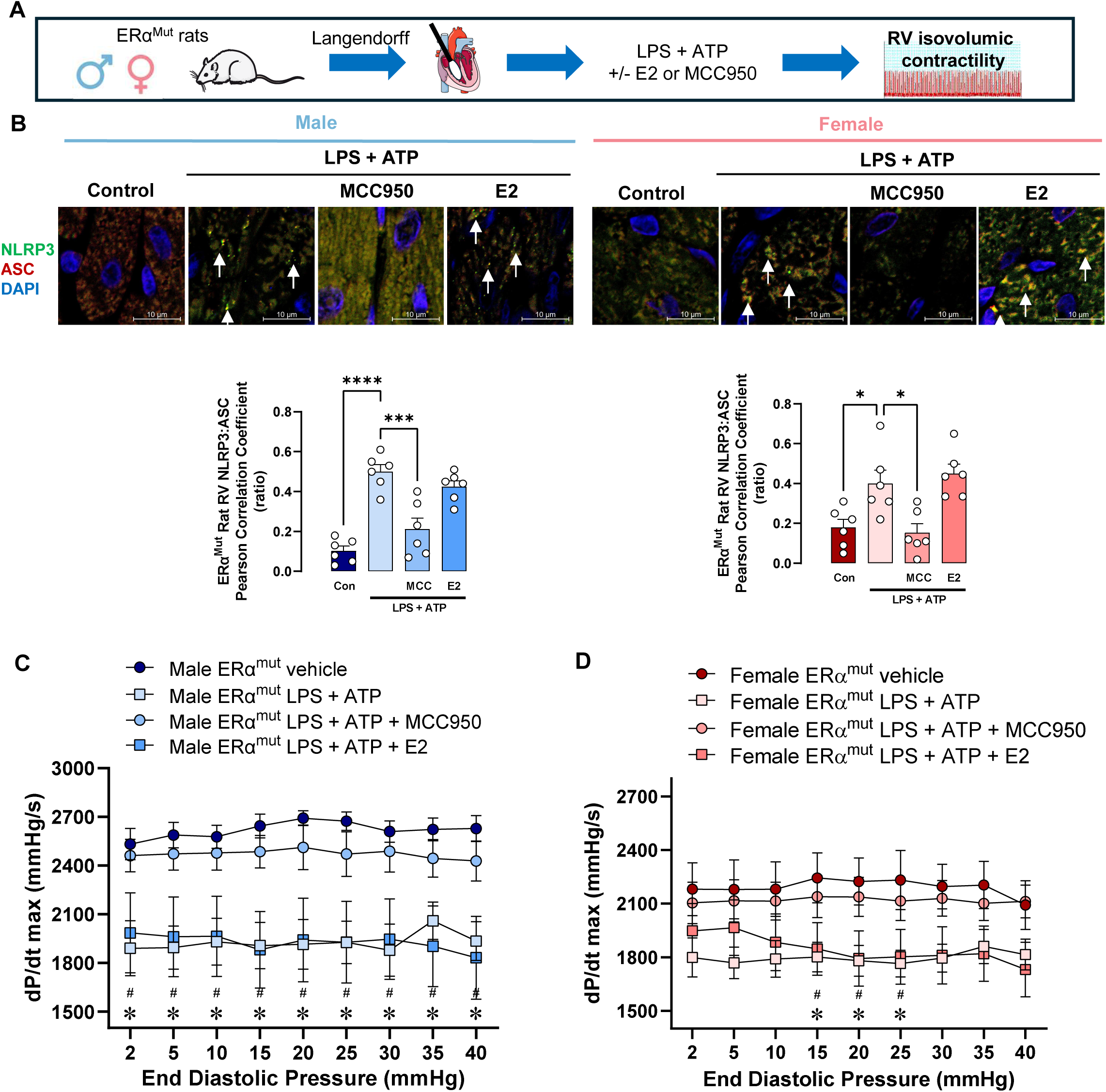
Loss of ERα prevents E2 from attenuating RV contractile dysfunction in male RVs and renders female RVs susceptible to contractile dysfunction after NLRP3 activation. (**A**) Experimental design. (**B**) Representative immunofluorescence images and quantification of fluorescence intensity of NLRP3-ASC co-localization in RVs from male or female control rat hearts perfused with LPS (1 μg/mL, 130 min prior to start of dP/dt max measurement and continued throughout measurement) + ATP (2 mM, 10 min prior to start of dP/dt max measurement and continued throughout measurement), in the presence or absence of MCC950 (1 μM, 30 min prior to LPS and continued throughout LPS exposure) or E2 (1 nM, 30 min prior to LPS and continued throughout LPS exposure). Rat RVs were stained for NLRP3 (green), ASC (red), and DAPI (blue). Images were obtained at 100x magnification. Co-localization images were analyzed in ten different fields per animal, and averaged values for each animal are shown. NLRP3 and ASC co-localization (yellow; indicated by arrows) was quantified by determining Pearson correlation coefficient. Representative RVCM images were digitally zoomed 5x. Maximum rate of rise in RV pressure during systole (dP/dt max) in male (**C**) and female (**D**) ERα loss-of-function mutant (ERα^mut^) rats in an isolated perfused heart model. dP/dt max was measured at increasing amounts of RV end-diastolic pressure. Data are shown as mean ± SEM (N=5-8/group). *p<0.05 for male LPS + ATP vs. male vehicle group, ^#^p<0,05 for male LPS + ATP vs male LPS + ATP + MCC950 by one-way ANOVA with Holm-Šídák post-test.

### Lack of functioning ERα eliminates sexual dimorphisms in RVCM NLRP3 activation in vivo and prevents E2 from attenuating NLRP3 activation

To investigate the role of ERα in alleviating MCT-induced NLRP3 activation in females and in mediating protective effects of E2, we employed ERα^mut^ rats, treated them with MCT ± E2 in vivo, and isolated their RVCMs (*Fig. 8A*). As previously seen in the WT (*Fig. 4*), in male MCT-PH ERα^mut^ rats, we observed robust increases in RVCM NLRP3 and ASC abundance, NLRP3-ASC co-localization, and caspase-1 and IL-1β activation, as well as decreases in RVCM contractility and transient Ca^2+^ *(Fig. 8B-F & E13*). However, in contrast to the female WT, female MCT-PH ERα^mut^ rats also exhibited NLRP3 activation, RVCM contractile dysfunction, and Ca^2+^ dysfunction *(Fig. 8B-F & E13*). In both male as well as female MCT-PH ERα^mut^ rats, E2 was unable to prevent RVCM NLRP3 activation, Ca^2+^ signaling impairments and contractile dysfunction. These data indicate that ERα is necessary in female RVCMs to prevent NLRP3 activation and NLRP3-induced contractile dysfunction, and for E2 to attenuate RVCM NLRP3 activation and NLRP3-induced contractile dysfunction in male MCT-PH rats.

**Fig 8:**
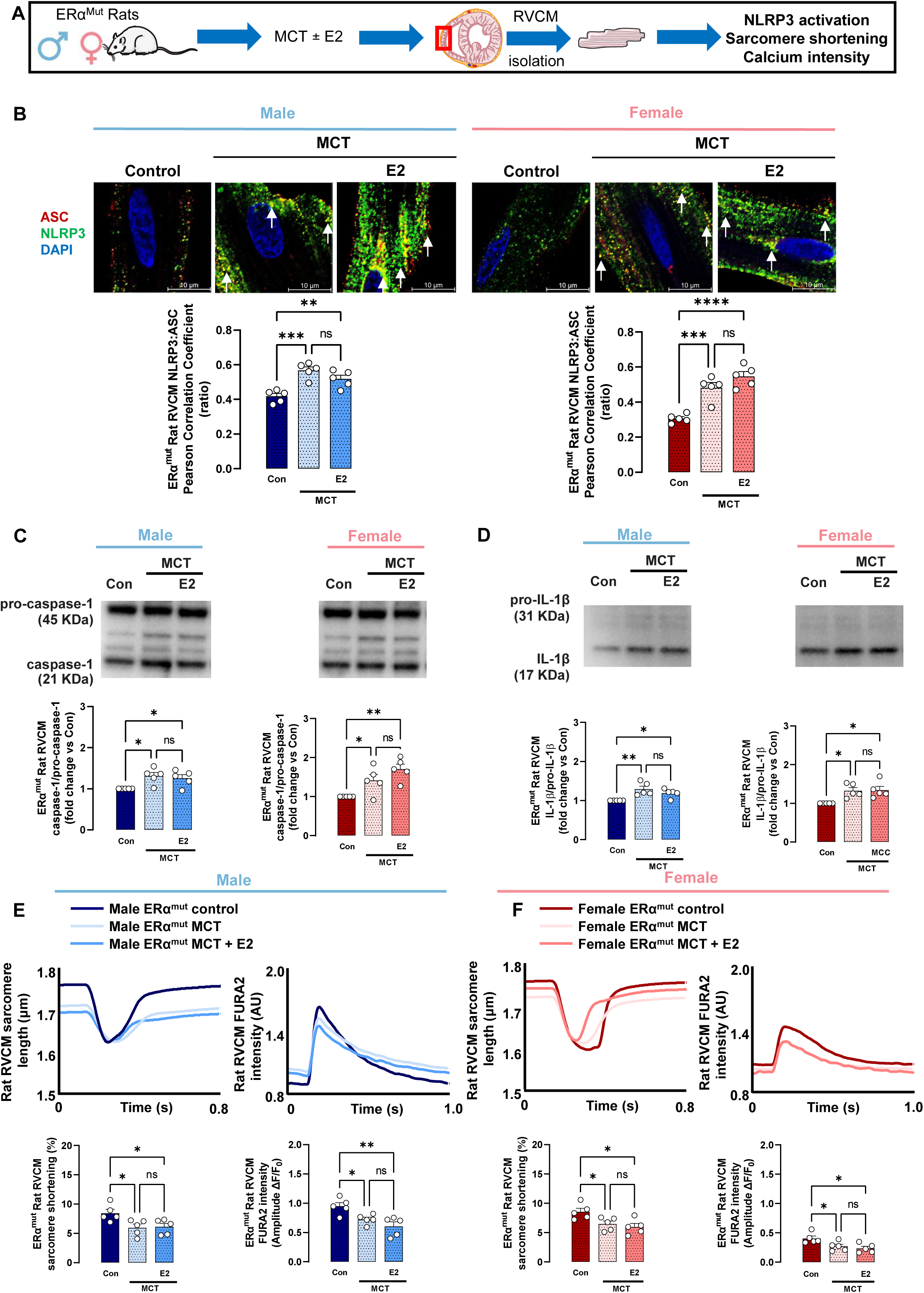
Lack of functioning estrogen receptor α (ERα) eliminates sexual dimorphisms in NLRP3 activation in RVCMs from MCT-PH rats and prevents E2 from attenuating NLRP3 activation in these cells. (**A**) Experimental design. (**B**) Representative immunofluorescence images and quantification of NLRP3-ASC colocalization in RVCMs from male or female ERα loss-of-function mutant (ERα^mut^) rats with MCT-PH. Subgroups of ERα^mut^ MCT-PH rats were treated with E2 (75 μg/kg/day via s.c. pellets) for 4 weeks (starting at MCT injection). Images were obtained at 100x magnification. RVCMs were stained for NLRP3 (green), ASC (red), and DAPI (blue). Images were obtained at 100x magnification. Co-localization images were analyzed in ten different RVCMs per animal, and averaged values for each animal are shown. NLRP3 and ASC co-localization (yellow; arrows) was quantified by determining Pearson correlation coefficient. Representative images were digitally zoomed 5x. (**C, D**) Representative Western blot images and densitometric quantification of caspase-1/pro-caspase-1 ratio (**C**) and IL-1β/pro-IL-1β ratio (**D**) in RVCMs from male or female ERα^mut^ MCT-PH rats with or without E2 treatment in vivo. (**E, F**) Representative tracings and quantification of rat RVCM sarcomere shortening and FURA2 intensity in RVCMs from male or female ERα^mut^ rats with MCT-PH with or without E2. Tracings of RVCM sarcomere shortening demonstrate changes in absolute length after electrical stimulation; quantification shows RVCM shortening in % (determined as [relaxed length - contracted length/relaxed length] x 100) in 10 RVCM per dish. Tracings of FURA2 intensity demonstrate arbitrary units (AU); quantification shows the ratio of change in fluorescence (ΔF) to baseline fluorescence (F_0_; ΔF/F_0_). *p<0.05, **p<0.01, ***p<0.001, ****p<0.0001; ns = not significant by one-way ANOVA with Holm-Šídák post-test. Each data point = RVCMs from one rat (means ± SEM).

### Lack of functioning ERα eliminates sexual dimorphisms in rat RVCM NLRP3 activation and prevents protective effects of E2

We next sought to study crosstalk between ERα and NLRP3 signaling in RVCMs independently of potential changes in the pulmonary vasculature in MCT-PH rats. We isolated RVCMs from ERα^mut^ rat RVs and treated them with LPS + ATP in vitro (*Fig. 9A*). We found that in males, similar to WT *(Fig. 5)*, LPS + ATP resulted in NLRP3 activation, Ca^2+^ signaling impairments, and contractile dysfunction (*Fig. 9B-F & E14*). In contrast to female WT RVCMs (*Fig. 5*), RVCMs from female ERα^mut^ rats also exhibited such changes (*Fig. 9B-F & E14*). In both male as well as female ERα^mut^ rat RVCMs, E2 was incapable of preventing LPA + ATP-induced NLRP3 activation, Ca^2+^ signaling impairments, and contractile dysfunction (*Fig. 9B-F & E14*). These data corroborate findings from *Fig. 8* and indicate a protective role of ERα in attenuating NLRP3 activation in RVCMs.

**Fig 9:**
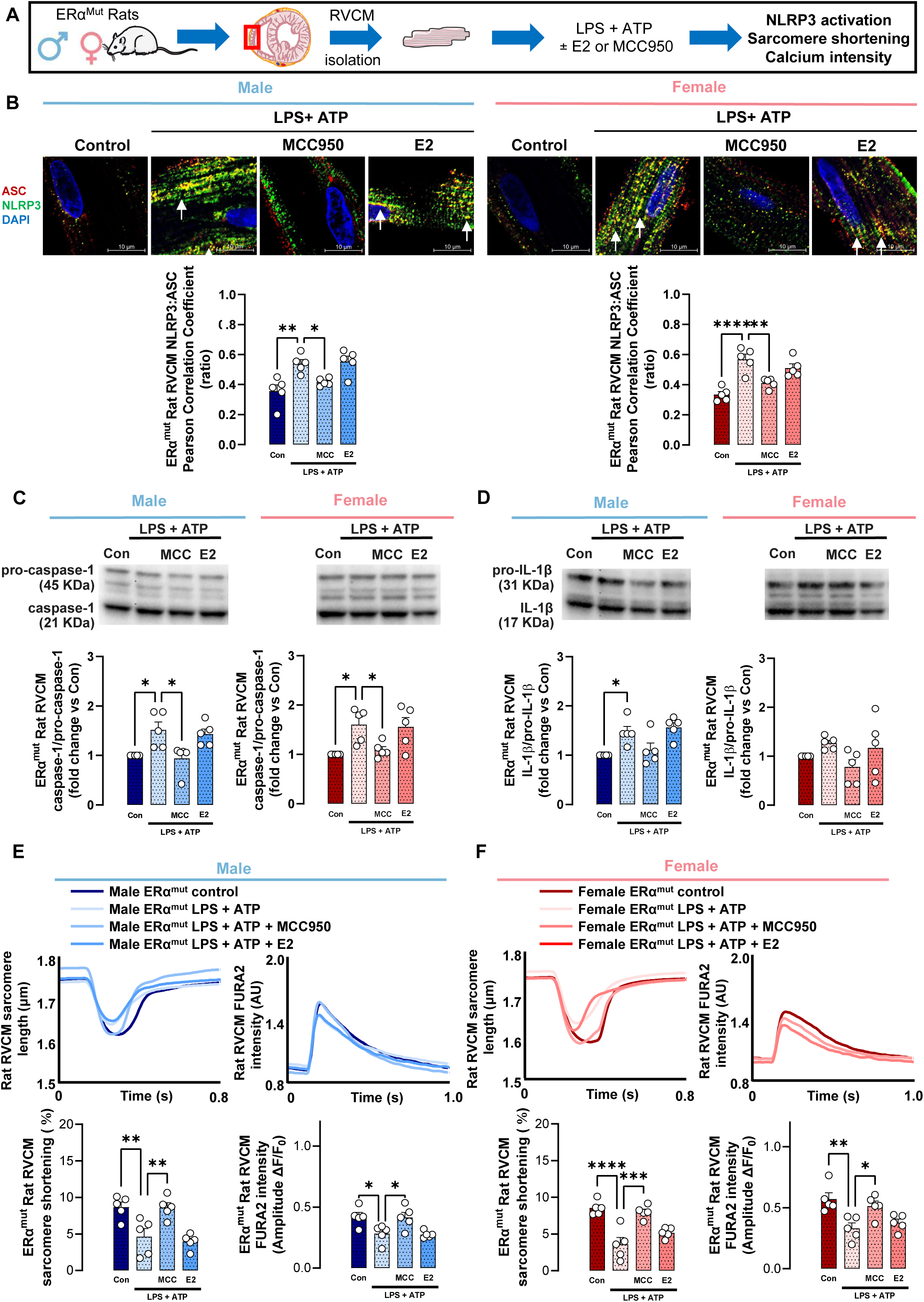
Lack of functioning estrogen receptor α (ERα) eliminates sexual dimorphisms in LPS + ATP-induced NLRP3 activation in rat RVCMs and prevents E2 from attenuating NLRP3 activation in these cells. (**A**) Experimental design. (**B**) Representative immunofluorescence images and quantification of NLRP3-ASC co-localization in RVCMs from male or female ERα loss-of-function mutant (ERα^mut^) rats after treatment with LPS and ATP in vitro. LPS (1 μg/mL) was given 4 hours prior to cell collection. ATP (2 mM) was given 10 min prior to cell collection. MCC950 (1 μM) or E2 (1 nM) were given 30 min or 24 hours prior LPS and ATP, respectively, and continued throughout the exposure. RVCMs were stained for NLRP3 (green), ASC (red), and DAPI (blue). Images were obtained at 100x magnification. Co-localization images were analyzed in ten different RVCMs per animal, and averaged values for each animal are shown. NLRP3 and ASC co-localization (yellow; arrows) was quantified by determining Pearson correlation coefficient. Representative images were digitally zoomed 5x. (**C, D**) Representative Western blot images and densitometric quantification of caspase-1/pro-caspase-1 ratio (**C**) and IL-1β/pro-IL-1β ratio (**D**) in RVCMs from male or female ERα^mut^ rats stimulated with LPS+ATP ± E2 or MCC950 in vitro as outlined in (B). (**E, F**) Representative tracings and quantification of rat RVCM sarcomere shortening and FURA2 intensity in RVCMs from male or female ERα^mut^ rats treated with LPS+ATP ± E2 or MCC950 in vitro as outlined in (B). Tracings of RVCM sarcomere shortening demonstrate changes in absolute length after electrical stimulation; quantification shows RVCM shortening in % (determined as [relaxed length - contracted length/relaxed length] x 100) in the RVCM per dish. Tracings of FURA2 intensity demonstrate arbitrary units (AU); quantification shows the ratio of change in fluorescence (ΔF) to baseline fluorescence (F_0_; ΔF/F_0_). *p<0.05, **p<0.01, ***p<0.001, ****p<0.0001; ns = not significant by one-way ANOVA with Holm-Šídák post-test. Each data point = RVCMs from one rat (means ± SEM).

### ERα directly interacts with NLRP3

To determine whether ERα’s inhibitory effects on NLRP3 activation are mediated by a direct interaction, we assessed for NLRP3 and ERα co-localization in human RVF tissue and hiPSC-CMs stimulated with ET-1 *(Fig. 10A&G, E15&16*). We found that NLRP3 and ERα indeed co-localized *(Fig. 10B&C, H&I*). A direct interaction between NLRP3 and ERα was confirmed by co-IP in RVs from male and female MCT-PH and PAB rats (*Fig. E17*). Interestingly, NLRP3 and ERα co-localization was increased 3- to 4-fold in male RVF-RVs and ET1-stimulated male hiPSC-CMs (vs control RVs and unstimulated hiPSC-CMs, respectively; *Fig. 10B&C, H&I*). On the other hand, no increase was noted in female RVF-RVs or ET1-stimulated hiPSC-CMs. This was accompanied by decreased ERα nuclear intensity in male (but not female) RVF-RVs and ET1-stimulated hiPSC-CMs *(Fig. 10D&J*). Higher NLRP3-ERα co-localization correlated with higher NLRP3 activation and lower nuclear ERα abundance *(Fig. 10E&F, K&L*), suggesting that the degree of NLRP3-ERα co-localization is associated with the degree of NLRP3 activation, and that increased NLRP3-ERα co-localization is associated with reduced nuclear ERα localization (and a potential decrease in ERα nuclear activity). Similar findings were noted in RVs from male and female MCT-PH and PAB rats (*Fig. E17*) and in male and female MCT-PH rat RVCMs (*Fig. E18*).

**Fig 10:**
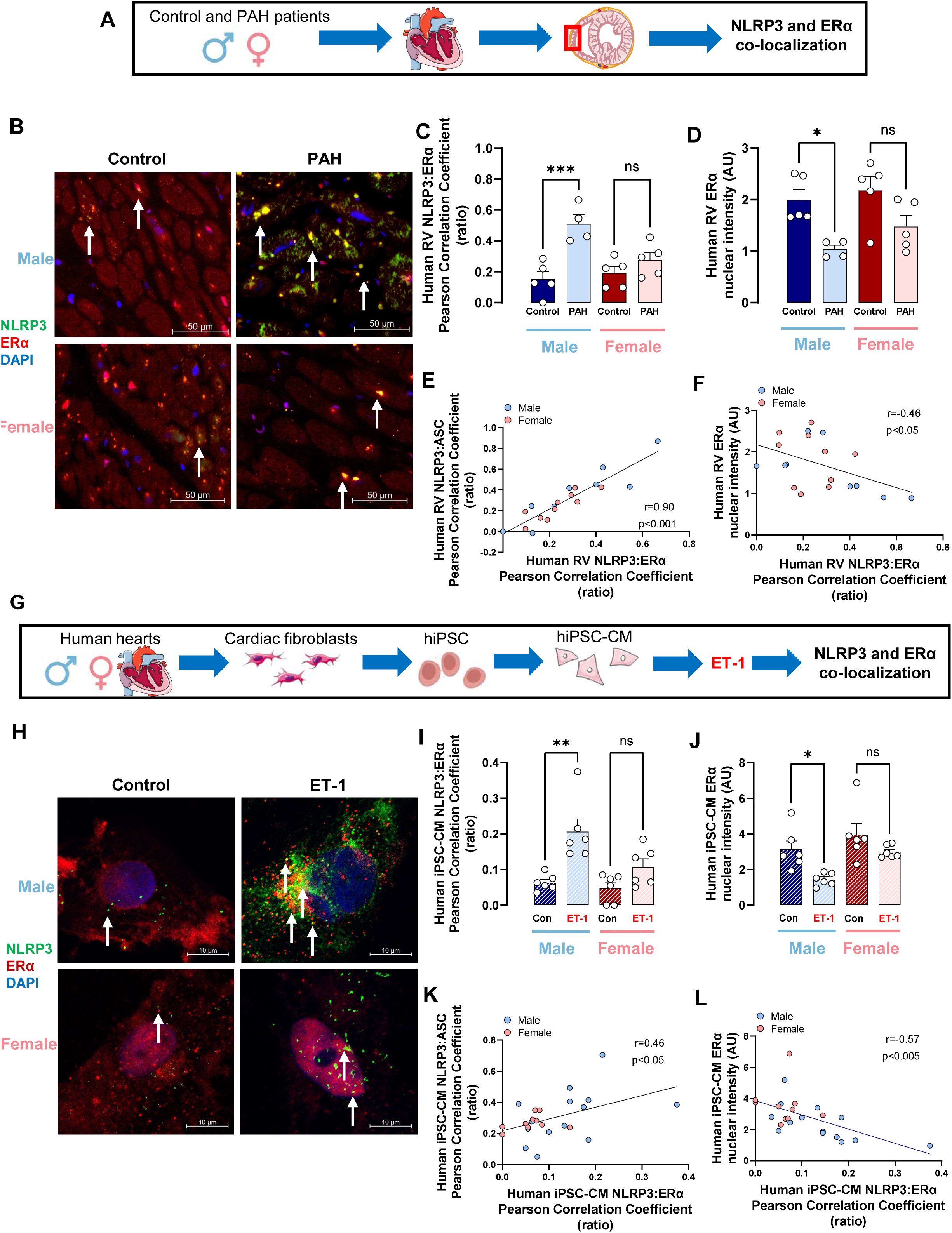
ERα co-localizes with NLRP3 in RVs from PAH patients with RV failure and in human induced pluripotent stem cell-derived cardiac myocytes (hiPSC-CMs). (**A**) Experimental design for PAH patient studies. (**B, C**) ERα-NLRP3 co-localization in RVs from PAH patients with RV failure. (**B**) Representative images and (**C**) quantification of co-localization of NLRP3 (green) with ERα (red). Magnification is 20x. Co-localization was analyzed in the whole RV sample; representative images are 40x. Arrows indicate ERα-NLRP3 co-localization (yellow). Co-localization was quantified by determining Pearson correlation coefficient between NLRP3 and ERα. Note sexual dimorphism and male bias with more pronounced co-localization in male PAH RVs. (**D**) ERα nuclear intensity in all experimental groups. (**E, F**) demonstrate correlations between ERα-NLRP3 co-localization intensity and (**E**) NLRP3 activation (NLRP3-ASC co-localization) and (**F**) nuclear ERα abundance. Note decrease in nuclear ERα abundance with increase in ERα-NLRP3 co-localization. (**G**) Experimental design for hiPSC-CM studies. (**H, I**) ERα-NLRP3 co-localization in hiPSC-CMs with and without endothelin 1 (ET-1; 1 nM, 4 hours prior to cell collection). (**H**) Representative images and (**I**) quantification of co-localization of NLRP3 (green) with ERα (red). Magnification is 100x. Co-localization (arrows) was quantified by determining Pearson correlation coefficient between NLRP3 and ERα. Note sexual dimorphism and male bias with more pronounced co-localization in male hiPSC-CMs after ET-1 treatment. (**J**) ERα nuclear intensity in all experimental groups. (**K, L**) demonstrate correlations between ERα-NLRP3 co-localization intensity and (**K**) NLRP3 activation (NLRP3-ASC co-localization) and (**L**) nuclear ERα abundance. Decrease in nuclear ERα abundance with increase in ERα-NLRP3 co-localization is again noted. *p<0.05, **p<0.01, ***p<0.01; ns = not significant by two-way ANOVA with Holm-Šídák post-test. Each data point = one PAH patient (**C-F**) or one hiPSC-CM line in ten different hiPSC-CMs per dish (**I-L)** (means ± SEM).

These data indicate that ERα and NLRP3 directly interact, and that NLRP3 activation is associated with reduced ERα abundance, suggesting this interaction prevents ERα nuclear translocation and activation.

### E2 improves RV function and survival in an ERα-dependent manner

To identify if E2’s effects on RVCM NLRP3 activation are associated with better RV function *in vivo*, we assessed E2’s effects on RV function via pressure-volume loops. We assessed this in male E2-treated MCT-PH rats from *Fig. 2*. Interestingly, E2 treatment indeed was associated with better RV adaptation, evidenced by higher Ees/Ea (*Fig. 11A&B*). E2’s effect size was similar to that of MCC950. E2’s effects on RV-PA coupling were accompanied by beneficial effects on survival (*Fig. 11C*). However, protective effects of E2 on RV-PA coupling and survival were absent in ERα^mut^ rats (from *Fig. 7*), suggesting that functioning ERα is necessary for E2 to exert RV-protective effects (Fig. 11D-F).

**Fig 11:**
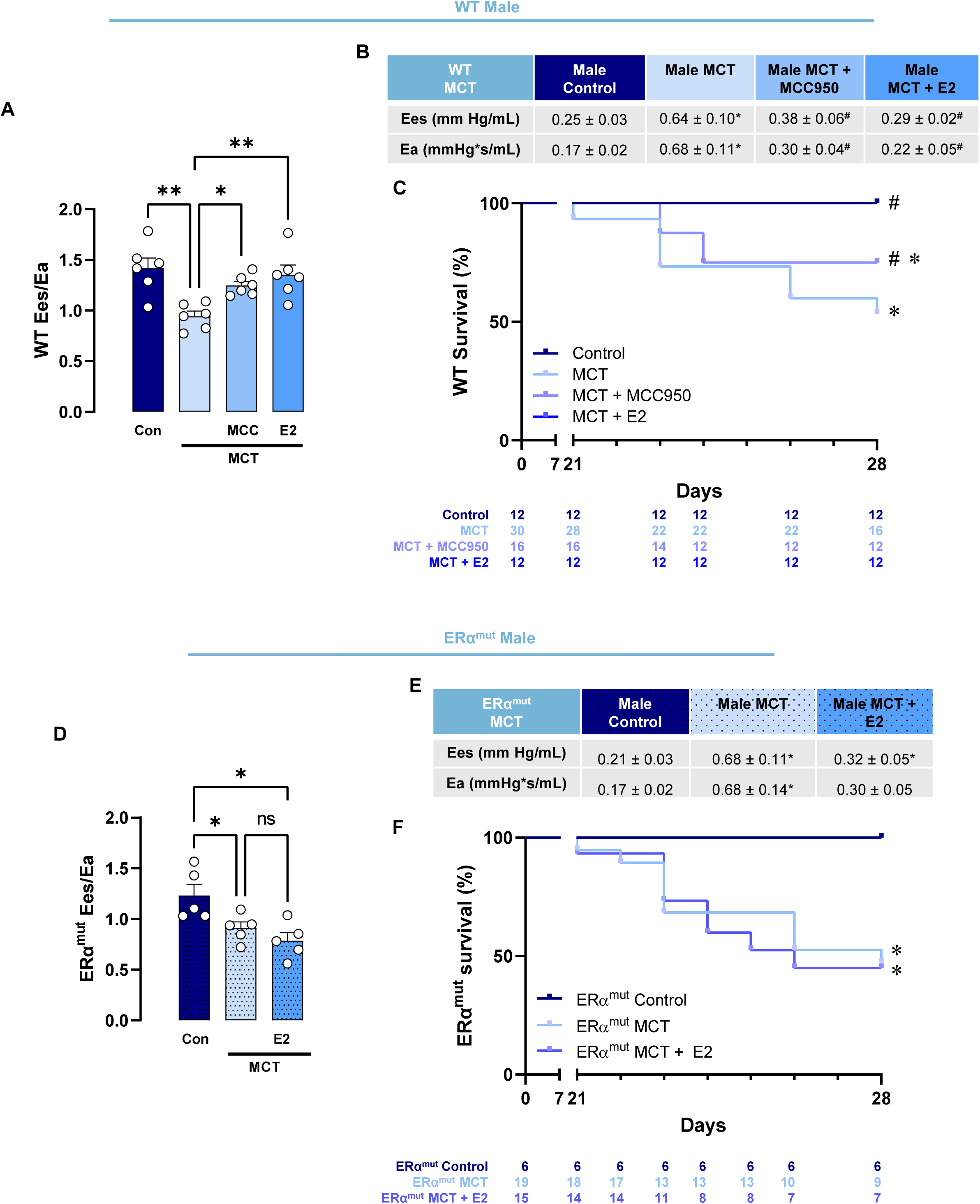
E2 improves RV-pulmonary artery (PA) coupling and survival in male MCT-PH rats in an ERα-dependent manner. Male wild type (WT) or ERα^mut^ MCT-PH rats were treated with MCC950 (WT only) or E2 (WT and ERα^mut^) as outlined in Fig. 2&8. End-systolic elastance (Ees) and arterial elastance (Ea) were measured using pressure-volume loops. Additional animals were utilized for survival studies. (**A**) Ees/Ea in WT MCT-PH groups. Individual values for Ees and Ea are listed in the table in (**B**). (**C**) Survival curves for WT MCT-PH groups. Animal numbers at risk are shown below x-axis. Note that curve for E2 group overlaps with curve of control group due to lack of mortality in either group. (**D-F**) Corresponding studies in ERα^mut^ MCT-PH rats (note that ERα^mut^ MCT-PH rats were not treated with MCC950). (**A, D**) *p<0.05, **p<0.01, ns = not significant by one-way ANOVA with Holm-Šídák post-test; (**B, E**) *p<0.05 vs. male WT or ERα^mut^ control, ^#^p<0.05 vs. male WT or ERα^mut^ MCT by one-way ANOVA with Holm-Šídák post-test; (**C, F**) *p<0.05 vs. male WT or ERα^mut^ control, ^#^p<0.05 for male MCT E2 or MCC950 vs. male WT MCT by Log-rank [Mantel-Cox] test. Each data point represents one rat (means ± SEM).

We also assessed RV-PA coupling and survival in OVX female E2-treated MCT-PH rats from *Fig 2*. As in males, E2 treatment was associated with better RV adaptation (higher Ees/Ea; *Fig. 12A&B*) and better survival (*Fig. 12C*).

**Fig 12:**
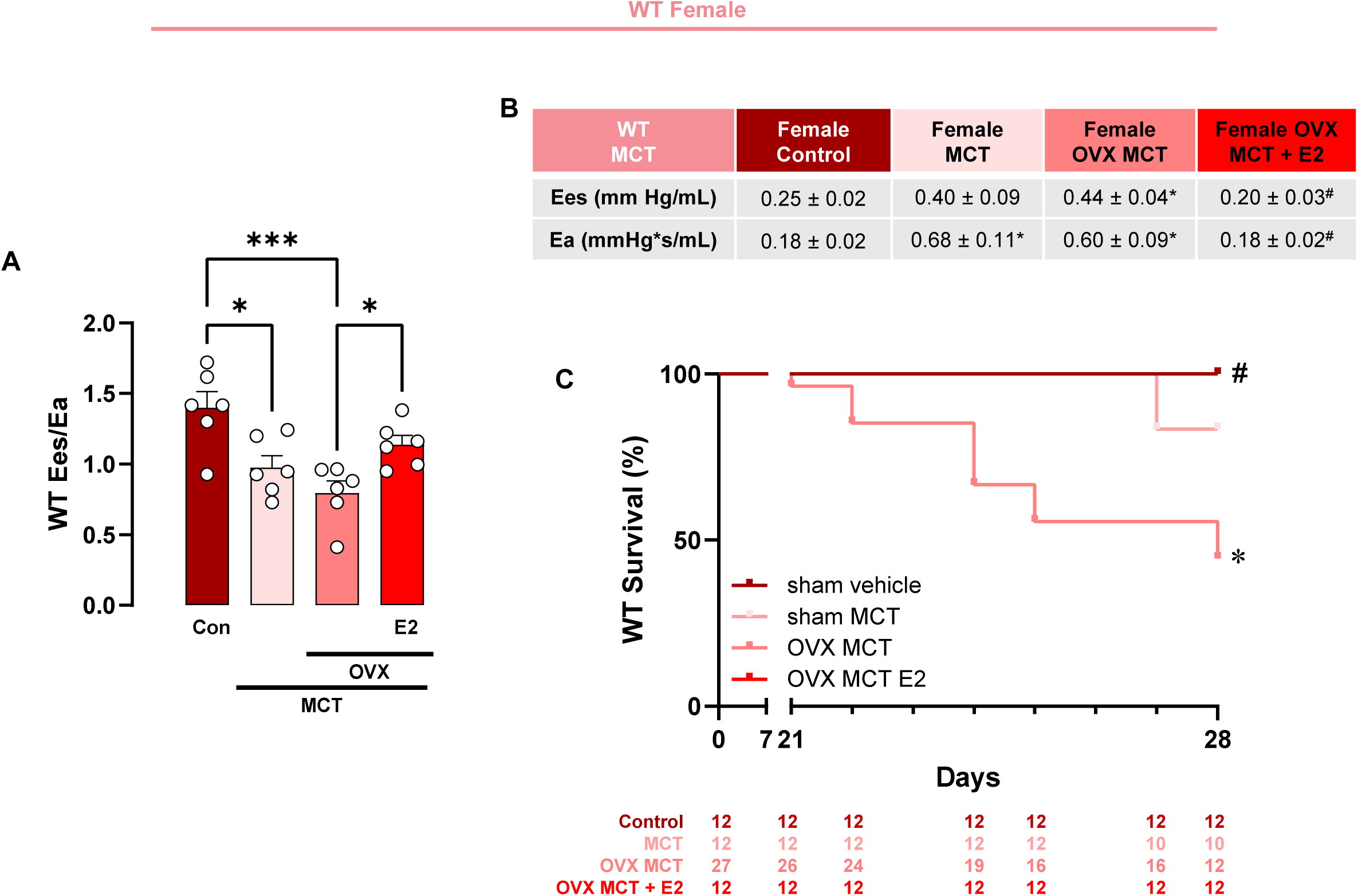
E2 improves RV-pulmonary artery (PA) coupling and survival in female OVX MCT-PH rats. Intact or OVX female wild-type (WT) MCT-PH rats were treated with E2 as outlined in Fig. 2. End-systolic elastance (Ees) and arterial elastance (Ea) were measured using pressure-volume loops. Additional animals were utilized for survival studies. (**A**) Ees/Ea in all experimental groups. Individual values for Ees and Ea are listed in the table in (**B**). (**C**) Survival curves. Animal numbers at risk are shown below x-axis. Note that curve for E2 group overlaps with curve of control group due to lack of mortality in either group. (**A**) *p<0.05 by one-way ANOVA with Holm-Šídák post-test; (**B**) *p<0.05, ***p<0.005 vs. control, ^#^p<0.05 vs. OVX MCT by one-way ANOVA with Holm-Šídák post-test; (**C**) *p<0.05 vs. control, ^#^p<0.05 for MCT E2 vs. OVX MCT by Log-rank [Mantel-Cox] test. Each data point represents one rat (means ± SEM).

These data indicate that E2, via ERα, improves RV adaptation and survival in PH rats with low endogenous E2 states.

## Discussion

We demonstrate for the first time that NLRP3 is upregulated in RVCMs in the setting of RVF and results in RVCM contractile dysfunction. We demonstrate RVCM NLRP3 upregulation in the setting of chronic pressure overload as well as with acute stressors and show that NLRP3 activation and NLRP3-induced RVCM contractile dysfunction are sexually dimorphic and male-biased. Loss of functioning ERα in females eliminates this sex bias, demonstrating that ERα is required for resilience of female RVCMs during pressure overload and NLRP3 activation. E2, via ERα, prevents RVCM NLRP3 activation and NLRP3-induced RVCM contractile dysfunction in males. ERα’s effects on NLRP3 appear to be mediated in a direct manner via ERα-NLRP3 interaction. Sexual dimorphisms in NLRP3 activation and E2 protection were observed with several stimuli and across species. No NLRP3 activation was noted in the LV, suggesting that NLRP3 activation in the setting of PH is RV-specific. These data identified a novel mechanism underlying the well-known sexual dimorphisms in RV adaptation to pressure overload, where pre-menopausal female RVs are more resilient (5–9). Inhibiting NLRP3 signaling via the E2-ERα-axis or with NLRP3 inhibitors may be a novel therapeutic intervention in low endogenous E2 states such as male sex or post-menopause.

While NLRP3 activation in recruited macrophages contributes to RVF development, we now report a novel, macrophage-independent mechanism of NLRP3-induced RV dysfunction. In addition, we identified novel regulators of NLRP3 activation as well as novel NLRP3 downstream targets, thus gaining a better understanding of previously unknown mechanisms of RV and RVCM NLRP3 regulation and signaling. While NLRP3 inflammasome signaling was originally described in immune cells, recent studies demonstrated that NLRP3 also plays a role outside of the immune system (21, 22, 39). The current study further contributes to an emerging paradigm of cardiomyocyte NLRP3 signaling contributing to cardiac dysfunction. To the best of our knowledge, this is the first study of NLRP3 inflammasome signaling specifically in the RV and in RVCMs and to imply cardiomyocyte NLRP3 activation in the pathogenesis of RVF.

We demonstrate for the first time that sex differences exist in RV and RVCM contractile function after NLRP3 activation, and that ERα is responsible for protecting female RVs and RVCMs from NLRP3-induced decreases in contractile function. This is the first study that demonstrates a direct role of ERα in regulating contractile function in the RV.

We previously showed that E2 and ERα in RVCMs upregulate cardioprotective BMPR2-apelin signaling (14). We now demonstrate that effects of E2 and ERα extend beyond this axis and include anti-inflammatory effects by inhibiting NLRP3 inflammasome activation. This suggests that the E2-ERα axis protects RVCM via several pathways. Whether there is crosstalk between E2’s and ERα’s effects on the BMPR2-apelin axis and the NLRP3 inflammasome remains to be determined.

There is controversy on whether estrogens are protective or harmful in PAH (2, 5, 28, 40–42). While effects of E2 in the pulmonary vasculature are complex and cell- and context-dependent, effects of E2 in the RV have consistency been cardioprotective. Based on our current and prior data, enhancing estrogen signaling in the RV would be expected to be beneficial, especially in low endogenous estrogen states such as male sex or post-menopause. However, since endogenous E2 may contribute to pulmonary vascular remodeling in PAH, there has been interest in inhibiting E2 clinically. A recent study pursued aromatase inhibition to alleviate PAH. However, this study, performed in male and pre-menopausal women with relatively mild PAH, did not achieve its primary endpoint (43). No detrimental effects on RV function were noted in this cohort without overt RV dysfunction. We posit that a more precise approach to targeting estrogen signaling in PH is needed. Targeting cardiac ERα may allow for harnessing beneficial estrogenic effects in the RV while avoiding potential detriment in the pulmonary vasculature. Given E2’s pleiotropic effects and potential for undesired effects in the pulmonary vasculature, targeting ERα may represent a more specific approach to leverage beneficial effects of estrogenic signaling. In fact, ERα-specific agonists have been used to treat cardiovascular disorders without exerting any uterotropic effects (44). ERα agonists could target NLRP3 as well as BMPR2 and apelin and exert effects in both the RV as well as the pulmonary vasculature.

While we identified a new mechanism of how E2 and ERα inhibit NLRP3 activation and enhance RCVM contractile function, our studies raise several questions. First, we discovered that ERα and NLRP3 co-localize in RVCMs, suggesting that ERα inhibits NLRP3 via direct interaction. However, the exact mechanism underlying ERα-mediated NLRP3 inhibition remains unknown and is currently under investigation. Furthermore, it remains unknown if E2 and ERα interact with other components of the NLRP3 signaling cascade. The rapid onset of E2’s effects in the Langendorff studies suggests a non-genomic mechanism of action. However, E2 exposure was longer in in vitro and in vivo studies, suggesting that genomic mechanisms likely are at play as well. Of note, E2’s effects are dose-dependent, and NLRP3 inhibition was not consistently observed at higher (>10 nM) concentrations. This suggests that E2 effects in biological systems need to be interpreted in context of the concentrations administered.

Second, we made the surprising finding of increased ERα-NLRP3 co-localization in male RVs, RVCMs, and iPSC-CMs despite less NLRP3-ASC co-localization. This was associated with decreased ERα nuclear localization, suggesting inhibition of ERα nuclear translocation in males once it interacts with NLRP3. Such a constellation would be consistent with reduced genomic ERα signaling in males. Interestingly, reduced ERα nuclear abundance was not paralleled by increased cytoplasmic ERα levels, suggesting potential degradation of ERα in the cytoplasm.

Lastly, it remains unknown which factor(s) activate NLRP3 in RVCMs in PH. Our studies suggest that ET-1 can do this. Other factors, such as RVCM mechanical stretch, oxidative stress, sterile inflammation, and mitochondrial abnormalities may contribute as well and are currently under investigation.

*(Please see supplement for discussion of pulmonary vascular effects of NLRP3, modification of MCT administration in females, and personalized strategies for inhibiting inflammation in low-estrogen states.)*

In summary, our studies identified a novel, therapeutically targetable E2-ERα-NLRP3 axis in RVCMs. RVCM NLRP3 is upregulated in the setting of RVF and results in contractile dysfunction. We provide novel evidence that RVCM NLRP3 upregulation and NLRP3-induced RVCM contractile dysfunction are sexually dimorphic and male-biased.

ERα is required for resilience of female RVCMs in the setting of pressure overload and NLRP3 activation, and E2, via ERα, prevents RVCM NLRP3 activation and NLRP3-induced RVCM contractile dysfunction in males. Inhibiting RVCM NLRP3 upregulation via ERα may represent a novel therapeutic approach in males and postmenopausal females with RVF.

## Notes

### Competing Interest Statement

The authors have declared no competing interest.

