## Supplementary material for "Cardiomyocyte NLRP3 signaling in right heart failure is sexually dimorphic via estrogen receptor α": online data supplement

**Supplemental Methods**

**Study approval**

All human studies were approved by and performed in accordance with the Laval University and Institute Universitaire de Cardiologie et de Pneumologie de Québec biosafety and human ethics committees (CER-20773). All patients or their legal representatives (in case of autopsy) gave informed consent before beginning of the study. Generation of human induced pluripotent stem cell lines and execution of experiments were performed according to the guidance of the Institutional Biosafety Committee University of Minnesota (protocol, 2209-40365H). All rodent experiments were approved by National Jewish Health or Indiana University IACUCs (protocol 2024-0024) and adhered to NIH guidelines for care and use of laboratory animals.

**General**

All experiments and analyses were performed in accordance with recent recommendations (1, 2), including randomization and blinding at the time of intervention, endpoint collection, and analysis.

**Animal models**

**Monocrotaline-induced PH (MCT-PH)**

Male, intact female, or OVX female wild-type (WT) or ERα^mut^ rats (200–250 g) were subcutaneously (s.c.) injected with MCT (60 mg/kg; Sigma-Aldrich) in 3 mL of phosphate-buffered saline (PBS, Thermo Scientific Chemicals). Since female rats typically convert less MCT into the active metabolite dihydro-MCT, we used a higher volume of PBS than standard administration methods to prolong absorption of MCT to allow for more MCT-to-dihydro-MCT conversion to induce a more robust phenotype in females. After MCT injection, subgroups of animals received MCC950 (NLRP3 inhibitor, 10 mg/kg/day, s.c. injection (3)) or E2 (75 μg/kg/day, s.c. pellets; resulting in physiological range plasma E2 levels (4)) for four weeks, followed by endpoint analysis.

**Pulmonary artery banding (PAB)**

Male and female WT rats (150–180 g) were anesthetized by inhaled isoflurane and orotracheally intubated. A small incision was made through the intercostal muscle and extended toward the sternum and the spine, and the thoracic cavity opened. The PA was then dissected, and a 4–0 silk suture was passed around the PA and constricted using a 19 G needle hub as described previously (4). The thoracic cavity was then closed, re-evacuated, and monitored for refilling of air, and muscle layers sutured. Sham controls underwent an identical surgical procedure without PA constriction.

**Hemodynamic assessment in rodents**

Hemodynamic assessments were performed under isoflurane anesthesia (2%) as described previously (4). Right ventricular systolic pressure (RVSP) was measured with a 2-Fr Millar catheter (Houston, TX) using LabVIEW software (National Instruments, Houston, TX).

For pressure-volume loop assessment, the heart and great vessels were exposed via a thoracotomy. A 21-gauge needle was used to puncture the RV wall apically. The needle was immediately replaced by a 1.9F PV admittance catheter (Transonic Scisense, ON, Canada) introduced along the long axis of the chamber. Once PV waveforms recorded were reproducible in steady-state conditions, the inferior vena cava was slowly occluded with a suture line (Fine Science Tools Braided Silk Suture 5-0, Foster City, CA, USA) for a few seconds to change the venous return (occlusion conditions). Three occlusions were performed on each animal. Data were collected in LabChart Software (AD Instruments Inc., Version 8.0 Pro, Colorado Springs, CO, USA).

**Rat RVCM isolation**

Adult rat RVCMs were prepared from ventricular tissue using a Langendorff apparatus (5). Briefly, rats were first given 200 μL heparin (1,000U/mL) via intraperitoneal injection to prevent thrombus formation in coronary arteries. Rats were anesthetized using isoflurane, and following cervical dislocation the heart was excised and immediately cannulated at the aortic root. Hearts were perfused at a constant rate of 4 mL/min at 37°C starting with 4 minutes of perfusion buffer (113 mM NaCl, 4.7 mM KCl, 0.6 mM KH2PO4, 0.6 mM Na2HPO4, 1.2 mM MgSO4, 10 mM HEPES, 12 mM NaHCO3, 10 mM KHCO3, 30 mM Taurine, 10 mM 2,3-butanedione monoxime, 5.5 mM D-(+)-glucose, 10 mM creatine monohydrate, pH 7.4). Subsequently, enzymatic digestion was achieved by 3 minutes of perfusion with calcium-free digestion buffer (350 U/mL Collagenase Type 2, Worthington Biochemical Corporation) followed by 8 minutes of perfusion with digestion buffer containing 50 μM CaCl2. Hearts were removed from the perfusion apparatus, atria were removed, and ventricles were gently mechanically disrupted using transfer pipettes until the tissue was sufficiently digested. The cell suspension was filtered through a 250 μm mesh, and the cardiomyocytes were allowed to settle by gravity for 10 minutes in stopping buffer (10% FBS and 12.5 μM CaCl2 in perfusion buffer). The supernatant was removed and cardiomyocytes were resuspended in an additional 10 mL stopping buffer. CaCl2 was added incrementally to a final concentration of 1.4 mM. After centrifugation at 30 xg, the final cardiomyocyte pellet was resuspended in plating medium (2.5% FBS and 1.4 mM CaCl2 in perfusion buffer) and plated on laminin-coated glass coverslips for 30 minutes in a 12 37°C incubator at 5% CO2. Cardiomyocytes were equilibrated in culture medium (MEM + Earle’s salts + L-glutamine [Gibco 10-010-CV] with 0.2% BSA, 10 mM HEPES, 10 mM NaHCO3, 10 mM creatine monohydrate and 0.5% insulin-transferrin-selenium) for 1 hour prior to assessment.

**Rat RVCM contractility assessment and calcium measurement**

Cellular contractility and relaxation of adult rat cardiomyocytes were measured using the IonOptix system (6). The cells were stimulated at 1 Hz in Tyrode’s solution (40 mM NaCl, 6 mM KCl, 2 mM CaCl2, 1 mM MgCl2, 10 mM glucose, 2.5 mM pyruvate, 5 mM HEPES, pH 7.4, 37°C). Each plate of cardiomyocytes was incubated with FURA2 AM for 30 minutes, and washed solution 3 times with Tyrode. After this, each plate of cardiomyocytes was stimulated for no more than 20 minutes and 3-9 independent traces per plate were measured. IonWizard software (IonOptix) was used to analyze data using edge detection methods. The traces from each myocyte measured were averaged per plate. Rate of shortening (-dL/dt; µm/sec) and FURA2 AM intensity (ΔF/F_0_; amplitude) were measured. Peak height (H) as a percentage of baseline (BL) for sarcomere shortening was calculated and is the percentage of myocyte shortening in relation to baseline; this measurement is indicative of peak ventricular contractility, and H as relative fluorescence unities (RFU) and is the amplitude FURA2 AM fluorescence amplitude in relation to baseline; this measurement is indicative of peak cytosolic free calcium.

**Ex vivo isolated perfused rat heart preparation**

Rat hearts were rapidly isolated and perfused via the aorta at a constant pressure (60 mmHg) with oxygenated Krebs–Henseleit bicarbonate buffer (95% O_2_ + 5% CO_2)_, in a temperature‐controlled (37°C) Langendorff apparatus as previously described(7, 8). Electrodes were placed at the aorta cannula and cardiac apex, and hearts were paced at 350 beats/minute (Grass Instruments, S44 stimulator). Parameters of RV systolic and diastolic function (developed pressure, maximum rate of pressure development [d*P*/d*t* _max_], and maximum rate of relaxation [d*P*/d*t* _min_]) were measured through a high‐compliance fluid‐filled balloon with a diameter of 3 mm inserted into the RV and connected to a pressure transducer. Each balloon was pretested for filling capacity before the beginning of the experiments. Parameters were recorded using a PowerLab data acquisition system (AD Instruments, Australia), and transducer calibration was performed before each experiment following the manufacturer's instructions. Hearts were perfused with LPS (1 μg/mL, 130 min prior to start of dP/dt max measurement and continued throughout measurement) + ATP (2 mM, 10 min prior to start of dP/dt max measurement and continued throughout measurement) or vehicle, in the presence or absence of MCC950 (1 μM; 30 min prior to LPS and continued throughout LPS exposure) or E2 (1 nM; 30 min prior to LPS and continued throughout LPS exposure). Frank-Starling relationship was achieved in each heart while filling the balloon with saline until RV end‐diastolic pressure (RVEDP) increased in 5 mmHg increments from 0 to 40 mmHg. RV NLRP3 activation with LPS+ATP was confirmed by an increase in NLRP3 and ASC colocalization (*Fig. 3&7*).

**Human RV tissue**

Human RV free wall tissue was collected as described previously (9, 10) from warm autopsies (<3 hours after death) or cardiac surgery. The RVF group consisted of patients with congenital heart disease without LV involvement and patients with idiopathic or scleroderma-associated end-stage PAH (age 49.9 ± 5.5 years; 70% female). RVF was defined as decreased tricuspid annular plane systolic excursion and/or death from RVF (9). The presence of RVF was confirmed during autopsy. Control group tissues were obtained from donors with coronary artery disease or aortic stenosis without evidence of PAH or RV hypertrophy by echocardiography or histological analysis(9, 10) (45.1 ± 3.8 years; 77% female).

**Human RV RNA-sequencing analysis**

Previously performed RNA-Seq in human RV tissue from control and decompensated RVF patients(11) deposited in the NCBI Gene Expression Omnibus (GEO, [GSE240941](https://www.ncbi.nlm.nih.gov/geo/query/acc.cgi?acc=GSE240941)) was queried for NLRP3 activators and NLRP3 signaling components.

**Human RV proteomics analysis**

Previously performed proteomic analyses in human RV tissue from control and decompensated RVF patients (12) deposited in the NCBI Gene Expression Omnibus (GEO,  [GSE198618](https://www.ncbi.nlm.nih.gov/geo/query/acc.cgi?acc=GSE198618)) were queried for NLRP3 activators and NLRP3 signaling components.

**Human induced pluripotent stem cell (hiPSC) and hiPSC-derived cardiomyocyte generation**

hiPSCs were obtained, differentiated and validated as described previously (13). Briefly, failed human donor hearts were obtained from the University of Wisconsin. Hearts with prespecified conditions (e.g., coronary artery disease, impaired function, long ischemia time) were excluded. Hearts were flushed with cardioplegia solution, excised, and transported in ice-cold cardioplegia solution. 25-30 g of the left ventricle was minced, homogenized, and digested with digestion media containing DMEM and Liberase TM. The mixture was filtered and centrifuged to obtain cell pellets. The cells were resuspended in complete media and plated. Non-adherent cells were washed away, and adherent cells were cultured until they reached ~90% confluence.

Adult left ventricular cardiac fibroblasts (aLVCFs) were reprogrammed into human induced pluripotent stem cells (hiPSCs). Six hiPSC lines were reprogrammed, from both male and female donors aged 21-34 years (see *Suppl. Table 1*). These were reprogrammed using CytoTune Sendai virus according to the suppliers’ directions. The lines were validated for both Sendai virus depletion pluripotency as detailed in the publication.

For maintenance the hiPSCs were cultured on Cultrex RGF-coated plates in mTesR media. Upon reaching 70-80% confluency, cells were passaged using ReLesR onto Cultrex-RGF-coated plates. For hiPSC-CM differentiations the cells were plated onto Cultrex-RGF-coated plates in mTesR. The differentiations was initiated two days post seeding using in RPMI/B27 minus insulin media with varying concentrations of CHIR depending on the lines. 48-hours later the media was changed to RPMI/B27 with insulin containing 7.5 uM IWP2. After this the media was changes every 2-3 days with fresh RPMI/B27 with insulin. Once differentiated the hiPSC-CM were replated and purified using lactate metabolic selection media. On day 40, CM purity was assessed via flow cytometry for the CM-specific marker cardiac troponin T (cTnT). All 6 lines displayed >95% cTnT population (See suppl. table 1 for details) (14, 15).

**NLRP3 activation assessment in human iPSC-CM and rat RVCMs**

To investigate the NLRP3 activation in human iPSC-CMs and rat RVCMs, these cells was treated with NLRP3 activators in the presence or absence of MCC950 or E2. Briefly, after differentiation and purification, iPSC-CMs were seeded onto chamber slides at a densirty of 1000 cells/cm^2^ and treated with NLRP3 activators endothelin-1 (ET-1; 1 nM, 4 h) (16) in absence or presence of E2 (1 nM, 24 h) or NLRP3 inhibitor MCC950 (1 µM, 30 min) (4, 17).

RVCMs isolated from male or female WT or ERα^mut^ MCT rats treated with E2 or MCC950 and equilibrated in a culture medium for 24 hours. To confirm the NLRP3 activation without the MCT interference, rat RVCMs isolated from healthy male or female WT or ERα^mut^ rats were treated with NLRP3 activators lipopolysaccharide (LPS; 1 µg/mL, 4 h) and ATP (2 mM, 10 min) (3, 18) in the absence or presence of E2 (1 nM, 24 h) or NLRP3 inhibitor MCC950 (1 µM, 30 min). In addition, rat RVCMs were treated with endothelin-1 (ET-1; 10 nM, 4 h) in the absence or presence of E2 (1 nM, 24 h)(16) or NLRP3 inhibitor MCC950 (1 µM, 30 min). After the incubation, the cells separated for functional (contractility and calcium measurement), western blot, or immunofluorescence studies.

**Immunofluorescence studies in human RV tissue**

Immunofluorescence labeling for NLRP3, ASC, and ERα was performed using formalin-fixed paraffin-embedded human RV sections (4μm thick). Antigen retrieval was performed by heating samples in 0.01 M citrate buffer (10 mM Sodium Citrate, 0.05% Tween-20, 226 pH 6.0). Primary antibodies used were rabbit monoclonal anti-NLRP3 (1:200; NBP2-67639, Novus Bio, RRID:AB_3353608), goat Anti-TMS1/ASC antibody (1:200; ab175449, ABCAM, RRID:AB_3096354), and mouse anti-ERα antibody (1:500; NBP2-61764, Novus Bio, RRID:AB_3351174). Secondary fluorochrome-conjugated anti-rabbit antibody (1:500; Alexa Fluor 488; A-21210, Thermo Fisher, RRID:AB_2535796), fluorochrome-conjugated anti-goat antibody (1:200; Alexa Fluor 594; A-11057, Thermo Fisher, RRID:AB_2534104), fluorochrome-conjugated anti mouse antibody [(1:200; Alexa Fluor 645; A-31571, Thermo Fisher, RRID:AB_162542) were used. DAPI staining was used to visualize nuclei. Negative controls were performed during each experiment by incubating secondary antibodies alone or by using rabbit (1:500; sc-2027, Santa Cruz, RRID:AB_737197), mouse (1:500; sc-2025, Santa Cruz , RRID:AB_737182), and goat (1:500; 31245, Invitrogen, RRID:AB_2421585) IgG isotype control and following all protocol steps, including incubation with the secondary antibody. Rat RVs imaging for each antibody was taken at the identical exposure time for each experimental condition/magnification. Images were taken using a Confocal Zeiss LSM 700 microscope with camera, and Zen Black 3.5 100x magnification. Human RV images were taken using a slide scanner, AxioScan with camera, and Zen Black 3.5 software at 20x. In addition, representative images in humans were taken using a Zeiss Observer Z1 with camera, and Zen Black 3.5 software at 40x.

**Immunofluorescence studies in rat RVCMs and human iPSC-CMs**

CMs (1000 cells/cm^2^) were plated on Nunc® Lab-Tek® Chamber Slide™ system 8 wells, Permanox® slide, 0.8 cm2/well, sterile coated with laminin. To verify viability, prior to plating, cells were stained with Trypan Blue solution (Corning). Cells were fixed with 4% paraformaldehyde and blocked with background terminator (biocare, BT967). Primary antibodies used were rabbit monoclonal anti-NLRP3 (1:200; NBP2-67639, Novus Bio, RRID:AB_3353608), goat Anti-TMS1/ASC antibody (1:200; ab175449, ABCAM, RRID:AB_3096354), and mouse anti-ERα antibody (1:500; NBP2-61764, Novus Bio, RRID:AB_3351174). Secondary fluorochrome-conjugated anti-rabbit antibody (1:500; Alexa Fluor 488; A-21210, Thermo Fisher, RRID:AB_2535796), fluorochrome-conjugated anti-goat antibody (1:200; Alexa Fluor 594; A-11057, Thermo Fisher, RRID:AB_2534104), fluorochrome-conjugated anti mouse antibody [(1:200; Alexa Fluor 645; A-31571, Thermo Fisher, RRID:AB_162542) were used. Images generated by the samples stained by Alexa Fluor 645 were later changed to red using Zen Black 3.5 software (ZEISS XRM Technology)], and anti-fade DAPI (Thermo Fisher) mounting media were used. Images were taken using a Confocal Zeiss LSM 700 microscope with camera, and Zen Black 3.5 software at 100x (hiPSC-CMs) 100x (Rat RVCMs) magnification.

**Rat RV tissue homogenization**

Rat RV tissue or RVCMs were homogenized using an Omni international tissue grinder (ThermoFisher) in ice-cold RIPA lysis buffer (Thermo Fisher) containing proteinase inhibitor cocktail (EMD-Millipore-Sigma Aldrich, St. Louis, MO) and PhosStop inhibitor cocktail 9 (Roche, Pleasanton, CA). After homogenization, lysate was sonicated for ten one-second pulses at 100% power and then centrifuged. The supernatant was saved and used as RV or LV lysate. Cell lysis RVECs and RVCMs were lysed using 10x Cell lysis buffer diluted with molecular biology grade water (ThermoFisher) and supplemented with Cell Signaling Biotechnology (Danvers, Mass) proteinase inhibitor cocktail (EMD-Millipore-Sigma Aldrich, St. Louis, MO) and PhosStop inhibitor cocktail (Roche, Pleasanton, CA). Cells were lysed using the manufacturers protocol.

**Western blot analysis**

Protein concentration was measured using Bradford Protein Assay (Pierce-Thermo Fisher). Rat RV tissue and RVCM were collected and homogenized as described previously. Rabbit monoclonal caspase-1 Antibody (14F468) (1:1000; NB100-56565, Novus Bio, RRID:AB_837823), IL-1 beta/IL-1F2 Antibody (1:1000; NB600-633, Novus Bio, RRID:AB_10001060) primary antibodies were used on rat RV tissue and RVCMs homogenates. All antibodies were diluted in Pierce Protein-Free T20 blocking buffer 206 (ThermoFisher). Goat-anti-rabbit HRP (1:20000; AC2114, Azure BioSystems, RRID:AB_3661796) and Goat-anti-mouse HRP secondary antibody. HRP (1:20000; AC2115, Azure BioSystems, RRID:AB_3661795) secondary antibodies were diluted in Pierce Protein-Free T20 Blocking 10 208 Buffer. Rats RV and RVCM Western blots were normalized to stain free gels total Protein (Bio-rad). Densitometry was performed using Image J.

**Immunoprecipitation studies**

RVCMs from male and female WT rats were serum-starved overnight and then harvested. After Cells were lysed and incubated overnight with rotation in a cold room with Super agarose beads (Santa Cruz Biotech) bound with rabbit monoclonal anti-NLRP3 (1:500; NBP2-12446, Novus Bio, RRID:AB_2750946). Beads were then washed, followed by the addition of sample loading buffer directly to the beads. Samples were then boiled and run on a western blot. anti-ERα antibody (1:200; NB300-560, Novus Bio, RRID:AB_10001032) was used to detect complex formation. IPs with IgG controls were used to demonstrate absence of non-specific antibody binding.

**Statistics**

Results are expressed as mean ± SEM. At least 3 biologically independent experiments (run in technical duplicates) were performed for all in vitro studies and reported as *N*. Statistical analyses were performed with GraphPad Prism 9. Sample sizes were estimated by power calculation. Two-tailed Student’s *t* test, as well as one-way ANOVA and two-way ANOVA with Holm-Šidák post hoc correction were used for comparison of experimental groups. Correlations were determined using Pearson’s coefficient (*R*). Survival analysis was determined using Simple Survival Analysis (Kaplan-Meier). Statistically significant difference was accepted at *P* less than 0.05.

**Supplemental Discussion**

Of note, loss of functioning ERα also results in a worse PH phenotype in the pulmonary vasculature in female MCT rats (19), suggesting that this receptor protects other compartments of the cardiopulmonary system in females. This is also evidenced by experiments demonstrating upregulation of homeostasis and survival pathways in pulmonary artery endothelial cells from PAH patients after treatment with ERα agonist (19).

We modified the MCT rat model by increasing the volume of MCT solution administered in both males and females. Interestingly, this led to comparable hemodynamic alterations between male and female rats. In fact, the female rats used in this study had more RV hypertrophy than their male counterparts (*Suppl Table 3*). We believe that the higher volume of subcutaneous MCT administered to females resulted in a prolonged absorption period, thus allowing for more conversion of MCT to the active metabolite dihydro-MCT and a more robust PH phenotype in females. This strategy may allow for overcoming limitations of traditional MCT models, where females, due to less cytochrome P450 activity, typically generate less dihydro-MCT and have a less severe phenotype (20, 21). However, we did not rely on the MCT model alone, and studies in PAB rats, hiPSC-CMs, and human RV tissue corroborated results obtained in MCT rats.

Many studies of anti-inflammatory approaches for heart failure or PAH have been negative or equivocal and exhibited considerable inter-subject variability (22-24). One theory for the relative lack of efficacy or modest effect sizes observed of such approaches is that the populations studied were too heterogenous and exhibited various degrees of inflammation activation. The finding that NLRP3 inflammasome activation in RVF is not homogenous but limited to male sex as well as sex hormone loss (OVX) or ERα loss-of-function conditions in females (which in humans occur with menopause) identifies a potential novel therapeutic target that allows for a more personalized treatment approach for RV failure and a more targeted approach to treating inflammation in PAH. We suggest that specifically targeting NLRP3 inflammasome activation in low estrogen states represents a potential novel therapeutic strategy. Importantly, NLRP3 inflammasome signaling and its downstream target activation not only can be targeted with MCC950, but also with newer agents such as SR9009 or the NINJ1 antibody D1 (25-27). Since NLRP3 is also activated in the pulmonary artery wall cells in PH (25), targeting NLRP3 could be a two-pronged approach with benefits in both the RV as well as the pulmonary vasculature, making this an “ideal” treatment approach that targets both components of the cardiopulmonary axis in PH.
