## Supplementary material for "Cardiomyocyte NLRP3 signaling in right heart failure is sexually dimorphic via estrogen receptor α": Fig. E

### Supplemental tables

| Line | Sex | Age | Race | Clinical history |
|---|---|---|---|---|
| **M4** | Male | 21 | Caucasian | Previously healthy, normal EF by echocardiogram |
| **M7** | Male | 21 | Caucasian | Previously healthy, normal EF by echocardiogram |
| **M9** | Male | 23 | Caucasian | Previously healthy, normal EF by echocardiogram |
| **F5** | Female | 26 | Caucasian | Previously healthy, normal EF by echocardiogram |
| **F6** | Female | 34 | Caucasian | Previously healthy, normal EF by echocardiogram |
| **F7** | Female | 33 | Caucasian | Previously healthy, normal EF by echocardiogram |

**Table E1: Characterization of left ventricular fibroblast donors used for generation of human iPSCs and human iPSC-derived cardiomyocytes.** Lines represent the cell lines generated by each donor. EF = ejection fraction.

| PAH patients | Male Control | Male PAH | Female Control | Female PAH |
|---|---|---|---|---|
| **RVSP (mmHg)** | 19.89 ± 2.81 | 100** | 24.00 ± 3.61 | 86.00 ± 10.52*** |
| **TAPSE (mm)** | 24.33 ± 1.20 | N/A | 22.00 ± 2.00 | 14.60 ± 0.93** |
| **Cardiac Index (L/min/m$^2$)** | 2.83 ± 0.14 | 1.73 | 2.87 ± 0.10 | 1.65 ± 0.29** |

**Table E2: Hemodynamic and structural characterization of PAH and control RVs used for transcriptomic and proteomic quantification of NLRP3 activation in Fig. 1.** RSVP = right ventricular systolic pressure; TAPSE = tricuspid annular plane systolic excursion, CI = cardiac index. N = 3 to 9 per group, except for male PAH where n=1 per group. **p<0.01, ***p<0.001 vs respective control by one-way ANOVA with Holm-Šídák post-test (means ± SEM).

| PAH patients | Male Control | Male PAH | Female Control | Female PAH |
|---|---|---|---|---|
| **mPAP (mmHg)** | N/A | 54.00 ± 11.24 | N/A | 60.67 ± 10.23 |
| **RVSP (mmHg)** | 21.00 ± 2.00 | 72.00 ± 28.00 | 36.67 ± 8.57 | 82.67 ± 7.86 |
| **TAPSE (mm)** | 26.00 ± 3.00 | 14.78 ± 3.52 | 22.00 ± 5.00 | 15.22 ± 0.86 |
| **Cardiac Index ($L/min/m^2$)** | 1.65 ± 0.59 | 3.20 | 2.57 | 1.58 ± 0.27 |

**Table E3: Hemodynamic and structural characterization of PAH and control RVs used for immunofluorescence quantification of NLRP3 activation in Fig. 1.** mPAP = mean pulmonary artery pressure, RSVP = right ventricular systolic pressure; TAPSE = tricuspid annular plane systolic excursion, CI = cardiac index. N = 1 to 6 per group (means ± SEM).

| | Male Control | Male MCT | Male MCT + MCC950 | Male MCT + E2 |
|---|---|---|---|---|
| RVSP (mmHg) | 28.32 ± 1.74 | 48.45 ± 7.37* | 35.60 ± 5.86# | 30.45 ± 5.97# |
| Fulton index (RV/LV+S) | 0.33 ± 0.02 | 0.55 ± 0.06* | 0.43 ± 0.08# | 0.38 ± 0.02# |
| | **Female Control** | **Female MCT** | **Female OVX MCT** | **Female OVX MCT E2** |
| RVSP (mmHg) | 27.24 ± 3.38 | 43.47 ± 5.13* | 51.06 ± 7.35* | 26.89 ± 2.34$ |
| Fulton index (RV/LV+S) | 0.37 ± 0.02 | 0.66 ± 0.11* | 0.67 ± 0.06* | 0.40 ± 0.01$ |

**Table E4: Hemodynamic and structural characterization of RVs from control or monocrotaline (MCT) PH wild-type rats used in Fig. 2.** RVSP = right ventricular systolic pressure, RV/LV+S = right ventricle weight / left ventricle weight + septum weight. N=6 per group. *p<0.05 vs male or female control, #p<0.05 vs male MCT, $p<0.05 vs female OVX MCT by one-way ANOVA with Holm-Šídák post test (means ± SEM).

| | Male Sham | Male PAB | Female Sham | Female PAB |
|---|---|---|---|---|
| **RVSP (mmHg)** | 27.53 ± 1.38 | 62.55 ± 4.07* | 26.01 ± 1.03 | 63.27 ± 7.52* |
| **Fulton index (RV/LV+S)** | 0.32 ± 0.028 | 0.62 ± 0.04* | 0.28 ± 0.01 | 0.45 ± 0.04* |

**Table E5: Hemodynamic and structural characterization of RVs from control or pulmonary artery banding (PAB) wild-type rats used in Fig. S3.** RVSP = right ventricular systolic pressure, RV/LV+S = right ventricle weight / left ventricle weight + septum weight. N=5 per group. *p<0.05 vs male or female sham by one-way ANOVA with Holm-Šídák post test (means ± SEM).

| ERα$^{mut}$ MCT | Male Control | Male MCT | Male MCT + E2 | Female Control | Female MCT | Female MCT E2 |
|---|---|---|---|---|---|---|
| **RVSP (mmHg)** | 28.27 ± 3.47 | 47.83 ± 1.52* | 51.99 ± 2.35* | 33.43 ± 1.85 | 46.53 ± 4.58* | 56.36 ± 4.81* |
| **Fulton index (RV/LV+S)** | 0.34 ± 0.02 | 0.55 ± 0.06* | 0.43 ± 0.08 | 0.37 ± 0.02 | 0.67 ± 0.11* | 0.37 ± 0.02 |

**Table E6: Hemodynamic and structural characterization of RVs from control or monocrotaline (MCT) PH ERα$^{mut}$ rats used in Fig. 8.** RVSP = right ventricular systolic pressure, RV/LV+S = right ventricle weight / left ventricle weight + septum weight. N=6 per group. *$p<0.05$ vs male or female control by one-way ANOVA with Holm-Šídák post-test (means ± SEM).

### Supplemental figures

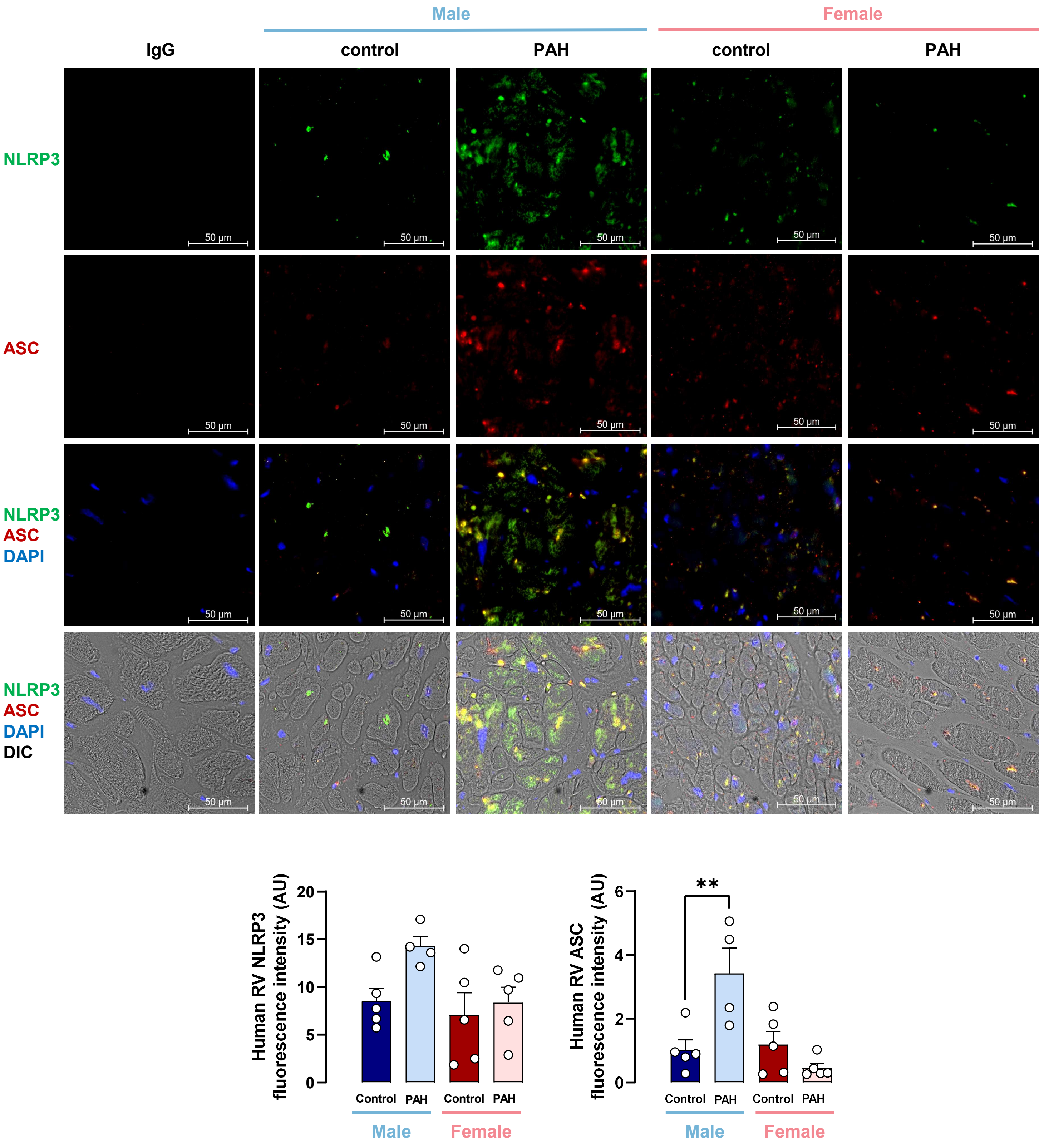

**Fig. E1: Male but not female human pulmonary artery hypertension patients with right ventricle (RV) failure exhibit increased NLRP3 and ASC expression.** Representative immunofluorescent images and quantification of fluorescence intensity of NLRP3 and ASC in RVs from male or female patients or controls. Images were obtained using Confocal Zeiss LSM 700 microscope and Zen Black 3.5 software at 20x magnification. RV samples were stained for NLRP3 (green), ASC (red) and DAPI (blue). Fluorescence intensity images were analyzed in the whole RV sample. Representative images were digitally zoomed 50x. **$p<0.01$ by two-way ANOVA followed by Holm-Šídák post-test. Each data point represents an RV sample from one patient (means ± SEM). AU = arbitrary units. DIC = Differential interference contrast.

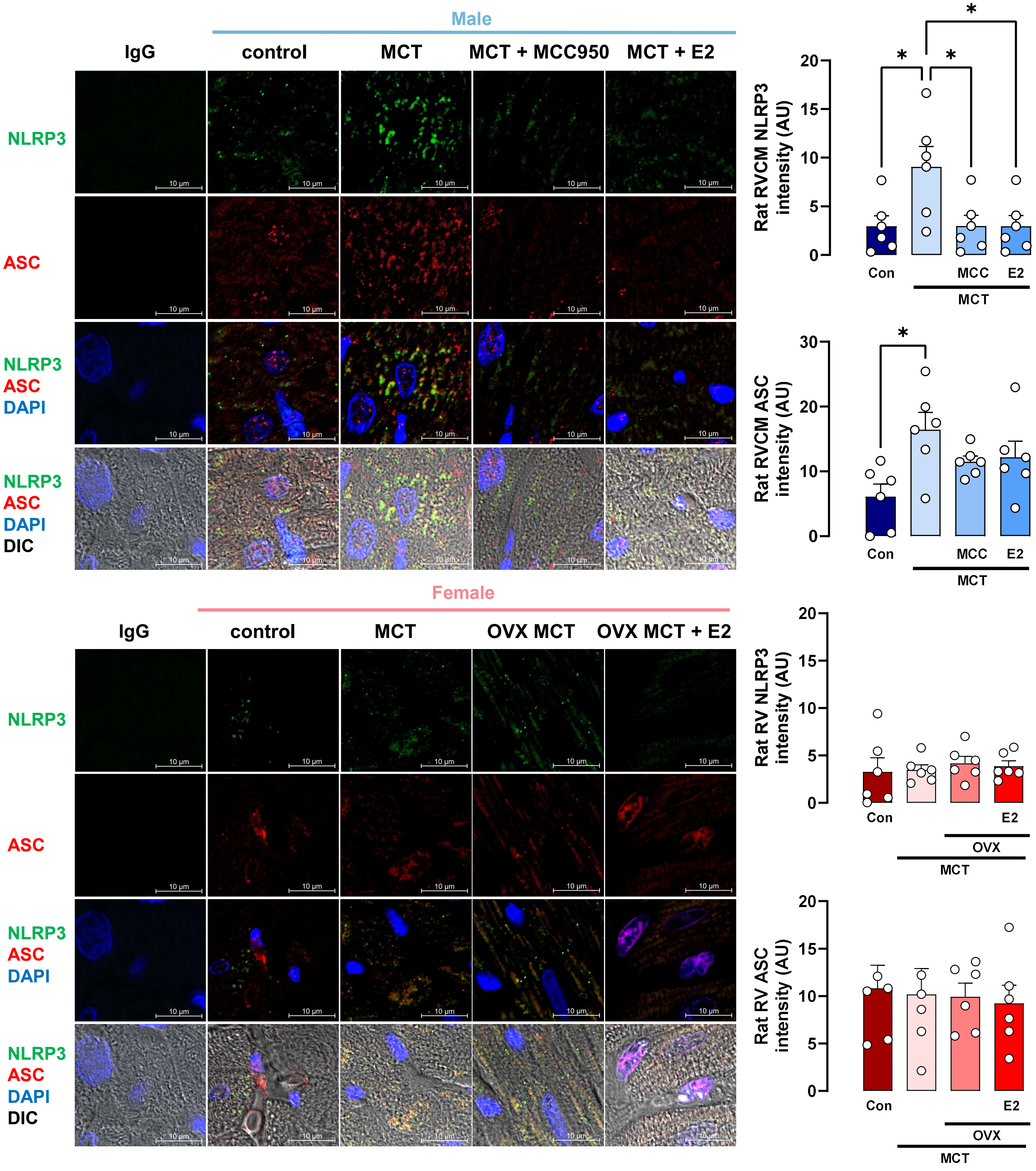

Male
IgG
control
MCT
MCT + MCC950
MCT + E2
NLRP3
ASC
NLRP3
ASC
DAPI
NLRP3
ASC
DAPI
DIC
10 µm
Female
IgG
control
MCT
OVX MCT
OVX MCT + E2
Rat RVCM NLRP3 intensity (AU)
Con
MCC
E2
MCT
Rat RVCM ASC intensity (AU)
Rat RV NLRP3 intensity (AU)
OVX
Rat RV ASC intensity (AU)

**Fig. E2: RV NLRP3 and ASC expression in rats with monocrotaline (MCT)-induced PH exhibits a male bias that is due to inhibitory effects of 17β-estradiol (E2).** RVs were isolated from male or intact or ovariectomized (OVX) female control or MCT rats. Subgroups of MCT-PH males were treated with E2 (75 μg/kg/day by subcutaneous [s.c.] pellets) or NLRP3 inhibitor MCC950 (10 mg/kg by s.c. injection) for 4 weeks (starting at MCT injection). Subgroups of OVX MCT-PH females were treated with E2 (75 μg/kg/day by s.c. pellets) for 4 weeks (starting at MCT injection). Representative immunofluorescence images and quantification of fluorescence intensity of RV NLRP3-ASC colocalization. RVs were stained for NLRP3 (green), ASC (red), and DAPI (blue). Fluorescence intensity was analyzed in ten different fields, and values were then averaged. Images were obtained at 100x magnification. Representative images were digitally zoomed 5x. *$p<0.05$ by one-way ANOVA with Holm-Šídák post-test. Each data point = one rat (means ± SEM). AU = arbitrary units. DIC = differential interference contrast.

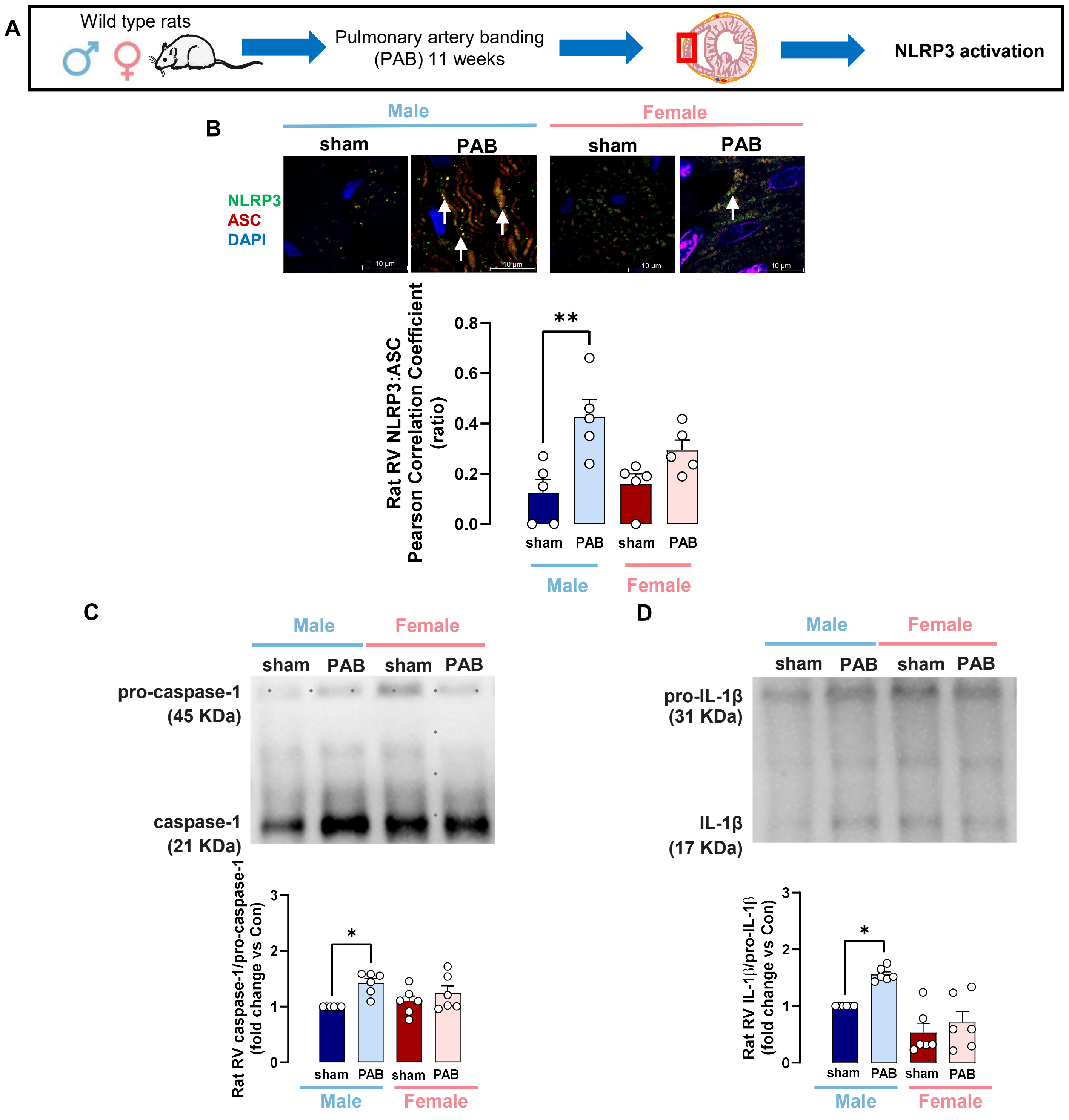
A
Wild type rats
Pulmonary artery banding (PAB) 11 weeks
NLRP3 activation
B
Male
Female
sham
PAB
NLRP3
ASC
DAPI
10 µm
Rat RV NLRP3:ASC Pearson Correlation Coefficient (ratio)
0.0
0.2
0.4
0.6
0.8
**
C
pro-caspase-1 (45 KDa)
caspase-1 (21 KDa)
Rat RV caspase-1/pro-caspase-1 (fold change vs Con)
*
D
pro-IL-1β (31 KDa)
IL-1β (17 KDa)
Rat RV IL-1β/pro-IL-1β (fold change vs Con)
0
1
2
3

**Fig. E3: Male rats with pulmonary artery banding (PAB)-induced RV failure exhibit more pronounced RV NLRP3 inflammasome activation than female PAB rats.** (**A**) Experimental design. (**B**) Representative immunofluorescence images and quantification of fluorescence intensity of NLRP3-ASC colocalization in RVs from male (blue shaded bars) or female (red shaded bars) sham or PAB rats. Images were obtained at 100x magnification. RVs were stained for NLRP3 (green), ASC (red), and DAPI (blue). Co-localization images were analyzed in 10 different fields, and values were then averaged. Representative images were digitally zoomed 5x. NLRP3 and ASC co-localization (white arrows) were quantified by determining Pearson correlation coefficient. Representative Western blot images and densitometric quantification of (**B**) caspase-1/pro-caspase-1 ratio and (**C**) IL-1β/pro-IL-1β ratio in RVs from male or female sham or PAB rats. $^*p<0.05$, $^{**}p<0.01$ by one-way two-way ANOVA with Holm-Šídák post test. Each data point = one rat (means ± SEM).

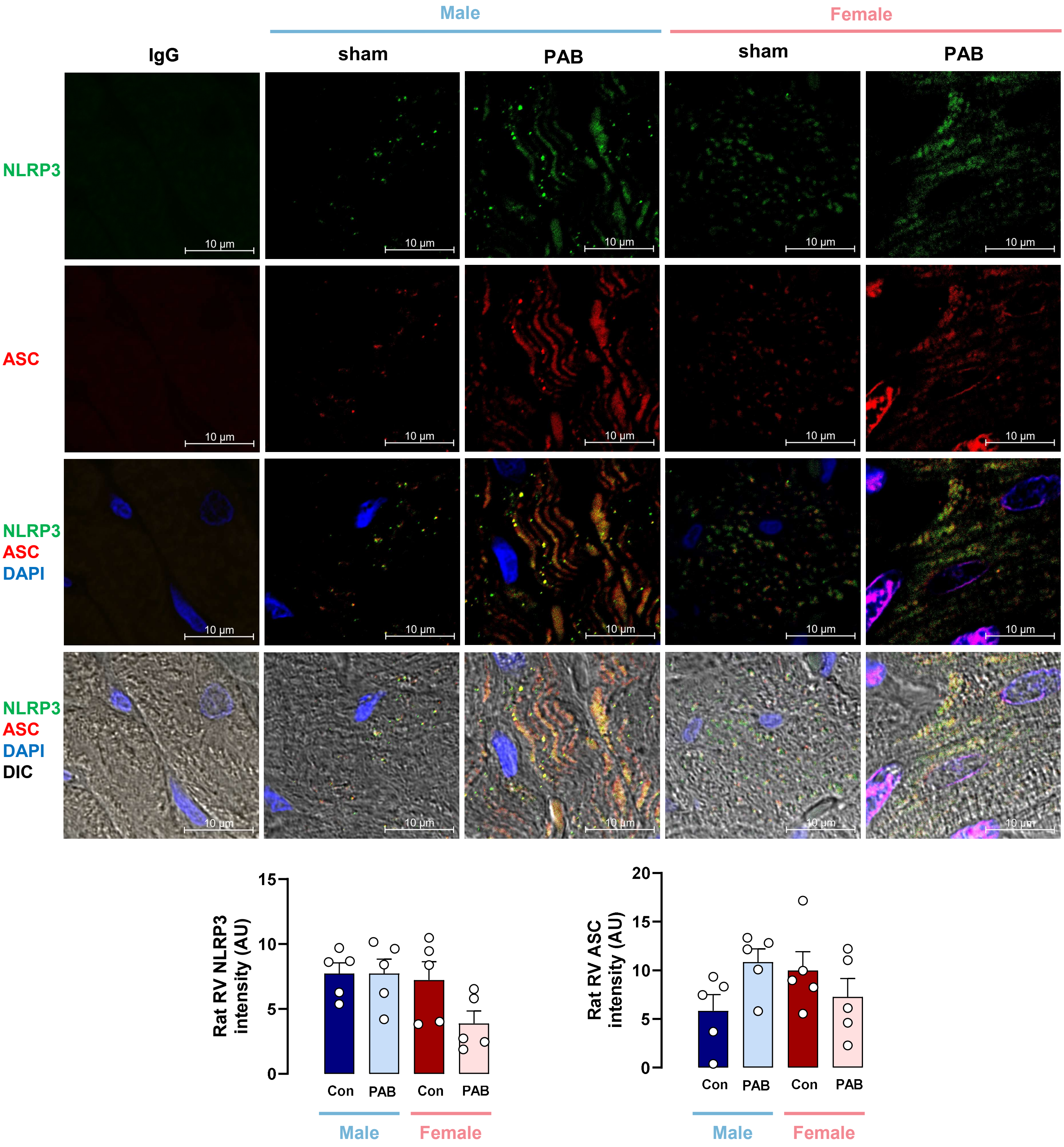
Male
Female
IgG
sham
PAB
sham
PAB
NLRP3
ASC
NLRP3
ASC
DAPI
NLRP3
ASC
DAPI
DIC
10 µm
Rat RV NLRP3 intensity (AU)
15
10
5
0
Con
PAB
Con
PAB
Male
Female
Rat RV ASC intensity (AU)
20
15
10
5
0
Con
PAB
Con
PAB
Male
Female

**Fig. E4: NLRP3 and ASC expression in male and female pressure-overloaded RVs from pulmonary artery banding (PAB) rats**. Representative immunofluorescent images, quantification of fluorescence intensity of NLRP3, and ASC in sham and PAB male and female rat RVs. Images were obtained using a Confocal Zeiss LSM 700 microscope and Zen Black 3.5 software at 100x magnification. Rat RV samples were stained for NLRP3 (green), ASC (red), and DAPI (blue). Fluorescence intensity images were analyzed in 10 different fields, and representative images were digitally zoomed 5x. *$p<0.05$ male or female sham vs PAB by two-way ANOVA followed by Holm-Šídák post-test. Each data point = one rat (means ± SEM). DIC = differential interference contrast.

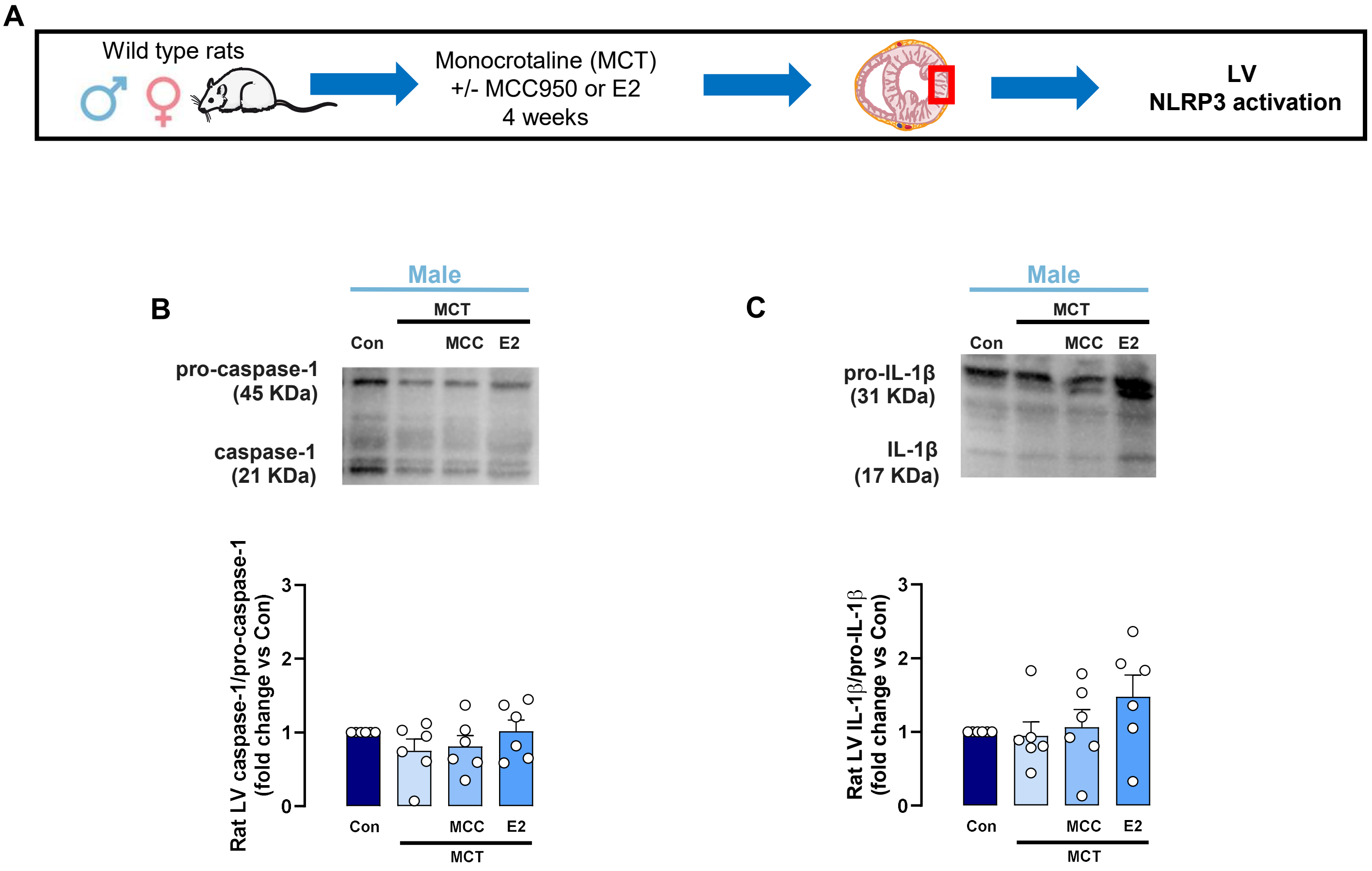


**Fig. E5: Lack of NLRP3 inflammasome activation in left ventricles (LVs) of male MCT-PH rats**. (**A**) Experimental design. Representative Western blot images and densitometric quantification of (**B**) caspase-1/pro-caspase-1 ratio and (**C**) IL-1β/pro-IL-1β ratio in RVCMs from male or female control or MCT rats as well as male MCT-PH rats treated with MCC950 or E2. Data not significant by one-way ANOVA with Holm-Šídák post-test. Each data point = LV sample from one rat (means ± SEM).

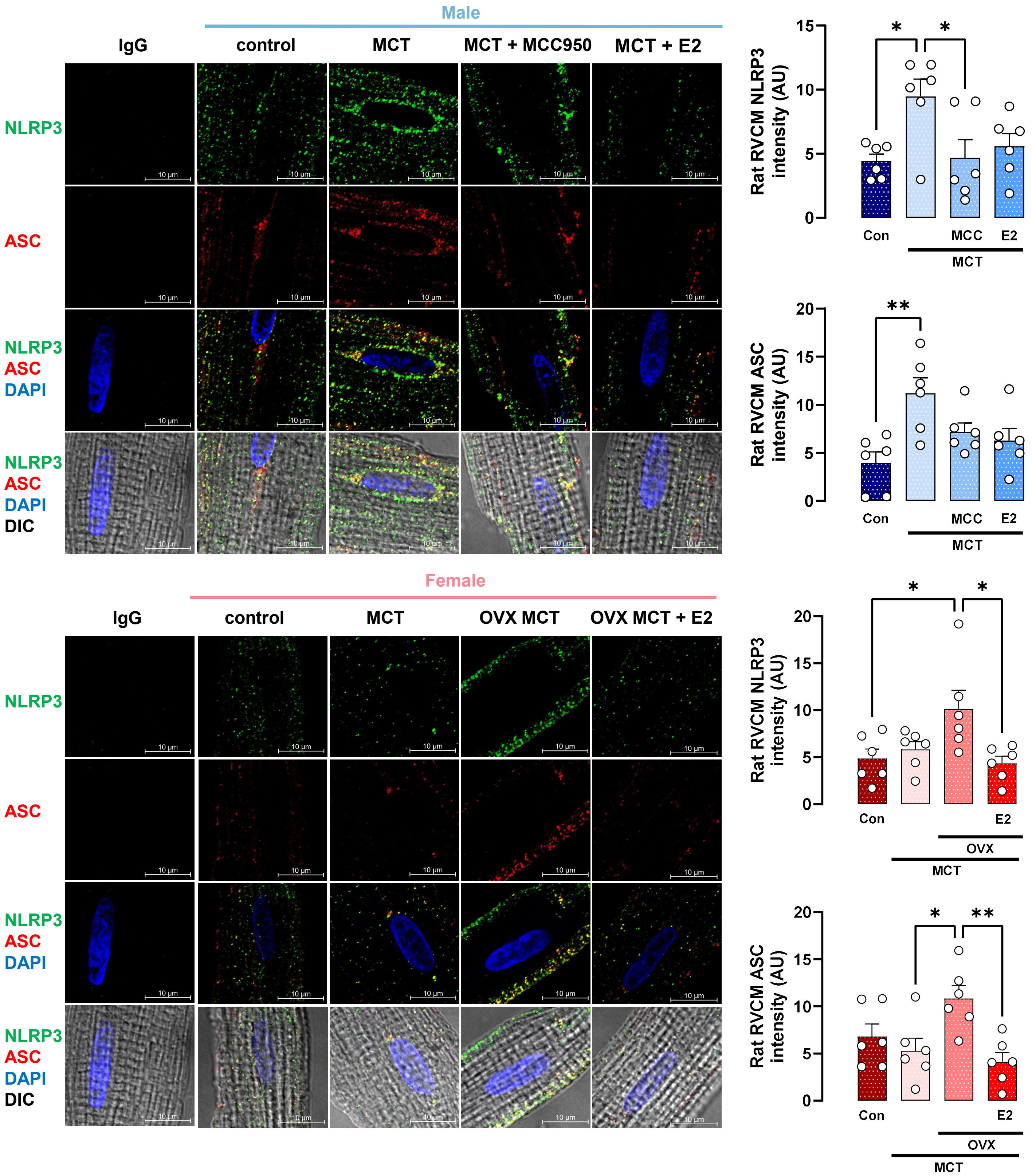
Male
IgG
control
MCT
MCT + MCC950
MCT + E2
NLRP3
ASC
NLRP3
ASC
DAPI
NLRP3
ASC
DAPI
DIC
10 μm
Rat RVCM NLRP3 intensity (AU)
Con
MCC
E2
MCT
Rat RVCM ASC intensity (AU)
Female
IgG
control
MCT
OVX MCT
OVX MCT + E2
OVX

**Fig. E6: RV cardiomyocyte (RVCM) NLRP3 and ASC expression in MCT-PH rats are sexually dimorphic, male-biased, and attenuated by E2**. Representative immunofluorescence images and quantification of fluorescence intensity of NLRP3 and ASC RVCMs from male or female control or MCT rats. Subgroups of MCT-PH males were treated with NLRP3 inhibitor MCC950 (10 mg/kg by subcutaneous [s.c.] injection) or E2 (75 µg/kg/day by s.c. pellets) for 4 weeks (starting at MCT injection). Subgroups of MCT-PH females were ovariectomized (OVX) and treated with E2 (75 µg/kg/day by s.c. pellets) for 4 weeks (starting at MCT injection). Rat RVCMs were stained for NLRP3 (green), ASC (red), and DAPI (blue). Images were obtained at 100x magnification. Fluorescence intensity was analyzed in ten different RVCMs per animal, and averaged values for each animal are shown. Representative RVCM images were digitally zoomed 5x. *$p<0.05$, **$p<0.01$ by one-way ANOVA with Holm-Šídák post-test. Each data point = RVCMs from one rat (means ± SEM). DIC = differential interference contrast.

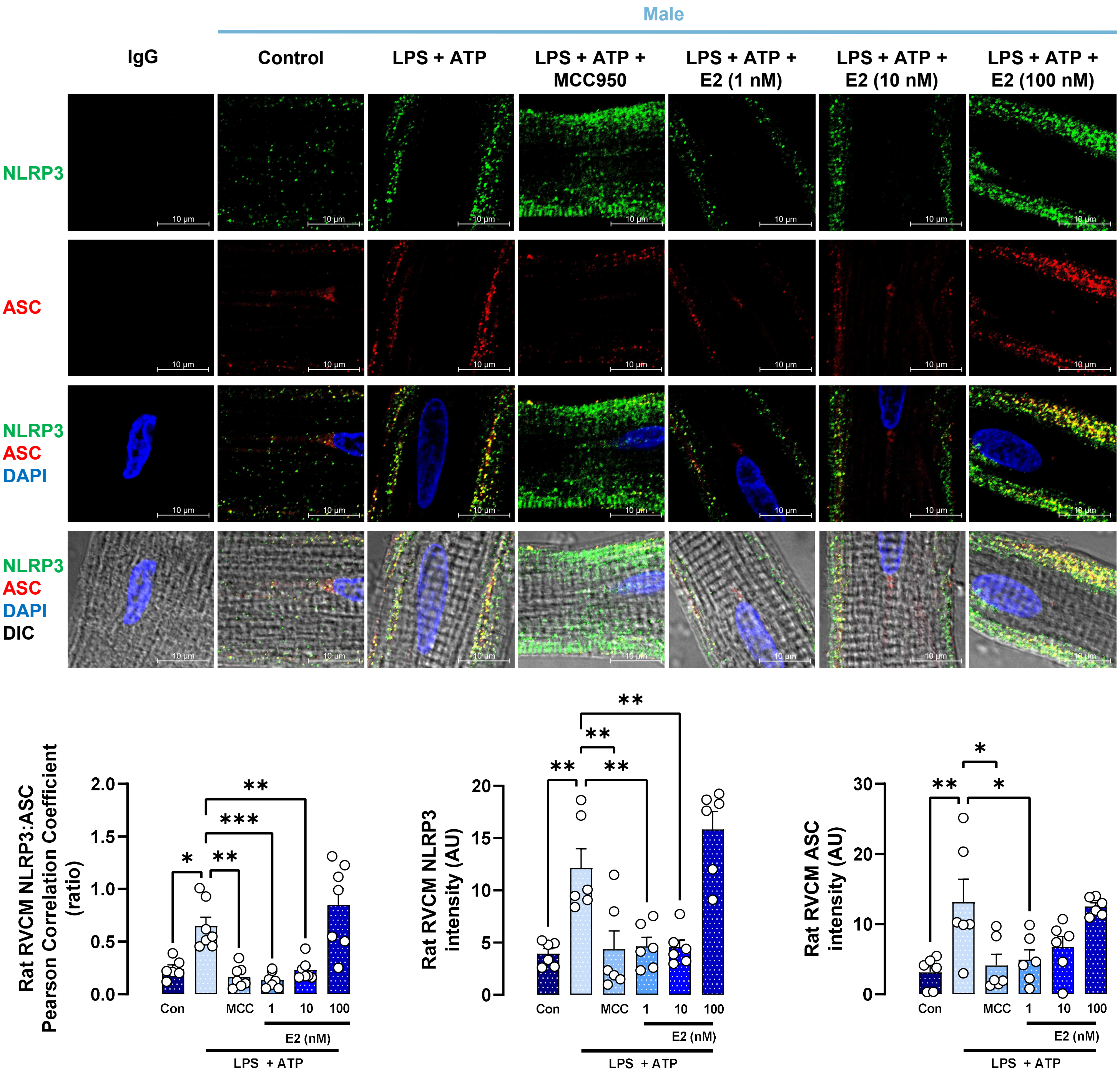

Male
IgG
Control
LPS + ATP
LPS + ATP + MCC950
LPS + ATP + E2 (1 nM)
LPS + ATP + E2 (10 nM)
LPS + ATP + E2 (100 nM)
NLRP3
ASC
NLRP3 ASC DAPI
NLRP3 ASC DAPI DIC
10 µm
Rat RVCM NLRP3:ASC Pearson Correlation Coefficient (ratio)
Rat RVCM NLRP3 intensity (AU)
Rat RVCM ASC intensity (AU)
Con
MCC
1
10
100
E2 (nM)
LPS + ATP

**Fig. E7: Lipopolysaccharide (LPS)- and adenosine triphosphate (ATP)-induced NLRP3 activation in male RVCMs is prevented by E2 in a dose-dependent manner**. Representative immunofluorescence images and quantification of fluorescence intensity of NLRP3 and ASC in RVCMs from male control rats treated in vitro with LPS+ATP in absence or presence of NLRP3 inhibitor MCC950 or E2. RVCMs were treated with LPS (1 μg/mL) for 4 hours and ATP (2 mM) for 10 min prior to cell collection. MCC950 (1 μM) was given 30 min prior to LPS and continued throughout LPS and ATP exposure. E2 (1 nM) was given for 24 hours prior to LPS and continued throughout LPS and ATP exposure. Rat RVCMs were stained for NLRP3 (green), ASC (red), and DAPI (blue). Images were obtained at 100x magnification. Fluorescence intensity was analyzed in ten different RVCMs per animal, and averaged values for each animal are shown. Representative RVCM images were digitally zoomed 5x. *$p<0.05$, **$p<0.01$, ***$p<0.001$ by one-way ANOVA with Holm-Šídák post-test. Each data point = RVCMs from one rat (means ± SEM). DIC = differential interference contrast.

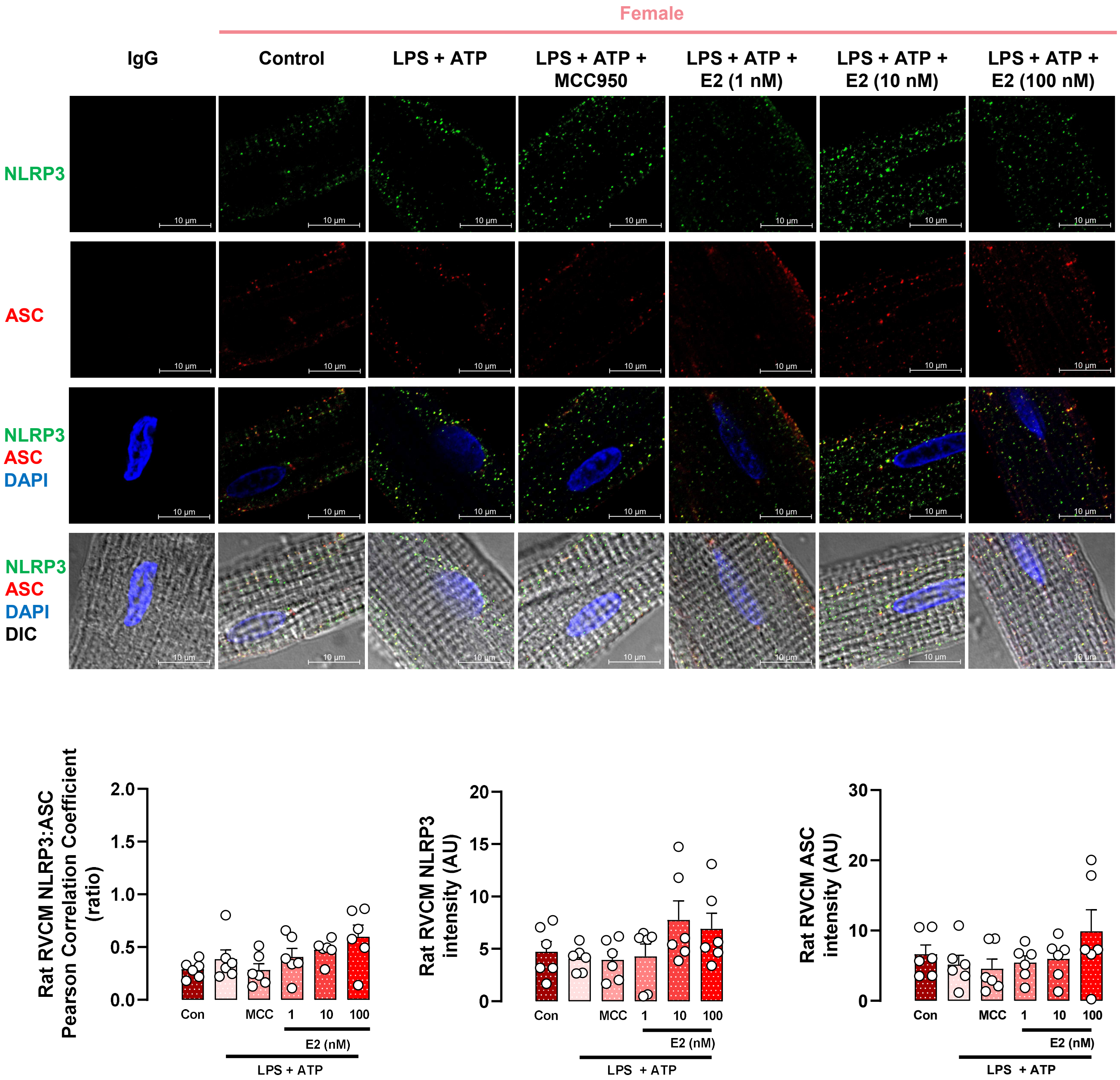
Female
IgG
Control
LPS + ATP
LPS + ATP + MCC950
LPS + ATP + E2 (1 nM)
LPS + ATP + E2 (10 nM)
LPS + ATP + E2 (100 nM)
NLRP3
ASC
NLRP3 ASC DAPI
NLRP3 ASC DAPI DIC
10 µm
Rat RVCM NLRP3:ASC Pearson Correlation Coefficient (ratio)
2.0
1.5
1.0
0.5
0.0
Con
MCC
1
10
100
E2 (nM)
LPS + ATP
Rat RVCM NLRP3 intensity (AU)
20
15
10
5
0
Rat RVCM ASC intensity (AU)
30
20
10
0

**Fig. E8: Female RVCMs are resistant to lipopolysaccharide (LPS)- and adenosine triphosphate (ATP)-induced NLRP3 activation**. Representative immunofluorescence images and quantification of fluorescence intensity of NLRP3 and ASC in RVCMs from female control rats treated in vitro with LPS+ATP in absence or presence of NLRP3 inhibitor MCC950 or E2. RVCMs were treated with LPS (1 μg/mL) for 4 hours and ATP (2 mM) for 10 min prior to cell collection. MCC950 (1 μM) was given 30 min prior to LPS and continued throughout LPS and ATP exposure. E2 (1 nM) was given for 24 hours prior to LPS and continued throughout LPS and ATP exposure. Rat RVCMs were stained for NLRP3 (green), ASC (red), and DAPI (blue). Images were obtained at 100x magnification. Fluorescence intensity was analyzed in ten different RVCMs per animal, and averaged values for each animal are shown. Representative RVCM images were digitally zoomed 5x. Each data point = RVCMs from one rat (means ± SEM). DIC = differential interference contrast.

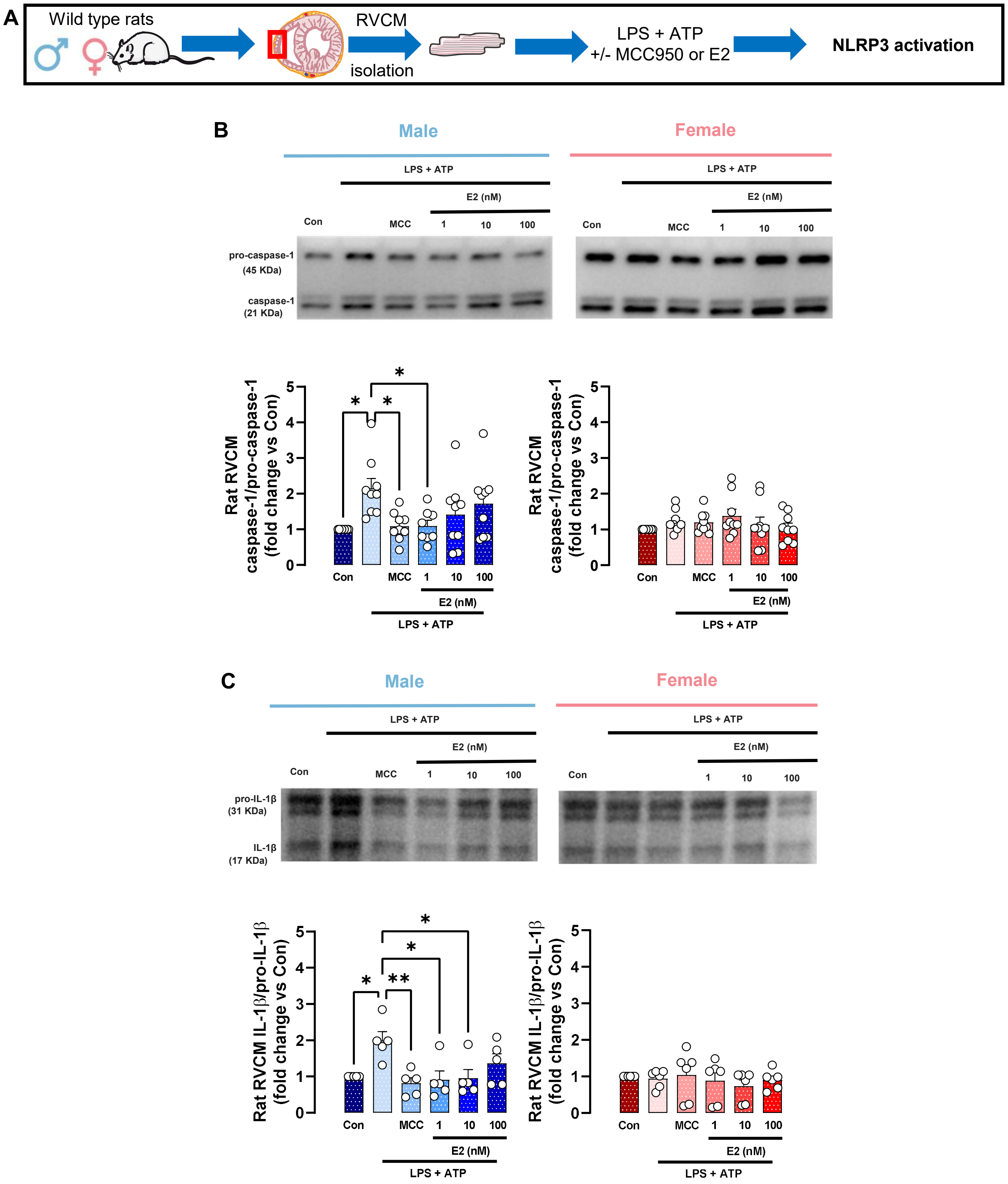
A
Wild type rats
RVCM
isolation
LPS + ATP
+/- MCC950 or E2
NLRP3 activation
B
Male
Female
LPS + ATP
E2 (nM)
Con
MCC
1
10
100
pro-caspase-1
(45 KDa)
caspase-1
(21 KDa)
Rat RVCM
caspase-1/pro-caspase-1
(fold change vs Con)
C
pro-IL-1β
(31 KDa)
IL-1β
(17 KDa)
Rat RVCM IL-1β/pro-IL-1β
(fold change vs Con)

**Fig. E9: Lipopolysaccharide (LPS)- and adenosine triphosphate (ATP)-induced NLRP3 activation in RVCMs is sexually dimorphic and prevented by E2 in male rat RVCMs.** (**A**) Experimental design. (**B, C**) . Representative Western blot images and densitometric quantification in RVCMs from male or female control rats treated in vitro with LPS+ATP in absence or presence of NLRP3 inhibitor MCC950 or E2. RVCMs were treated with LPS (1 µg/mL) for 4 hours and ATP (2 mM) for 10 min prior to cell collection. MCC950 (1 µM) was given 30 min prior to LPS + ATP treatment. E2 (1, 10, or 100 nM) was given for 24 hours prior to LPS + ATP treatment. *$p<0.05$, **$p<0.01$ by ANOVA followed by Holm-Šídák post-test. Each data point = one rat (means ± SEM).

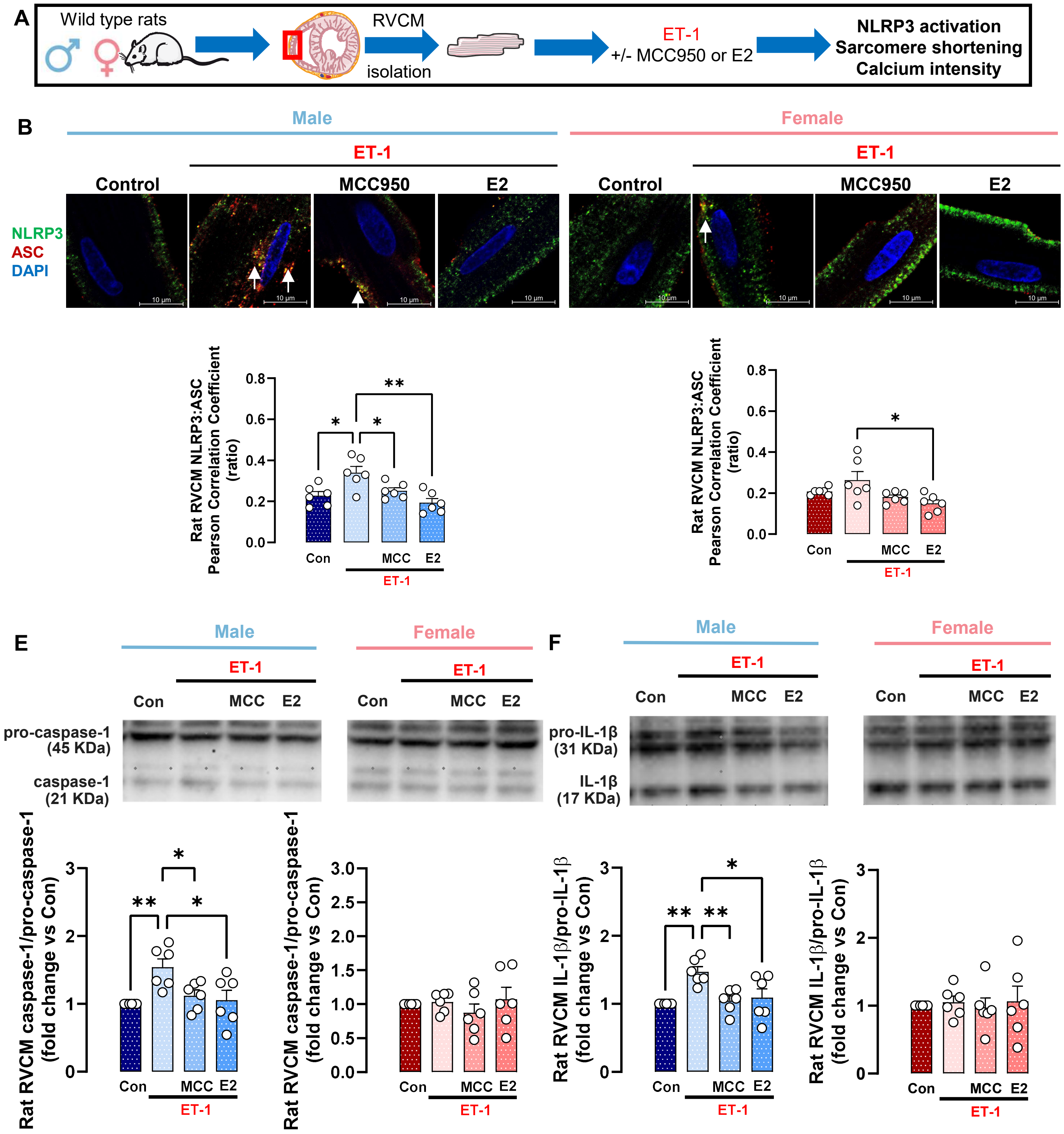

A
Wild type rats
RVCM
isolation
ET-1
+/- MCC950 or E2
NLRP3 activation
Sarcomere shortening
Calcium intensity
B
Male
Female
ET-1
Control
MCC950
E2
NLRP3
ASC
DAPI
10 µm
Rat RVCM NLRP3:ASC Pearson Correlation Coefficient (ratio)
Con
MCC
E2
E
F
pro-caspase-1 (45 KDa)
caspase-1 (21 KDa)
pro-IL-1β (31 KDa)
IL-1β (17 KDa)
Rat RVCM caspase-1/pro-caspase-1 (fold change vs Con)
Rat RVCM IL-1β/pro-IL-1β (fold change vs Con)

**Fig. E10: Endothelin 1-induced NLRP3 activation in rat RVCMs is sexually dimorphic and prevented by E2 in male rat RVCMs.** (**A**) Experimental design. (**B**) Representative immunofluorescence images and quantification of fluorescence intensity of NLRP3-ASC co-localization in RVCMs from male or female control rats treated in vitro with endothelin-1 (ET-1; 1 nM) given 4 hours prior to cell collection +/- MCC950 (1 μM; administered 30 min prior to ET-1 and continued throughout ET-1 exposure) or E2 (1 nM, administered 24 hours prior to ET-1 and continued throughout ET-1 exposure). Rat RVCMs were stained for NLRP3 (green), ASC (red), and DAPI (blue). Images were obtained at 100x magnification. Co-localization images were analyzed in ten different RVCMs per animal, and averaged values for each animal are shown. NLRP3 and ASC co-localization (yellow; indicated by arrows) was quantified by determining Pearson correlation coefficient. Representative RVCM images were digitally zoomed 5x. (**C, D**) Representative Western blot images and densitometric quantification of caspase-1/pro-caspase-1 ratio (**C**) and IL-1β/pro-IL-1β ratio (**D**) in RVCMs from male or female control rats treated in vitro with ET-1 in absence or presence of MCC950 or E2 as outlined above. *$p<0.05$, **$p<0.01$ by one-way ANOVA with Holm-Šídák post-test. Each data point = RVCMs from one rat (means ± SEM).

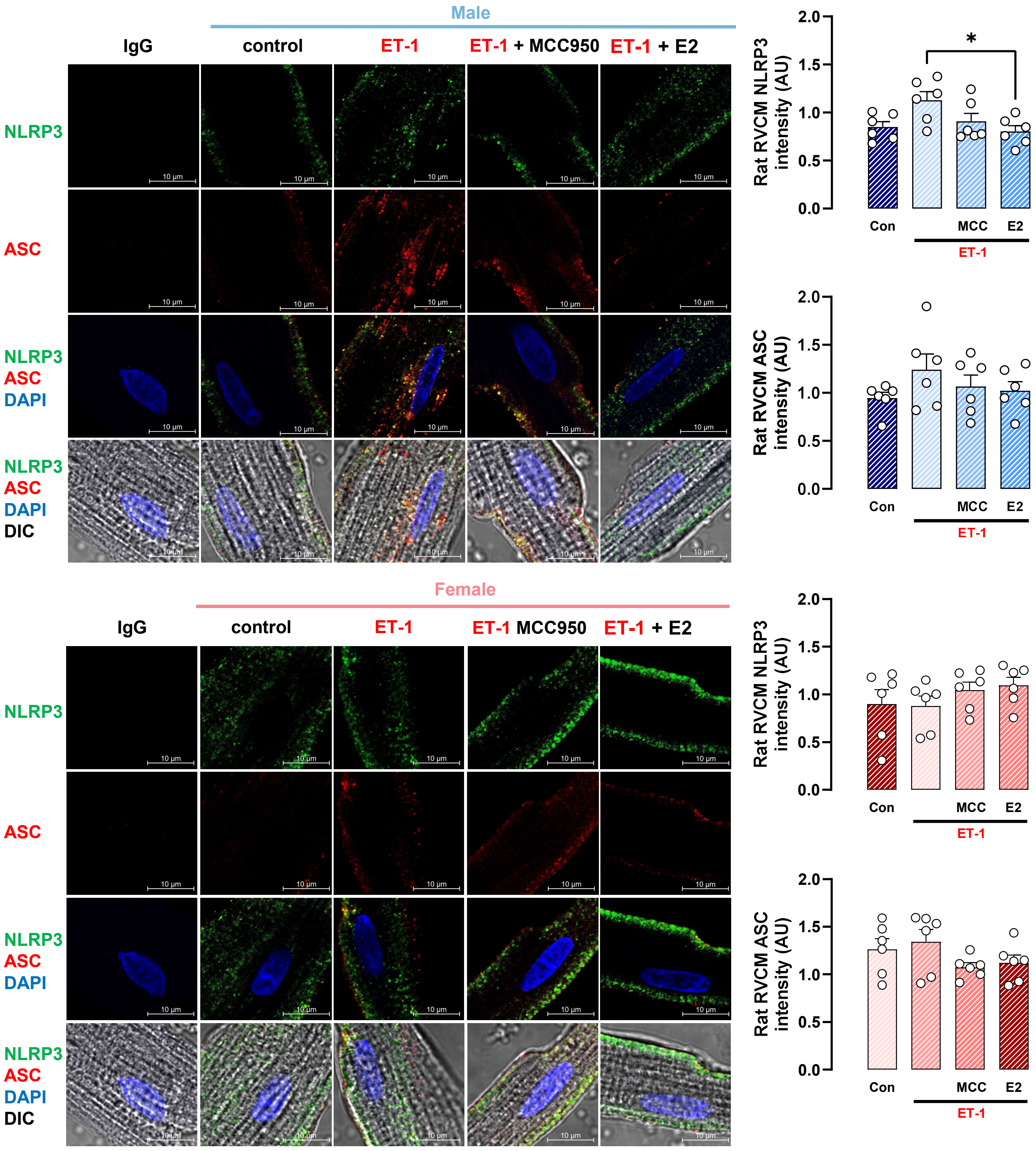
Male
IgG
control
ET-1
ET-1 + MCC950
ET-1 + E2
NLRP3
ASC
NLRP3
ASC
DAPI
NLRP3
ASC
DAPI
DIC
10 µm
Rat RVCM NLRP3 intensity (AU)
*
Con
MCC
E2
ET-1
Rat RVCM ASC intensity (AU)
Con
MCC
E2
ET-1
Female
IgG
control
ET-1
ET-1 MCC950
ET-1 + E2
NLRP3
ASC
NLRP3
ASC
DAPI
NLRP3
ASC
DAPI
DIC
Rat RVCM NLRP3 intensity (AU)
Con
MCC
E2
ET-1
Rat RVCM ASC intensity (AU)
Con
MCC
E2
ET-1

**Fig. E11: Endothelin-1 effects on NLRP3 and ASC expression in rat RVCMs.** Representative immunofluorescence images and quantification of fluorescence intensity of NLRP3 and ASC in RVCMs from male or female control rats treated in vitro with endothelin-1 (ET-1; 1 nM) given 4 hours prior to cell collection +/- MCC950 (1 μM; administered 30 min prior to ET-1 and continued throughout ET-1 exposure) or E2 (1 nM, administered 24 hours prior to ET-1 and continued throughout ET-1 exposure). Rat RVCMs were stained for NLRP3 (green), ASC (red), and DAPI (blue). Images were obtained at 100x magnification. Fluorescence intensity was analyzed in ten different RVCMs per animal, and averaged values for each animal are shown. Representative RVCM images were digitally zoomed 5x. *$p<0.05$ by one-way ANOVA with Holm-Šídák post-test. Each data point = RVCMs from one rat (means ± SEM). DIC = differential interference contrast.

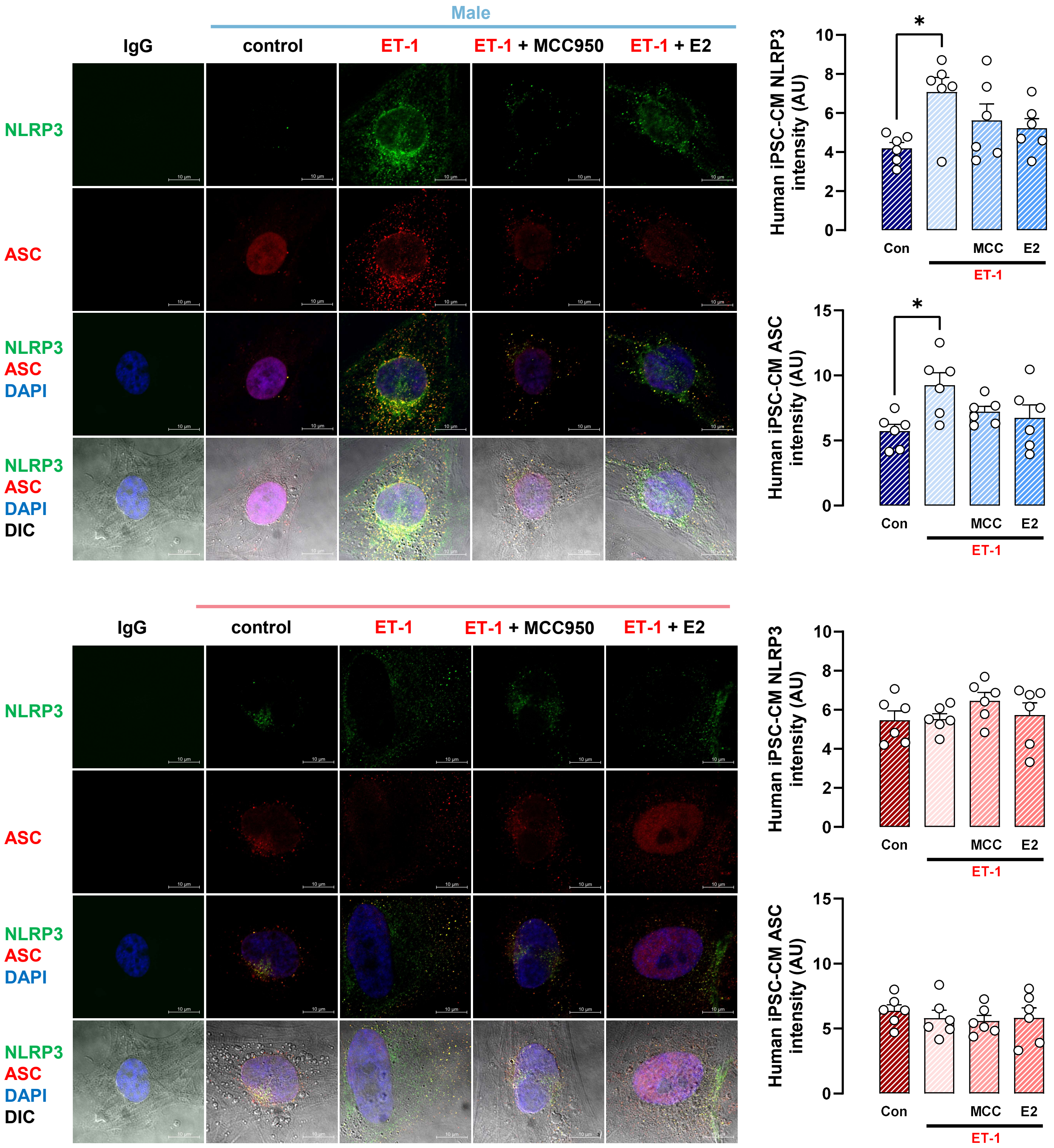
Male
IgG
control
ET-1
ET-1 + MCC950
ET-1 + E2
NLRP3
ASC
NLRP3
ASC
DAPI
NLRP3
ASC
DAPI
DIC
Human iPSC-CM NLRP3 intensity (AU)
Human iPSC-CM ASC intensity (AU)
Con
MCC
E2
ET-1
*
IgG
control
ET-1
ET-1 + MCC950
ET-1 + E2
NLRP3
ASC
NLRP3
ASC
DAPI
NLRP3
ASC
DAPI
DIC
Human iPSC-CM NLRP3 intensity (AU)
Human iPSC-CM ASC intensity (AU)
Con
MCC
E2
ET-1

**Fig. E12: Endothelin 1-induced NLRP3 and ASC expression is sexually dimorphic, male biased, and prevented by E2 in human induced pluripotent stem cell-derived cardiomyocytes (iPSC-CMs)**. Representative immunofluorescence images and quantification of fluorescence intensity of NLRP3 and ASC in human iPSC-CMs treated with endothelin-1 (ET-1; 1 nM) given 4 hours prior to cell collection +/- MCC950 (1 µM; administered 30 min prior to ET-1 and continued throughout ET-1 exposure) or E2 (1 nM, administered 24 hours prior to ET-1 and continued throughout ET-1 exposure). Cells were stained for NLRP3 (green), ASC (red), and DAPI (blue). Fluorecence intensity was quantified in ten different human iPSC-CMs per patient and values were then averaged. Each data point = one human iPSC-CM clone (means ± SEM). *$p<0.05$ by one-way ANOVA with Holm-Šídák post-test. DIC = differential interference contrast.

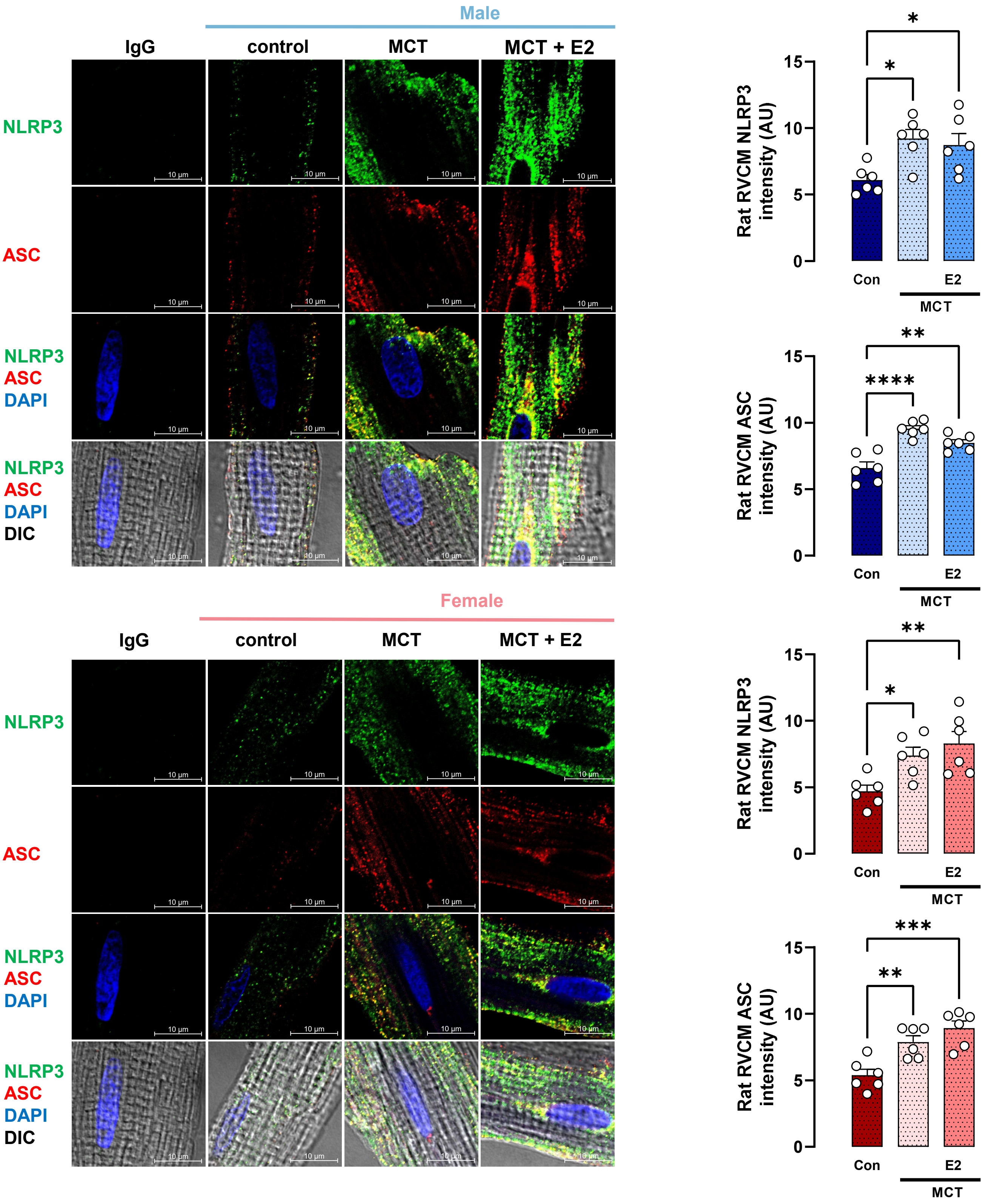

Male
IgG
control
MCT
MCT + E2
NLRP3
ASC
NLRP3
ASC
DAPI
NLRP3
ASC
DAPI
DIC
10 µm
Female
IgG
control
MCT
MCT + E2
NLRP3
ASC
NLRP3
ASC
DAPI
NLRP3
ASC
DAPI
DIC
Rat RVCM NLRP3 intensity (AU)
Rat RVCM ASC intensity (AU)
Rat RVCM NLRP3 intensity (AU)
Rat RVCM ASC intensity (AU)
0
5
10
15
Con
E2
MCT
*
**
***
****

**Fig. E13: Estrogen receptor α (ERα) is necessary for E2 to attenuate NLRP3 and ASC expression in male and female rat MCT-RVCMs**. Representative immunofluorescent images, quantification of fluorescence intensity of NLRP3, and ASC fluorescence intensity in male ERα loss of function ($ER\alpha^{mut}$) rat right ventricular cardiomyocytes (RVCM) from control or MCT males treated with E2 (75 µg/kg/day, s.c. pellets). The images were obtained using Confocal Zeis LSM 700 microscope and Zen Black 3.5 software at 100x magnification. $ER\alpha^{mut}$ rat RVCMs were stained for NLRP3 (green), ASC (red), and DAPI (blue). Fluorescence intensity images were analyzed in 10 different RVCM. Representative RV images were digitally zoomed 5x. *$p<0.05$, **$p<0.01$, ****$p<0.0001$ by one-way ANOVA followed by Holm-Šídák post-test (means ± SEM). DIC = differential interference contrast.

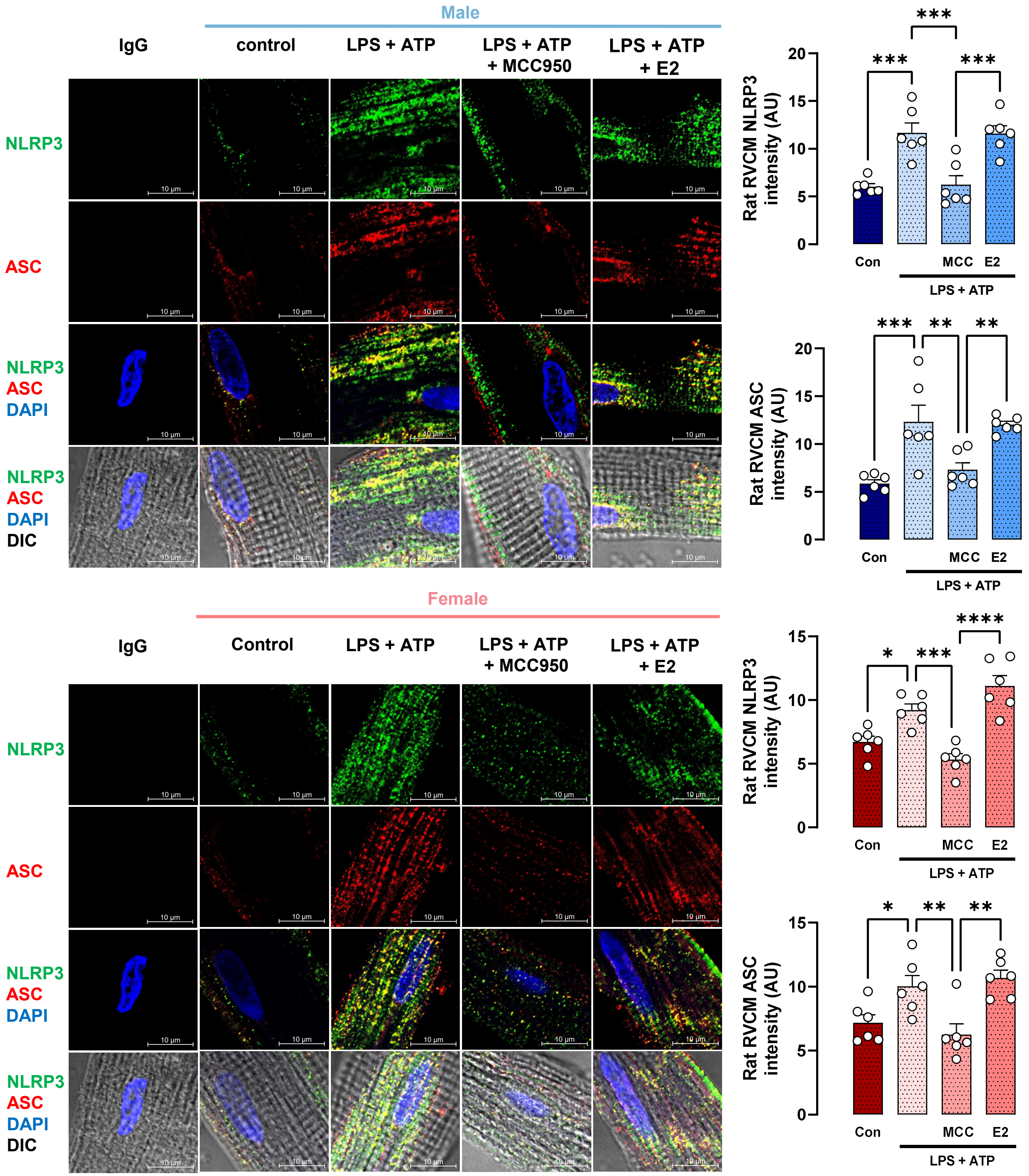

Male
IgG
control
LPS + ATP
LPS + ATP + MCC950
LPS + ATP + E2
NLRP3
ASC
NLRP3 ASC DAPI
NLRP3 ASC DAPI DIC
10 μm
Rat RVCM NLRP3 intensity (AU)
Rat RVCM ASC intensity (AU)
Con
MCC
E2
LPS + ATP
***
**
Female
IgG
Control
LPS + ATP
LPS + ATP + MCC950
LPS + ATP + E2
*
****

**Fig. E14: Lack of functioning estrogen receptor α (ERα) eliminates sexual dimorphisms in LPS + ATP-induced increases in NLRP3 and ASC expression in rat RVCMs and prevents E2 from preventing increases in NLRP3 and ASC abundance**. Representative immunofluorescence images and quantification of NLRP3 and ASC fluorescence intensity in RVCMs from male or female ERα loss of function mutant ($ER\alpha^{mut}$) rats after treatment with LPS and ATP in vitro. LPS (1μg/mL) was given 4 hours prior to cell collection. ATP (2 mM) was given 10 min prior to cell collection. MCC950 (1μM) or E2 (1 nM) were given 30 min or 24 hours prior LPS and ATP, respectively, and continued throughout the exposure. RVCMs were stained for NLRP3 (green), ASC (red), and DAPI (blue). Images were obtained at 100x magnification. Fluorescence intensity was analyzed in ten different RVCMs per animal, and averaged values for each animal are shown. Representative images were digitally zoomed 5x. *p<0.05, **p<0.01, ***p<0.001, ****p<0.0001 by one-way ANOVA with Holm-Šídák post-test. Each data point = RVCMs from one rat (means ± SEM). DIC = differential interference contrast.

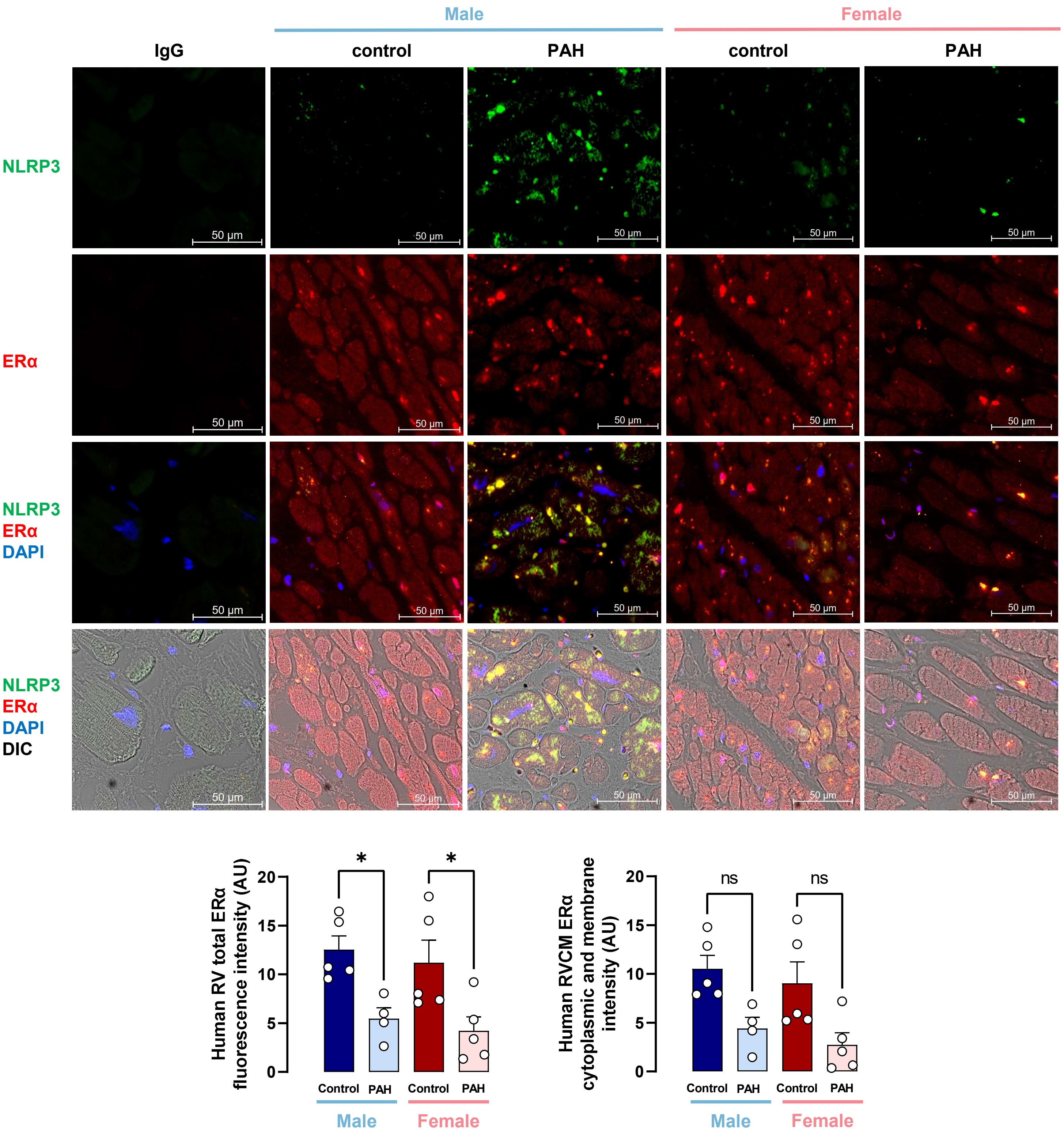


**Fig. E15: Human pulmonary artery hypertension patients with RV failure exhibit reduced nuclear estrogen receptor α (ERα) abundance in RVCMs.** Representative immunofluorescent images, total ERα fluorescence intensity, and cytoplasmic and membrane ERα fluorescence intensity in RVs from male or female PAH patients with RV failure were analyzed in whole RV slide under 20x magnification. Representative images are 40x. *$p<0.05$, ns = not significant by two-way ANOVA followed by Holm-Šídák post-test. Each data point represents one RV sample from one unique patient (means ± SEM). DIC = differential interference contrast.

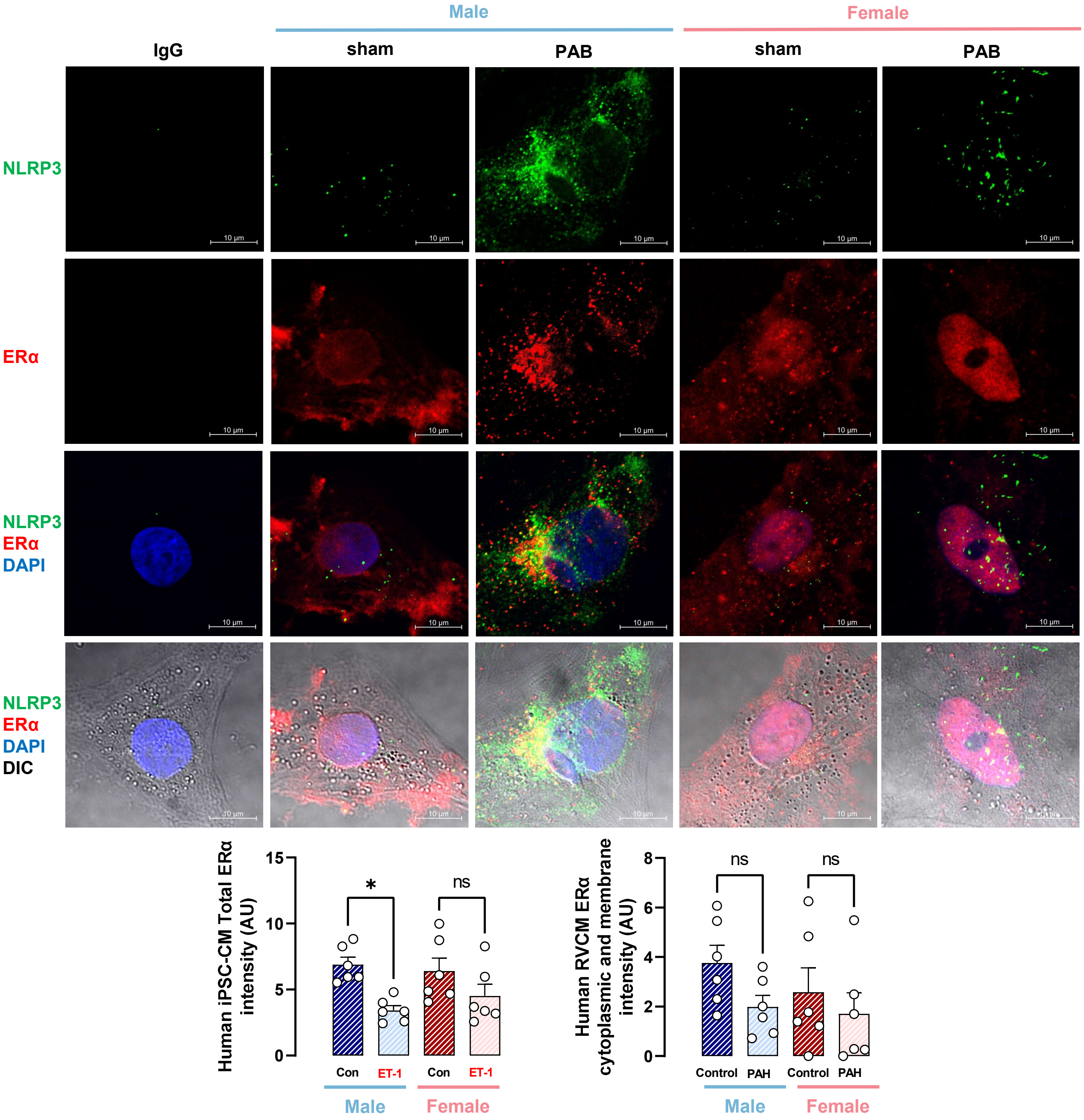
Male
Female
IgG
sham
PAB
sham
PAB
NLRP3
ERα
NLRP3
ERα
DAPI
NLRP3
ERα
DAPI
DIC
10 µm
Human iPSC-CM Total ERα intensity (AU)
15
10
5
0
*
ns
Con
ET-1
Con
ET-1
Male
Female
Human RVCM ERα cytoplasmic and membrane intensity (AU)
8
6
4
2
0
ns
ns
Control
PAH
Control
PAH
Male
Female

**Fig. E16: Endothelin 1-induced reduction in total ERα abundance in human induced pluripotent stem cell-derived cardiomyocytes (hiPSC-CMs) is sexually dimorphic**. Representative immunofluorescent images, total ERα fluorescence intensity, and cytoplasmic and membrane fluorescence intensity in hiPSC-CMs treated with endothelin-1 (ET-1; 1 nM) given 4 hours prior to cell collection +/- MCC950 (1 μM; administered 30 min prior to ET-1 and continued throughout ET-1 exposure) or E2 (1 nM, administered 24 hours prior to ET-1 and continued throughout ET-1 exposure). hiPSC-CMs were stained for NLRP3 (green), ERα (red) and DAPI (blue). Fluorescence intensity was quantified in ten different hiPSC-CMs per patient and values were then averaged. Each data point = one human iPSC-CM clone (means ± SEM). *$p<0.05$, ns = not significant by one-way ANOVA with Holm-Šídák post-test. DIC = differential interference contrast.

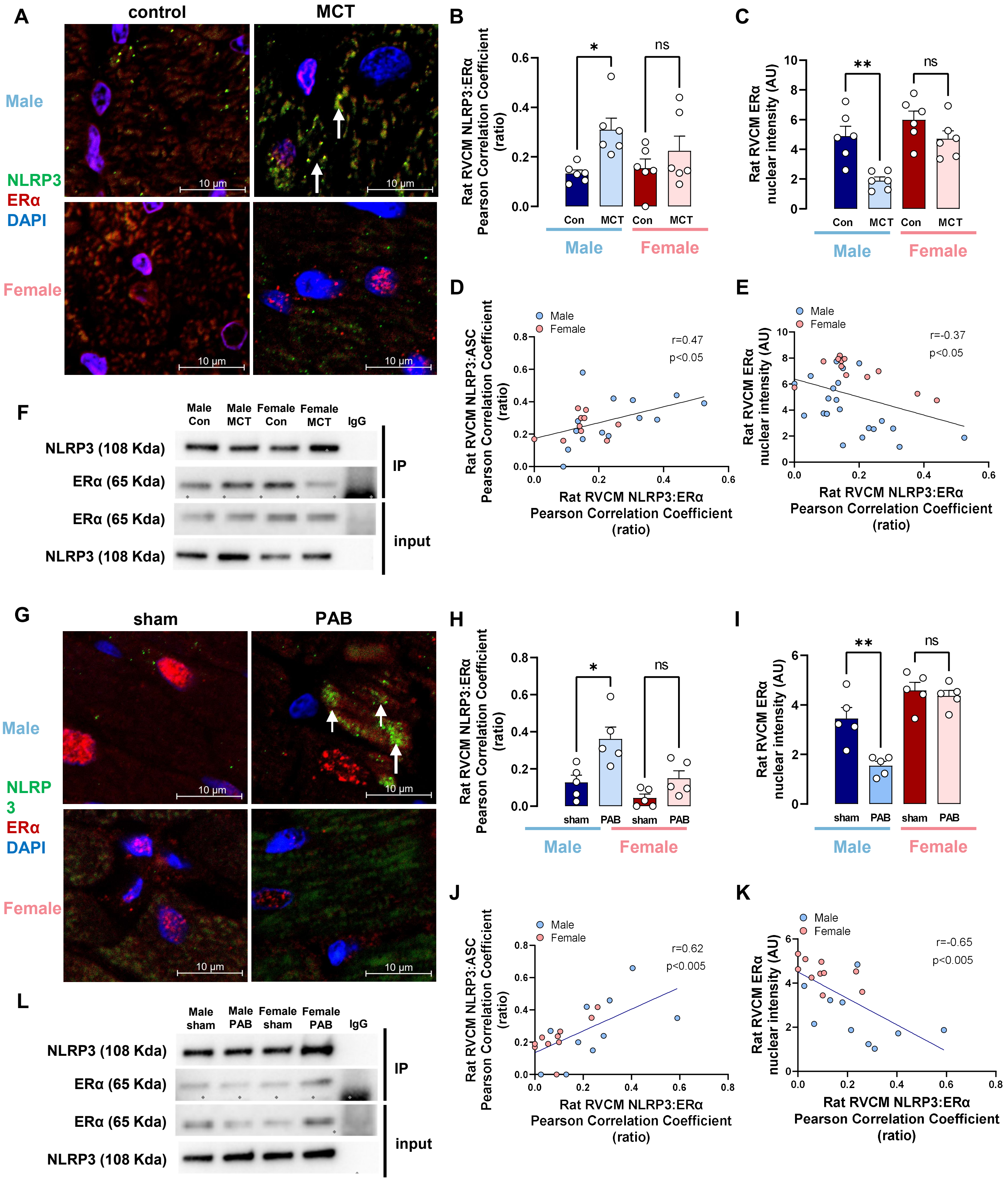
A
control
MCT
Male
NLRP3
ERα
DAPI
Female
10 μm
B
Rat RVCM NLRP3:ERα Pearson Correlation Coefficient (ratio)
*
ns
Con
MCT
Male
Female
C
Rat RVCM ERα nuclear intensity (AU)
**
ns
D
Male
Female
r=0.47
p<0.05
Rat RVCM NLRP3:ASC Pearson Correlation Coefficient (ratio)
Rat RVCM NLRP3:ERα Pearson Correlation Coefficient (ratio)
E
r=-0.37
p<0.05
F
Male Con
Male MCT
Female Con
Female MCT
IgG
NLRP3 (108 Kda)
ERα (65 Kda)
IP
input
G
sham
PAB
NLRP
3
H
I
J
r=0.62
p<0.005
K
r=-0.65
p<0.005
L
Male sham
Male PAB
Female sham
Female PAB

**Fig. E17: Male rats exhibit more pronounced RVCM NLRP3 and ERα interaction than female rats in the setting of RV pressure overload.** (**A, B**) ERα-NLRP3 co-localization in RVs from MCT-PH rats. Representative images (**A**) and quantification (**B**) of co-localization of NLRP3 (green) with ERα (red). Magnification is 20x. Co-localization was analyzed in the whole RV sample; representative images are 40x. Arrows indicate ERα-NLRP3 co-localization (yellow). Co-localization was quantified by determining Pearson correlation coefficient between NLRP3 and ERα. Note sexual dimorphism and male bias with more pronounced co-localization in male MCT-PH RVs. (**C**) depicts ERα nuclear intensity. (**D, E**) demonstrate correlations between ERα-NLRP3 co-localization intensity and NLRP3 activation (NLRP3-ASC co-localization; **D**) and nuclear ERα abundance (**E**). Note decrease in nuclear ERα abundance with increase in ERα-NLRP3 co-localization. (**F**) Representative images for the Co-Immunoprecipitation of NLRP3 and ERα in RVs from MCT-PH rats. (**G, H**) ERα-NLRP3 co-localization in RVs from PAB rats. Representative images (**G**) and quantification (**H**) of co-localization of NLRP3 (green) with ERα (red). Magnification is 20x. Co-localization was analyzed in whole RV sample; representative images are 40x. Arrows indicate ERα-NLRP3 co-localization (yellow). Co-localization was quantified by determining Pearson correlation coefficient between NLRP3 and ERα. Note sexual dimorphism and male bias with more pronounced co-localization in male PAB RVs. (**I**) depicts ERα nuclear intensity. (**J, K**) demonstrate correlations between ERα-NLRP3 co-localization intensity and NLRP3 activation (NLRP3-ASC co-localization; **J**) and nuclear ERα abundance (**K**). Note decrease in nuclear ERα abundance with increase in ERα-NLRP3 co-localization. (**I**) Representative images for co-immunoprecipitation of NLRP3 and ERα in RVs from PAB rats. *$p<0.05$, **$p<0.01$, ns = not significant by two-way ANOVA with Holm-Šídák post-test. Each data point = one rat (means ± SEM).

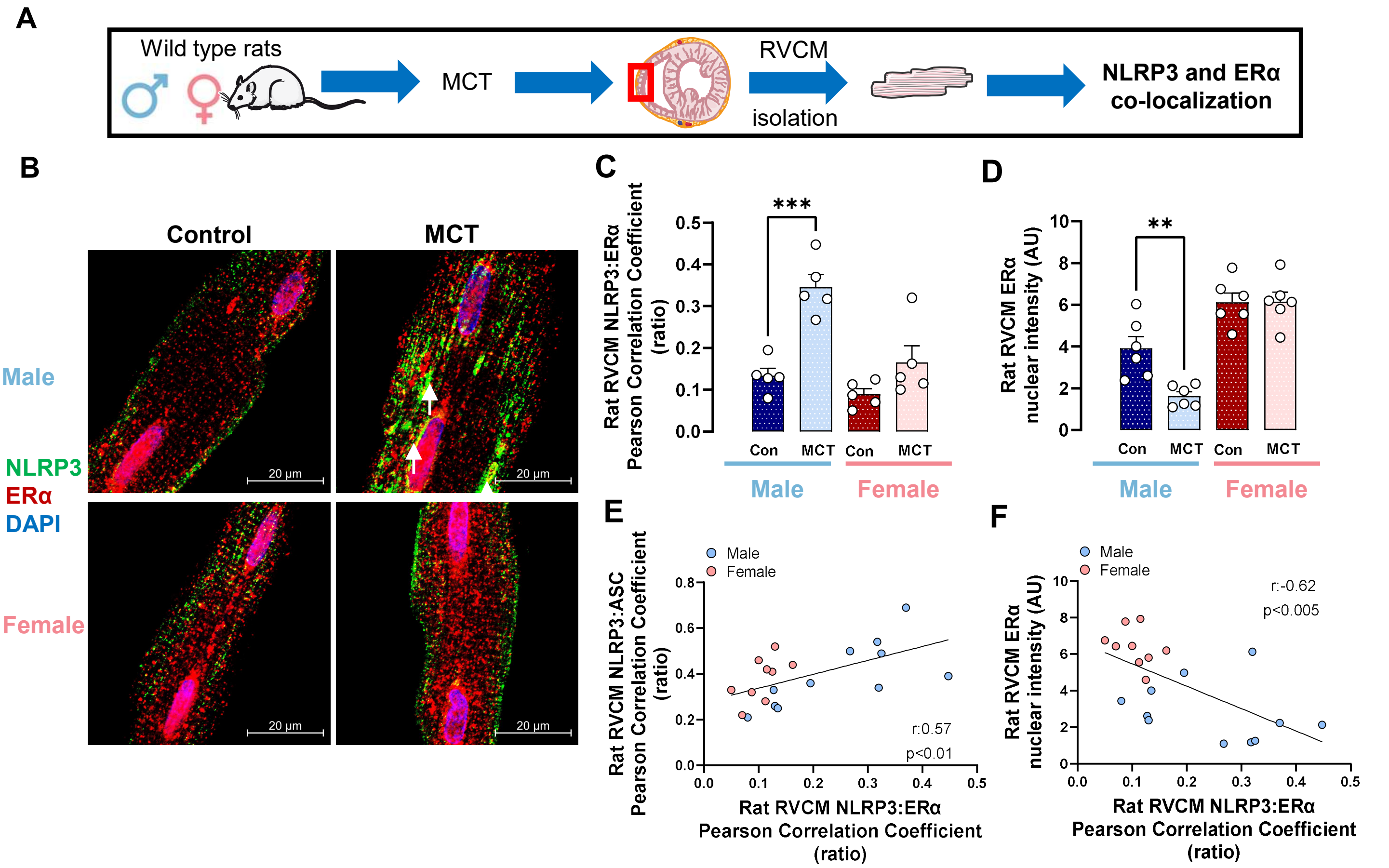


**Fig. E18: ERα colocalizes with NLRP3 in RVCMs from rats with MCT-induced RV failure.** (**A**) Experimental design. (**B, C**) ERα-NLRP3 co-localization in RVCMs from male or female control or MCT-PH rats. Representative images (**B**) and quantification (**C**) of co-localization of NLRP3 (green) with ERα (red). Magnification is 100x. Co-localization (arrows) was quantified by determining Pearson correlation coefficient between NLRP3 and ERα. Note sexual dimorphism and male bias with more pronounced co-localization in male MCT-PH RVCMs. (**D**) depicts ERα nuclear intensity. (**E, F**) demonstrate correlations between ERα-NLRP3 co-localization intensity and NLRP3 activation (NLRP3-ASC co-localization; **E**) and nuclear ERα abundance (**F**). Decrease in nuclear ERα abundance with increase in ERα-NLRP3 co-localization is again noted. **$p<0.01$, ***$p<0.01$ by two-way ANOVA with Holm-Šídák post-test. Each data point = RVCMs from one rat (means ± SEM).
